# Stack: In-Context Learning of Single-Cell Biology

**DOI:** 10.64898/2026.01.09.698608

**Authors:** Mingze Dong, Abhinav Adduri, Dhruv Gautam, Lujing Wu, Courtney Kernick, Mary Margaret Coons, Yi-Chen Chih, Christopher Carpenter, Rohan Shah, Chiara Ricci-Tam, Po-Yuan Tung, Nianzhen Li, Alexander Dobin, Yuval Kluger, Dave P. Burke, Theodore L. Roth, Yusuf H. Roohani

## Abstract

Foundation models trained on single-cell transcriptomic data offer the promise of identifying and predicting the diversity of cellular phenotypes across species, diseases, and other biological conditions. However, the current models are limited to their supervised training conditions and tasks, which limits their utility for biological discovery. Here, we present Stack, a foundation model trained on 149 million uniformly preprocessed human single cells that leverages tabular attention to generate representations for each cell informed by the cells in its context. Stack offers substantial improvements for downstream tasks in the zero-shot setting compared to baselines, whether they are zero-shot, fine-tuned, or trained from scratch on the target dataset. Stack can perform in-context learning from unlabeled cells representing arbitrary conditions, such as a chemical perturbation or a different donor, and predict the effect of those conditions on a target cell population without requiring data-specific fine-tuning. We apply Stack to generate *Perturb Sapiens*, the first human whole-organism atlas of perturbed cells, spanning 28 tissues, 40 cell types, and 892 drug, cytokine, and genetic perturbations. We validated subsets of *Perturb Sapiens* using *in vitro* stimulation profiles. Stack uniquely empowers prioritization of donor-specific perturbation effects, a capability we validated in our newly collected DiseasePert-3M data, comprising T cells from 40 donors across 14 diseases, stimulated with 11 cytokines. Overall, Stack presents a new modeling framework where cells themselves act as guiding examples at inference time, unlocking general-purpose in-context learning capabilities for single-cell biology.

## 1. Introduction

Despite the promise of current single-cell foundation models to leverage the unprecedented volumes of large-scale single-cell RNA sequencing data, they face significant limitations. They often fail to surpass classical approaches when employed in a zero-shot manner (Kedzierska et al., 2025; Liu et al., 2023). Even with dataset-specific fine-tuning, they struggle to improve over simple baselines in perturbation prediction (Wu et al., 2024; Li et al., 2024; Kernfeld et al., 2023; Ahlmann-Eltze et al., 2025). Recent models (such as Adduri et al., 2025 and Klein et al., 2025) show improvements in these capabilities; however, they still require extensive training data and supervision on biological conditions and tasks of interest. This limits their potential for *de novo* biological discoveries, such as inferring perturbation effects in novel biological contexts (e.g. unseen cell types for which only observational data is available) or performing novel tasks, such as predicting donor-specific variation in immune phenotypes.

Most current single-cell foundation models are constrained by fundamental design choices. First, their pre-training objectives operate at the single-cell level, training models to function as universal “denoisers” that exploit gene dependencies but cannot see shifts at the population scale. Second, the inherently noisy count distribution of gene expression profiles necessitates aggregating information across cells to enhance signal-to-noise ratios of gene expression patterns. This insight has been employed in State for perturbation effect prediction (Adduri et al., 2025), but remains underexplored for single-cell foundation models. This aggregation is also essential for encoding inter-cellular interactions or mutual information that would otherwise be neglected or misattributed to gene-level dependencies. Third, prior models have largely been used for downstream tasks through task-specific fine-tuning on limited data, rather than exploiting the demonstrated ability of large transformer based models to perform robust learning at inference time.

To address these limitations, we developed Stack, a self-supervised framework pre-trained on scBaseCount (Youngblut et al., 2025), the largest existing single-cell collection with 189 million high-quality human cells. Stack employs a tabular attention architecture over cell sets, enabling both intra- and inter-cellular information flow to produce context-aware representations. Through extensive evaluations, we show that Stack’s zero-shot embeddings consistently outperform not only existing single-cell foundation models, but also models trained from scratch or fine-tuned on each evaluation dataset, an outcome that, to our knowledge, has not been demonstrated by any existing single-cell foundation model.

Through a novel post-training procedure, Stack enables in-context learning (ICL) for single-cell biology: given prompt cells representing an arbitrary condition and query cells of interest, Stack predicts counterfactual query cell states, i.e. the gene expression state of query cells under the biological condition of the prompt. This formulation enables generalization to cell types never experimentally perturbed, across perturbation categories, and to tasks beyond those seen during training. Stack ranks first in 28 of 31 evaluations spanning perturbational, observational, and hybrid prompting tasks on entirely unseen datasets and conditions (Kernfeld et al., 2023; Adduri et al., 2025; Luecken et al., 2022). We leveraged Stack to construct *Perturb Sapiens*, the first human whole-organism perturbational atlas, spanning 28 tissues, 40 cell types, and 892 perturbations with multiple donor replicates, whose perturbation-, cell-type-, tissue-, and donor-specific effects we validated against independent *in vitro* stimulation datasets. Stack further enables identification of cytokines and disease donors associated with donor-specific perturbation responses, which we validated using our newly collected dataset, DiseasePert-3M, comprising 40 donors stimulated by 11 cytokines across 14 disease conditions, the largest existing single-cell perturbation dataset in donor and disease heterogeneity.

## 2. Results

### **2.1.** Stack leverages cellular context to learn cell representations that generalize across datasets and tasks

Stack is a large-scale self-supervised foundation model designed to learn fundamental dependencies across cells and genes from large single-cell data collections. We pre-trained Stack on human cells from the scBaseCount (Youngblut et al., 2025), the largest available human single-cell data collection that contains 189 million cells after strict quality control (Methods). The training dataset comprised 19,978 SRX samples and 148.8 million cells, with the remaining 20% of the data held out for validation and testing (Fig. 1A, Methods). Our high-quality training set is over four-fold larger than those of scGPT and Geneformer, and more than three-fold larger than that of UCE (Cui et al., 2024; Theodoris et al., 2023; Rosen et al., 2023). To accelerate model training, we developed a highly efficient h5py-based dataloader that reads consecutive chunks for each single-cell data sample and caches cell index sets (Fig. 1A, Methods). The dataloader achieves high input-pipeline throughput (around 1.6 × 10^4^ cells/second, over 75× faster than a similar model (Adduri et al., 2025)), sufficient to saturate GPU compute and complete pre-training in 2–3 days on a single H100 GPU.

**Figure 1.**
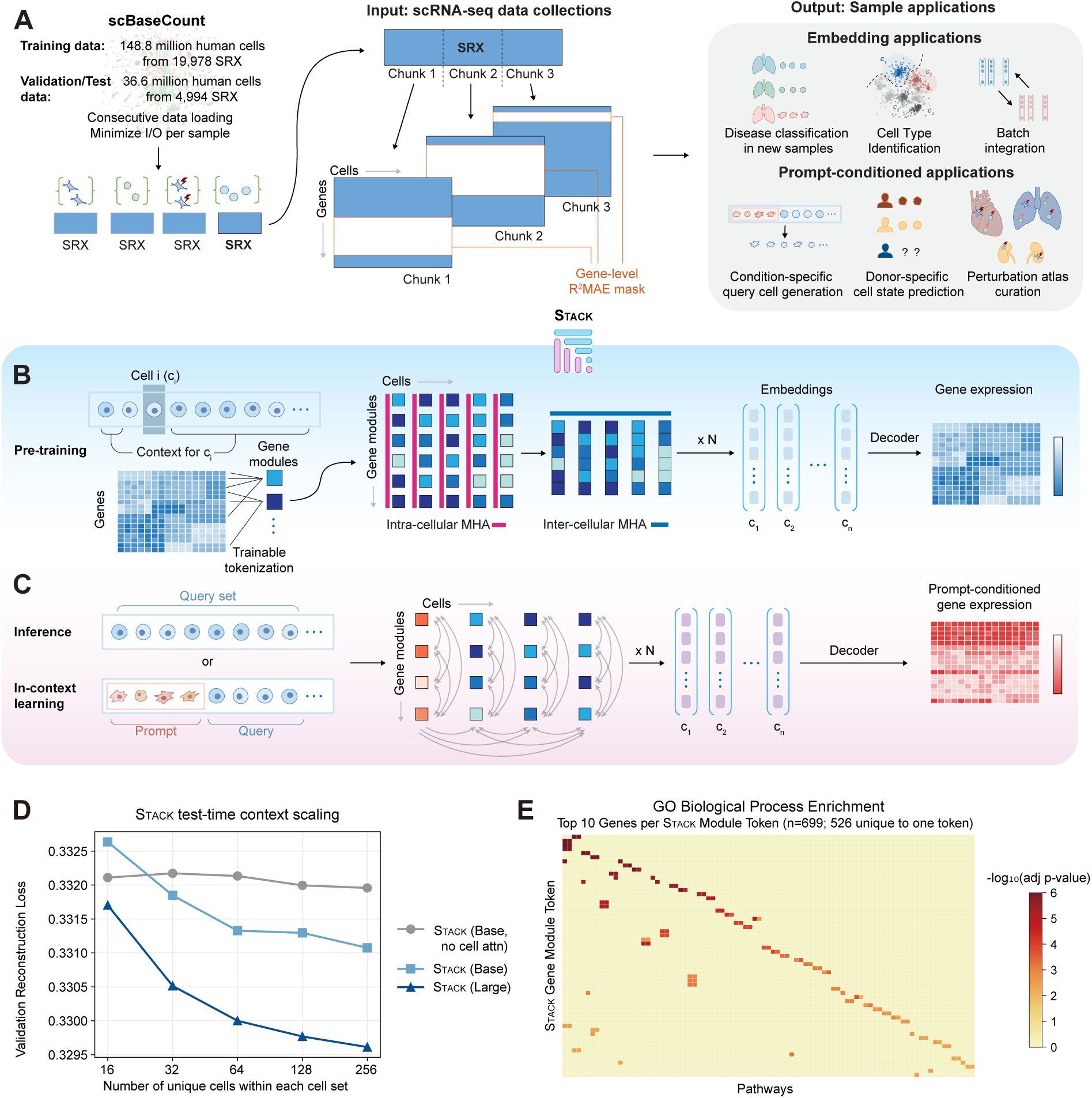
Stack: A single-cell foundation model that leverages cellular context. **A.** Overview of the Stack model. As input, Stack takes human single-cell data from scBaseCount comprising 148.8 million cells from 19,978 SRX files after preprocessing and filtering (Youngblut et al., 2025). Each file is chunked into consecutive cell sets of fixed size as model input. During pre-training, each input cell set is corrupted by masking a random subset of genes across all cells with a variable masking ratio (Dong et al., 2025). After pre-training, Stack enables zero-shot embedding analysis and a wide range of downstream tasks guided by designed prompts. **B.** Pre-training of Stack. Stack employs a single-layer perceptron to project cells into a set of gene module tokens. The tokens are passed through a tabular attention architecture that iteratively applies multi-head attention (MHA) along both gene and cell dimensions, followed by a feedforward network. After *𝑁* tabular attention blocks, the final tokens are concatenated and flattened into one-dimensional vectors per cell (embeddings), and a decoder projects the embeddings back to expression space (Methods). **C.** Inference-time learning of Stack. The model takes either a single query cell set or a concatenation of prompt and query cell sets as input. Stack performs in-context learning through the tabular attention architecture, and can output prompt-conditioned gene expression. **D.** Effect of unique cell numbers on validation reconstruction loss under different Stack settings. All Stack models presented here were trained on the scBaseCount subset using identical training and validation sets (Methods). **E.** Heatmap of adjusted *𝑝*-values from Gene Ontology (GO) biological process gene set enrichment analysis, showing the top 10 highest-importance genes (ranked by tokenization weight magnitude) for each Stack (Large) token after pre-training on full human scBaseCount. Module and pathway names, along with additional plot details, are provided in Fig. S2.

The input to Stack is a collection of cells, or a *cell set*, from a single experimental sample. We define each cell’s *context* as the remaining cells in its set. Stack makes use of a rectangular mask pre-training task that prevents simple imputation shortcuts and enforces single-cell level resolution. Within a mini-batch, a randomly sampled list of genes is masked for all cells. The model is trained to reconstruct gene expression distributions for each individual cell. The mask ratio is sampled from a uniform distribution to enhance feature learning (Dong et al., 2025) (Fig. 1A). After pre-training, Stack outputs cell-level embeddings and gene expressions for new single-cell datasets, empowering numerous applications in both observational and perturbational biology in a zero-shot manner (Fig. 1A), without the need for test-data-specific fine-tuning. Stack gains strong generalization power through its inference-time learning capacity, which is detailed later.

Stack abstracts the latent state of each cell as an ensemble of token vectors, which we term “gene module tokens”. Tokens are generated by projecting gene expression vectors into a latent space using a single-layer perceptron, producing a fixed number of tokens per cell. One of the innovations we introduce is that this tokenization module is trained end-to-end alongside the rest of the model, without relying on external gene semantic information. Because the number of tokens (100) is substantially smaller than the number of genes in the data, the model must implicitly learn meaningful gene groupings to preserve biological information. This also yields substantial scalability gains over gene-level tokenization.

A key innovation of Stack is a new tabular transformer block that enables both intra-cellular and inter-cellular information flow within the cell set (Fig. 1B). Each block stacks an intra-cellular multi-head attention (MHA) layer, an inter-cellular MHA layer, and a token-wise feedforward network (FFN) layer. In the intra-cellular MHA layer, the attention mechanism is performed on the gene module token sequence independently for each cell. In the inter-cellular MHA layer, the attention mechanism is on the cell set, with “cell tokens” defined as the concatenation of all gene module tokens. Our design draws inspiration from emerging tabular learning architectures such as TabPFN and TabICL (Hollmann et al., 2025; Qu et al., 2025) and additionally accounts for attention between different gene modules across cells. The final-layer gene module tokens are concatenated to form the cell embedding, and a cell-wise multilayer perceptron (MLP) decoder models the observed gene expression as a probabilistic function of this embedding. In addition to the masked gene reconstruction objective, Stack incorporates a distributional regularization on cell embeddings that improves generalization, motivated by the linear identifiability theory of latent variable models (Khemakhem et al., 2020; Dong et al., 2024) (Methods).

During pre-training, this architecture captures the dependency between each cell and its context (the remainder of the cell set), enabling greater control and refinement of predicted single-cell expression. Following a post-training procedure, it also allows for the modification of a cell set of interest (*query* set) using an artificially designed set of *prompt* cells that implicitly encodes auxiliary conditioning to guide the final model output. This output includes both embeddings and predicted expression values for each target cell (Fig. 1C). The framework offers two key advantages at inference: (1) The dependencies across cells in a set, encoded in inter-cellular attention layers during pre-training, generalize to unseen datasets to improve zero-shot performance without model updates; (2) It provides a backbone for in-context learning, enabling “cell prompt engineering” at inference. Specifically, the context can be altered to achieve desired outcomes for the query cells, such as shifting gene expressions to match a new donor or predicting the effects of perturbations. Cell prompts can be obtained from any single-cell observational or perturbational dataset, providing high-quality representations of diverse condition signals (such as disease state, genetic/chemical perturbations, donor variability, age) in a unified transcriptomic space. Simulating query data in the prompt context enables both generalization to new biological contexts (cell types, donors etc.) as well as novel predictive tasks (such as perturbation, age etc.) not encountered during training. Both rely solely on prompt-provided information at inference time.

For model evaluation, we also trained Stack on a version of the CELLxGENE collection (Program et al., 2025) that contains 73.7 million human cells, and a 60-million-cell subset of human scBaseCount in addition to the full scBaseCount. We observed a scaling behavior in terms of various validation metrics across configurations of Stack from 69 to 629 million parameters (Fig. S1A). Increasing hidden dimensionality results in an overall improvement of validation performance. Increasing network depth yields similar validation loss but improves performance on other metrics for the full scBaseCount dataset, while showing mixed effects on the scBaseCount subset (Fig. S1B). Scaling cell set size to 256 optimizes validation loss, while validation reconstruction metrics peak at a cell set size of 128 among the tested values (Fig. S1C). An ablation study confirmed that inter-cellular attention and latent space regularization in Stack both improve validation metrics (Fig. S1D). To assess the impact of informative cell context, we varied unique cells per cell set while holding total context size constant through repetition. Stack outperforms the ablation model without intercellular attention once unique cell numbers in the set exceed 32. The larger Stack model achieves lower validation loss, with gains widening as unique cells increase, indicating enhanced information aggregation capacity (Fig. 1D).

Finally, Stack tokenization yields highly specific gene groups. Among the top-10 genes per module (ranked by tokenization weight magnitude), 526 of 699 (75.3%) appear in exactly one gene-module token, even though we impose no explicit sparsity objective. Gene set enrichment analysis confirmed that these tokens are functionally coherent (Figs. 1E, S2).

### **2.2.** Stack generates superior embeddings by learning from cellular contexts at inference time

To evaluate the utility of Stack embeddings for individual samples on downstream tasks (Fig. 2A), we developed a comprehensive benchmarking framework that assesses the impact of cellular context and the quality of single-cell level representations, through probing and integration evaluations (Fig. 2B-C, Methods). The datasets for evaluation include: 1) five collections of observational samples, four representing distinct tissues (Kidney, Lymph Node, Brain, and Lung) drawn from a large number of donors (38–223), and Tabula Sapiens (De Boer et al., 2021; Li et al., 2025; Salcher et al., 2022; Gabitto et al., 2024; The Tabula Sapiens Consortium et al., 2022), 2) four large-scale perturbation datasets covering three major perturbation modalities (Drug: OpenProblems, Tahoe-100M; Signaling: Parse-PBMC or Parse; Genetic: X-Atlas:Orion or Xaira) (Luecken et al., 2025; Zhang et al., 2025; Parse Biosciences, 2023; Huang et al., 2025). The LUCA dataset was part of the scBaseCount or CELLxGENE training data, whereas the remaining evaluation datasets were not, thus corresponding to a zero-shot setting (see Methods for details). We first evaluated model embeddings on observational single-cell data by applying linear and multi-layer perceptron (MLP) probes to predict metadata across varying levels of subtlety and resolution, ranging from disease and physiological conditions to cell types. Importantly, our probing schemes incorporate carefully designed procedures, including balanced donor cell numbers, donor-level test set holdout, and group-level cross-validation for regularization hyperparameter optimization (Methods). This rigorously tests whether the model captures biological state signatures through self-supervised learning that generalize to held-out donors.

**Figure 2.**
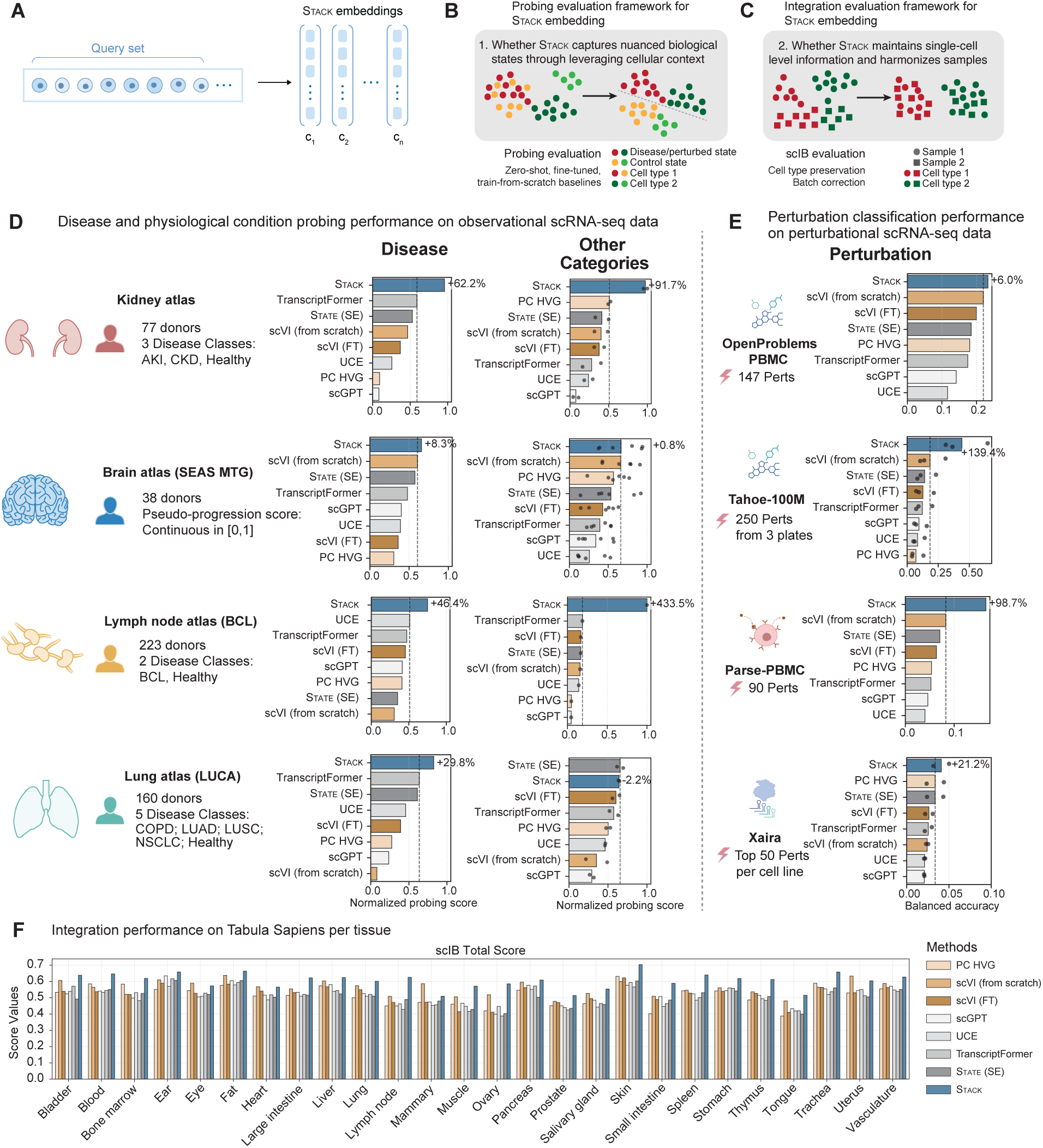
Evaluation of Stack embeddings. **A.** In the evaluations presented here, Stack serves as a context-aware embedding model, processing query cell sets from individual samples. **B.** Schematic illustration of the probing evaluation framework. **C.** Schematic illustration of the integration evaluation framework. **D.** Bar plots showing average linear probing scores for four observational atlases. Probing scores represent the balanced accuracy (for classification tasks) or Pearson *𝑟* (for regression tasks), which is first normalized for each cell type to a [0, 1] scale across all methods, and then averaged across all cell types. For the “other categories” group, each point represents one metadata prediction task (*𝑛* = 2, 6, 1, 2). AKI: Acute kidney injury. CKD: Chronic kidney disease. BCL: B-cell lymphoma. COPD: Chronic obstructive pulmonary disease. LUAD: Lung adenocarcinoma. LUSC: Lung squamous cell carcinoma. NSCLC: Non-small cell lung cancer. Other categories: Kidney (hypertension, diabetes history), Brain (Microinfarct pathology, ADNC, Braak stage, Thal phase, CERAD score, APOE4 status), Lymph node (LymphoMAP), LUCA (UICC stage, ever smoker). ADNC, Alzheimer’s Disease Neuropathologic Change. UICC, Union for International Cancer Control. **E.** Bar plots showing average linear probing scores on classifying perturbations from four datasets. Each point represents one evaluated plate for Tahoe (*𝑛* = 3) or one evaluated cell type for Xaira (*𝑛* = 2). In **D** and **E**, dashed lines indicate the performance of the best competing method in each evaluation. **F.** scIB evaluation for each tissue in Tabula Sapiens (The Tabula Sapiens Consortium et al., 2022). All Stack results presented here are based on one model with the (Large) setting pre-trained on full human scBaseCount.

We compared Stack with a list of methods that generate embeddings in zero-shot, fine-tuned, or train-from-scratch settings. These include principal components of highly variable genes (PC HVG), scGPT (Cui et al., 2024), UCE (Rosen et al., 2023), State (State Embedding or SE) (Adduri et al., 2025), Transcript-Former (Pearce et al., 2025), scVI pre-trained on the scBaseCount subset and fine-tuned on the target dataset (scVI FT) (Lopez et al., 2018; Gayoso et al., 2022), and scVI trained from scratch on the target dataset (scVI from scratch). In per-cell-type linear probing, Stack shows a substantial advantage compared with alternative methods across disease/other categories on the four observational datasets (Figs. 2D, S3A). The only exception is the classification of other categories in LUCA, a dataset partially included in the training sets of both Stack and State (SE), where Stack ranks second and underperforms State (SE) by 2.2%. To control for the information flow across cell types, we constructed a new cellular context setting where the cells included in each cell set are constrained to be of the same cell type. Stack still achieves the best overall performance among all methods, albeit by a smaller margin (Fig. S4). Removing this cellular context information by shuffling cells across donors results in a notable decrease in performance, with final results similar to those of the alternative methods (Fig. S4). To rule out the possibility that Stack’s advantage arises from simple information leakage from more informative cell types, we further evaluated probing performance in the overall best-performing cell type across all methods, and Stack’s advantage remained (Fig. S5). The association between Stack embedding performance and cellular context configurations suggests that Stack’s improvement stems from its ability to extract information from novel cellular contexts.

A similar advantage of Stack is observed with MLP probing across all four observational datasets (Fig. S3B). In either linear or MLP probing, all other advanced foundation models show inconsistent performance relative to baseline methods such as PC HVG and scVI trained from scratch on the dataset of interest (Figs. 2D, S3, S4). In terms of cell type classification, all methods demonstrate highly similar performance (Fig. S3C). The small variation in performance is likely due to the cleaner cell type signal and its manually annotated nature. The competitive performance of Stack suggests that the context-aware mechanism does not compromise Stack’s representation at single-cell resolution.

We next evaluated different models’ performance in classifying chemical, signaling, and genetic perturbations (Fig. 2E). Stack outperforms existing methods on all perturbational datasets tested and is the only one to consistently outperform scVI trained from scratch. Notably, Stack shows substantial improvements in discriminating between perturbation effects in large-scale datasets Tahoe and Parse, outperforming the best alternatives by approximately 100-140% despite being trained almost exclusively on observational data. All methods show low absolute performance in classifying genetic perturbations in the Xaira dataset, likely due to measurement noise and gene expression similarity across perturbations. These results suggest that aggregating context information is crucial for accurate predictions of subtle biological states.

We also evaluated cell type preservation and donor label correction performance of different embeddings on observational data through scIB integration metrics (Luecken et al., 2022). Across all four observational datasets, Stack ranks as the top performer, surpassing the best alternative (State (SE)) by +1.8% (Fig. S6A). In Tabula Sapiens, Stack outperforms alternative methods in 22 of 26 tissues (ranking second in eye, heart, mammary and uterus, where scVI from scratch performs best), demonstrating superior performance across human tissues (Fig. 2F). Scaling Stack training data and model size each lead to improvements in batch integration performance (Fig. S6B). UMAP visualization supports Stack’s effectiveness in clustering fine-grained cell states and integrating batch-level information (Fig. S6C). The results remain similar across model scales and extend to dataset label integration evaluations (Fig. S6D-E). These results establish Stack as a powerful embedding model that enhances zero-shot prediction and integration by leveraging cellular contexts.

### **2.3.** Stack enables in-context learning of novel predictive tasks with cells after post-training

During pre-training, Stack is exposed to sets of cells from the same biological sample (such as donor or experimental condition), limiting the base model’s utility for tasks where the context is engineered by the user to produce a desired cell state. This limitation parallels the necessity of supervised fine-tuning (SFT) and reinforcement learning (RL) for adapting pre-trained large language models to follow user instructions (Wei et al., 2022; Longpre et al., 2023; Ouyang et al., 2022). To teach Stack to follow instructions, we define a cell conditioning task that involves two cell populations: prompt and query. The prompt cells specify the desired biological state or condition, while the query cells specify the cell type of interest. The objective is to predict counterfactual states of query cells, i.e., their gene expression profiles under the biological condition of the prompt, encompassing tasks such as generalizing perturbation effects to new cell types and across datasets. Here, prompt and query cells can come from different datasets, comprise non-overlapping cell types, and their annotations may be unavailable. Therefore, a supervised approach is not feasible for this task, and a foundation model with zero-shot capabilities is crucial. The context-awareness of Stack makes it particularly suitable for these tasks.

We developed a post-training recipe via self-distillation (Zhou et al., 2021; Oquab et al., 2023) to adapt the Stack model for the task. This post-training process is closely related to masked language diffusion models (Sahoo et al., 2024), whose training objective is to recover masked tokens within input sequences with a masking ratio ranging from 0 to 1. In the post-training procedure, each cell set from a single biological sample is grouped by type and partitioned into two subsets: *prompt* cells, which remain visible, and *target* cells, which are held out, analogous to unmasked and masked tokens in masked diffusion models. During post-training, the target cells are replaced with type-matched *query* cells drawn from a different biological sample. Stack is then post-trained to reconstruct the held-out target cells from query cells, conditioned on the prompt cells. Through this post-training procedure, Stack learns to predict counterfactual states for any cell population conditioned on the prompt cells, enabling conditional cell state generation.

We use a pre-trained Stack model as a teacher to extract embeddings for the target cells (Fig. 3A). The student Stack model is optimized to predict target cell distributions in both embedding and gene expression spaces, with additional regularization terms. The teacher model is updated using exponential moving average of the student model’s parameters, allowing it to progressively adapt to new data distributions while preserving knowledge acquired during pre-training. To compute the distributional match in gene expression space as a training objective, we employ a zero-inflated normal distribution approximation that enables reparameterization. Finally, a multi-layer perceptron (MLP) classifier is trained to classify cells in the embedding space, where a lower score indicates greater similarity to the prompt condition and thus higher confidence in generation quality. At inference time, this score guides an iterative refinement procedure (Methods).

**Figure 3.**
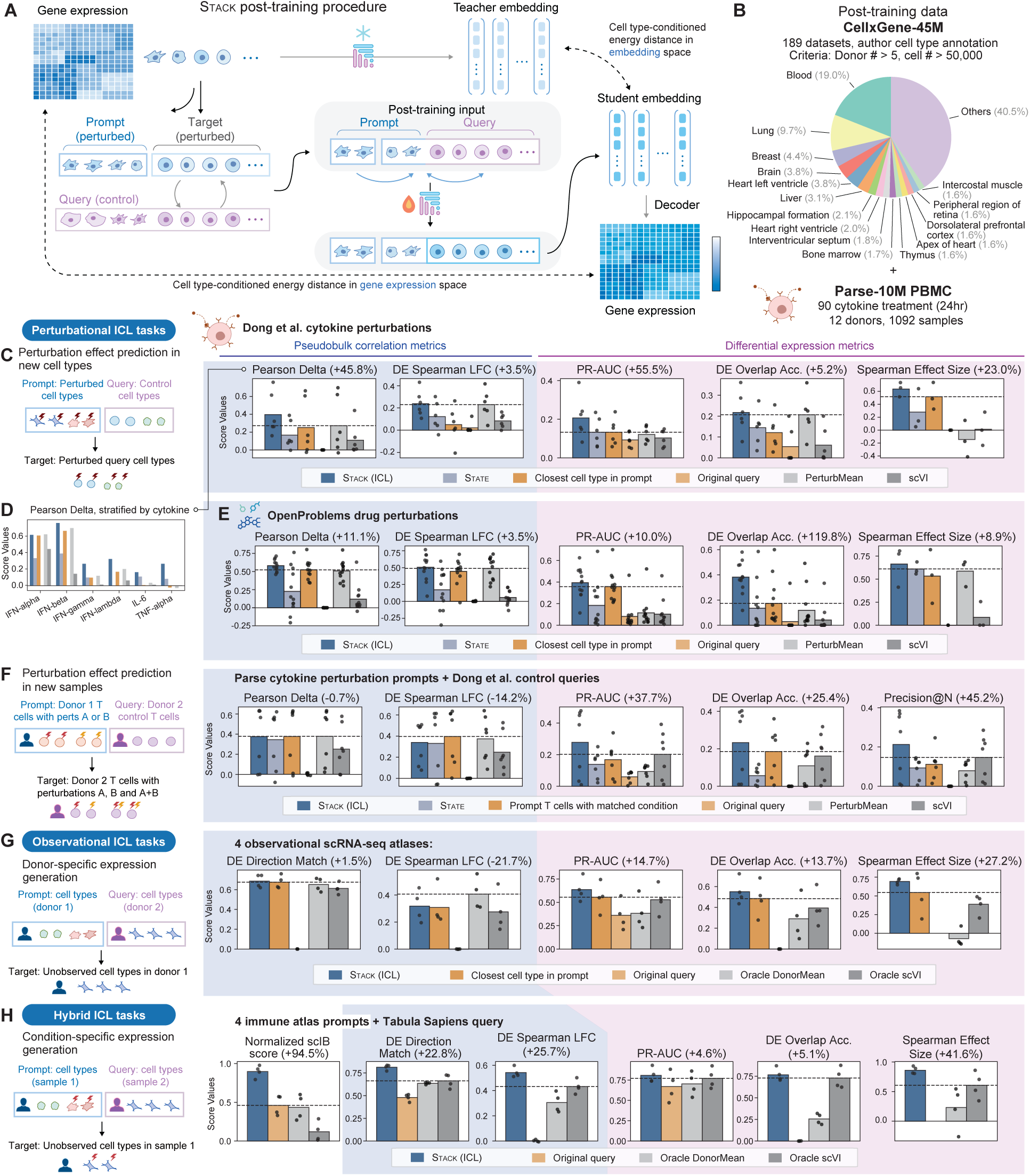
Post-training of Stack for in-context cell prompting tasks. **A.** Schematic illustration of the Stack post-training framework. Cell sets are organized by cell type and partitioned into prompt and target cell sets. During post-training, target cells are replaced with type-matched cells from different conditions (queries) to serve as model input. The model learns to predict gene expressions and embeddings of target cells via distributional alignment and self-distillation, with an MLP classifier guiding inference-time generation. **B.** Overview of post-training data. Training data comprised approximately 55M cells from curated CELLxGENE datasets and the Parse PBMC dataset (12 donors, 90 perturbations). **C.** Evaluation of perturbation effect prediction across cell types on the Dong et al. (2023) cytokine perturbation dataset (6 cytokines). In the first four panels, each point represents the average result of a cytokine condition. In the last panel, each point represents a cell type (B cells, myeloid cells, T cells). Percentages in titles represent the average improvement of Stack over the best non-Stack baseline. All Stack predictions were generated using a post-trained Stack (Large) model with a mask diffusion procedure (*𝑇* = 5). **D.** Pearson Delta evaluation results for Dong et al. (2023), stratified by cytokine. **E.** Evaluation of perturbation effect prediction across cell types on the OpenProblems drug perturbation dataset (Luecken et al., 2025) (12 drugs). Points represent drug conditions (first four panels) or cell types (last panel). **F.** Evaluation of T cell response prediction across samples. Each donor from Parse PBMC serves as a prompt, and two retained donors from (Dong et al., 2023) serve as queries. Each point represents the average result of a cytokine condition (7 single/combinatorial conditions). **G.** Evaluation of donor-specific gene expression generation across four atlases (Kidney: (De Boer et al., 2021); Lymph Node: (Li et al., 2025); Liver: (Edgar et al., 2025); PBMC: (Wells et al., 2025)). Non-overlapping cell types from sampled donor pairs serve as prompts and queries. Each point represents one evaluation dataset. **H.** Evaluation of condition-specific expression generation across four PBMC atlases. Drug perts: 15 conditions with all PBMC cell type expression profiles available in (Luecken et al., 2025); Aging: first 10 donors in (Wells et al., 2025); Parse donors: all control PBS conditions from 12 donors; Parse perts: first 20 perturbation conditions from donor 1. Immune cells from Tabula Sapiens (The Tabula Sapiens Consortium et al., 2022) serve as queries for all cases. Each point represents one evaluation case. For all scores, higher values indicate better performance. In **C** to **H**, dashed lines indicate the performance of the best competing method in each evaluation.

After post-training, we can use Stack as a conditional generative model to simulate novel cell populations. In the generative procedure, Stack receives cell sets containing concatenated prompt and query data. Stack predicts the gene expressions on all query positions, as well as their scores using the MLP classifier. At each iteration, we replace a fraction of highest-confidence query cells’ gene expression values with model predictions. This process is conducted in an iterative manner and finishes when the fraction reaches zero, at which point all query cells are replaced with predictions. Throughout the iteration, the fraction of prompt data in the input cell set is gradually increased to enable finer control (Methods).

For post-training data, we curated a large 55-million-cell scRNA-seq data collection comprising a set of large CELLxGENE datasets (>50,000 cells, >5 donors) and the Parse PBMC 10M dataset, which contains 12 donors and 90 cytokine perturbations (Fig. 3B). This training data is enriched for primary cells profiled from human tissues and blood, with a particular emphasis on immune cells. To efficiently post-train on our large data collection, we developed an extended post-training dataloader that maximizes cell-type-aware local chunking. We evaluated the post-trained model on four downstream cell prompting/in-context learning tasks spanning three categories: perturbational, observational, and hybrid ICL tasks (Box 1).

#### Box 1. In-context learning (ICL) tasks for cell prompting

- **Perturbational ICL** (prompts are perturbed cells and queries are control cells)

1. **Perturbation effect prediction in novel cell types from same sample**: We use randomly sampled perturbed cell types as the prompt, and control cells from remaining cell types as the query. Stack simulates the prompt specified perturbation condition in the query cell types (Fig. 3C–E). *Example*: Given a PBMC sample where only T cells were perturbed with IL-6, we use perturbed T cells as the prompt and unperturbed B cells/monocytes as the query to predict IL-6 effects on B cells/monocytes.
2. **Perturbation effect prediction in novel samples**: We use one single perturbed cell type (T cells here) as the prompt, and control T cells from another donor in a different dataset as the query. This task evaluates the model’s ability to recapitulate the response patterns observed in the query sample (Fig. 3F). *Example*: We collected a new PBMC sample and want to predict how T cells respond to IFN-*𝛽* stimulation based on a reference dataset. We use IFN-*𝛽*-perturbed T cells from a published reference dataset as the prompt, and control T cells from our new sample as the query.
- **Observational ICL** (prompts and queries are cells from observational scRNA-seq samples)

3. **Donor-specific expression generation**: We use sampled cell types from one donor as the prompt and non-overlapping cell types from another donor in the same dataset as the query. Stack predicts the expression profile of query cell types in the prompt donor (Fig. 3G). This task evaluates the model’s ability to capture donor-specific expression differences in observational datasets. *Example*: In a patient cohort, Patient A has profiled macrophages and T cells but lacks fibroblasts due to tissue availability. We use Patient A’s cells as the prompt and fibroblasts from Patient B as the query, to impute Patient A-specific fibroblast expression.
- **Hybrid ICL** (prompts and queries are from different observational or perturbation studies)

4. **Cross-dataset cell type generation**: We use sampled cell types from one condition as the prompt and non-overlapping cell types from another dataset as the query, and predict the query cell type profile in the prompt sample context (Fig. 3H). This task assesses the model’s capacity to generate cell types absent from an observational atlas or perturbation data of interest. *Example*: Using drug perturbed T cells and B cells as the prompt, and dendritic cells from an independent scRNA-seq atlas as the query, we predict perturbation responses in dendritic cells that were never experimentally perturbed.

Our evaluation leverages key metrics proposed in Cell-Eval (Adduri et al., 2025), which can be categorized into pseudo-bulk correlation metrics (Pearson Delta, DE Spearman LFC) and differential expression (DE) metrics (PR-AUC, DE overlap accuracy, Spearman effect size; see Methods). We also report the Jaccard similarity metric, which measures the overlap between two DE gene sets normalized by their union, thereby adjusting for both predicted and ground-truth DE gene set sizes. DE direction match and DE precision-at-N serve as auxiliary metrics when pseudobulk or DE metrics are less appropriate or not applicable (Methods). For task 4, where target and query belong to different datasets, we additionally evaluated scIB batch integration metrics. Alternative baselines include the query data itself (input baseline), the nearest/same cell type in the prompt sample, State (Adduri et al., 2025), and two strong baselines identified in Adduri et al. (2025) (PerturbMean/DonorMean, scVI). For perturbational ICL tasks, we employ a “synthetic control” approach that uses the unperturbed version of the prompt sample to additionally predict a control profile, which then serves as the reference for the perturbation effect predictions when computing Cell-Eval metrics. The same synthetic-control procedure is applied to the closest/same-cell-type baseline and scVI, resulting in stronger baselines. For the remaining tasks, Stack operates without auxiliary samples, whereas DonorMean and scVI baselines still require them; we therefore designate these baselines as “oracle” in those cases. The evaluation data includes seven datasets that were **never seen** during either Stack pre-training or post-training: 1. Open-Problems drug perturbation (Luecken et al., 2025), 2. Cytokine stimulation (Dong et al., 2023), 3. Immune aging (Wells et al., 2025), 4. Tabula Sapiens (The Tabula Sapiens Consortium et al., 2022), 5. Kidney atlas (De Boer et al., 2021), 6. Lymph node BCL (Li et al., 2025), 7. Liver atlas (Edgar et al., 2025). Additionally, we include the Parse PBMC dataset (Parse Biosciences, 2023) as prompts (not as queries) in our evaluations.

In task 1 (perturbation effect prediction across cell types) for the Dong et al. (2023) cytokine stimulation dataset, Stack not only exhibits an advantage across all metrics (Figs. 3C, S7), but generalizes the global effect of subtle cytokine perturbations, including IL-6 and TNF-*𝛼* across cell types, as demonstrated by Pearson Delta (Fig. 3D). The overall advantage also extends to the OpenProblems drug perturbation dataset, which comprises drug conditions unseen during model pre-training or post-training (Figs. 3E, S7). In task 2, Stack outperforms alternative methods (including perturbation effects of observed prompt T cells) in DE metrics but performs similarly in pseudobulk metrics, across both individual and combinatorial cytokine conditions. Because Parse and Dong et al. (2023) use different stimulation protocols in dosage and stimulation duration, generalizing subtle effects of several cytokines (TNF-*𝛼*, IL-6) alone yields near-zero pseudo-bulk and DE scores across all methods (Fig. 3F).

In task 3, oracle DonorMean shows the strongest performance in pseudobulk correlation metrics, while Stack maintains its lead in directional and DE metrics across four evaluated tissues (Figs. 3G, S7). Notably, Stack demonstrates particular strength in generating cell types across datasets (Task 4), as evidenced by scIB integration and Cell-Eval metrics (Figs. 3H, S7). This advantage may stem from the context-aware capacity acquired during Stack pre-training, as supported by the suboptimal performance of Stack trained from scratch on perturbational ICL tasks (Fig. S8).

As an in-context learning approach, Stack achieves the strongest overall performance across all ICL tasks, ranking first in 28 of 31 evaluation metrics, including those shown in Figs. 3C–H and the Jaccard similarities in Fig. S7. Closest/same-cell-type prompt cells emerge as a strong baseline when evaluated on Cell-Eval metrics, which has not been sufficiently addressed in previous studies. Despite being trained on both prompt and query data, State is outperformed by Stack across all metrics on the evaluated perturbation tasks. This is likely because State was designed for tasks with orders of magnitude more supervised data than those evaluated here. Stack’s success in low-data, cross-experiment tasks demonstrates its utility as a foundation model when supervised fine-tuning is insufficient, for example, when interrogating cell types that are difficult to perturb experimentally. The multi-step mask diffusion generative procedure shows modest advantages over Stack one-step prediction and alternative generative schemes across ICL tasks (Fig. S9). The competitive performance of Stack on unseen prompts and queries (Dong et al., 2023; Wells et al., 2025; The Tabula Sapiens Consortium et al., 2022), novel perturbations (Luecken et al., 2025), and cell types and tissues beyond peripheral blood (De Boer et al., 2021; Li et al., 2025; Edgar et al., 2025; Sikkema et al., 2023) underscores the generalizability of our approach to previously unencountered datasets and diverse biological tasks.

### 2.4. Stack generates a virtual whole-organism perturbational atlas

We employed Stack to construct a perturbational whole-organism atlas (*Perturb Sapiens*) via ICL. We selected all conditions with sufficient cell counts from (Parse Biosciences, 2023; Luecken et al., 2025; Zhu et al., 2025) as prompts (Methods), comprising 90 cytokine perturbations across 12 donors (Parse Biosciences, 2023), 136 drug perturbations across 3 donors (Luecken et al., 2025), and 666 genetic perturbations across 4 donors (Zhu et al., 2025). The tissue-balanced Tabula Sapiens compendium served as the query (Fig. 4A, Methods) (The Tabula Sapiens Consortium et al., 2022). UMAP visualization demonstrates a single-cell resolution map with tissue-specific and cell-type-specific expression for *Perturb Sapiens* (Figs. 4B, S10A–B). Inspection of the MLP classifier scores suggests a variation in generation confidence for different cell and tissue types (Figs. S10C–E). The classifier assigns low confidence (high logit value) to several rare cell types such as transitional epithelial cells across all perturbation modalities (drug/cytokine/genetic). We restricted our subsequent analyses to cells with logits smaller than a threshold (2.5) suggesting high confidence.

**Figure 4.**
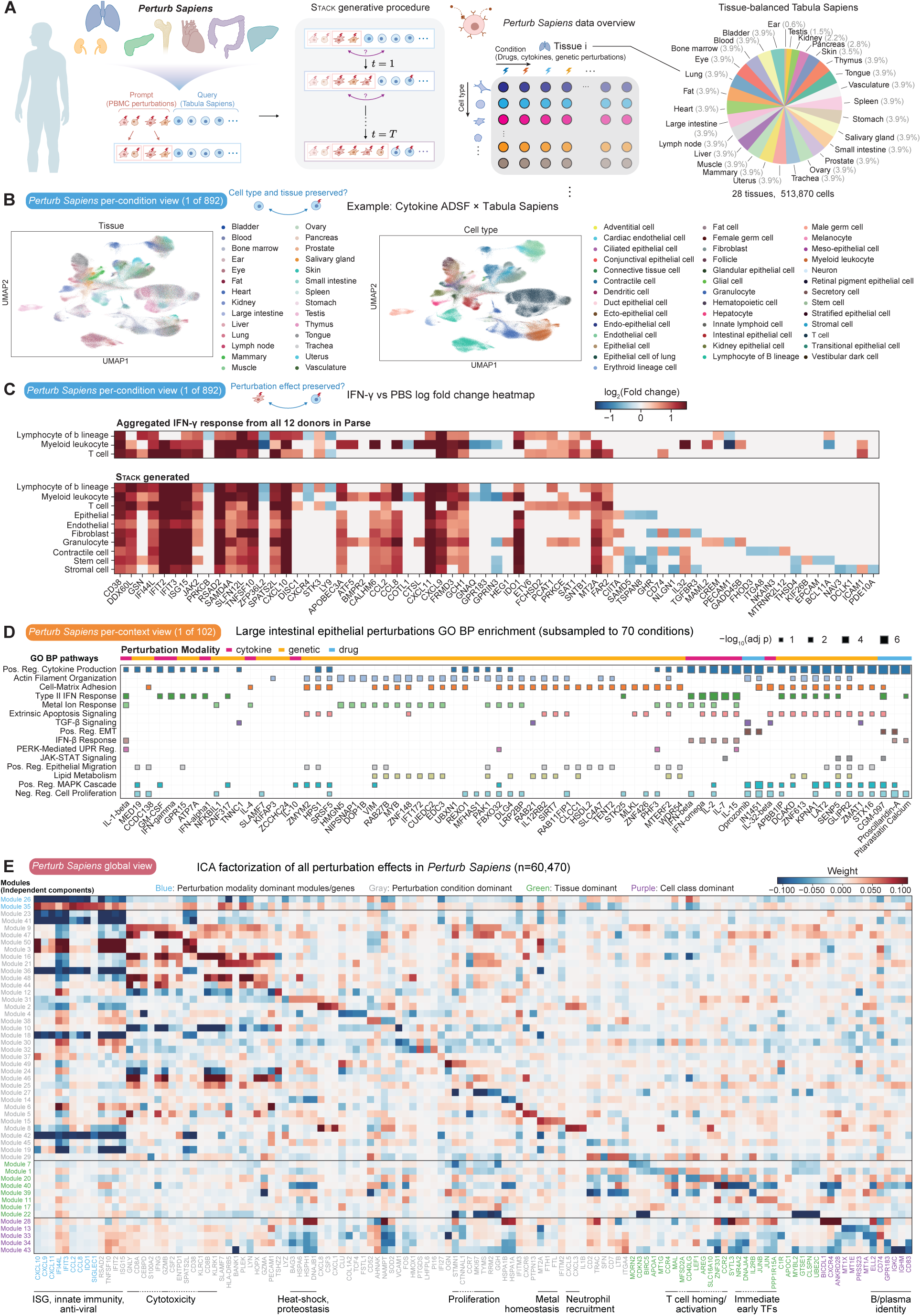
Analysis of a perturbational whole-organism atlas *Perturb Sapiens*. A. Overview of *Perturb Sapiens*. We utilize PBMC perturbation datasets (90 cytokines from Parse Biosciences (2023); 136 drugs from Luecken et al. (2025); 666 genetic perturbations from Zhu et al. (2025)) as prompts and Tabula Sapiens (The Tabula Sapiens Consortium et al., 2022) as queries (left). For each tissue and cell type, *Perturb Sapiens* comprises simulated gene expression profiles under drug, cytokine, and genetic perturbations. The resulting atlas spans 28 tissues and 513,870 cells per perturbation condition (right) (The Tabula Sapiens Consortium et al., 2022). **B.** UMAP visualization of a representative *Perturb Sapiens* condition (cytokine ADSF), colored by tissue and cell type. ADSF, adipose tissue-specific secretory factor. **C.** Log2-fold-change (LFC) heatmap comparing IFN-*𝛾 Perturb Sapiens* to control. Significantly differentially expressed genes (DEGs) are shown in color; non-significant genes are shown in gray. LFCs were aggregated as the median across donors, with per-donor *𝑝*-values combined using a Beta-distribution test based on the median *𝑝*-value (Methods), followed by Benjamini–Hochberg FDR correction. **D.** Gene Ontology (GO) Biological Process enrichment dot plot for large intestinal epithelial perturbations. For each perturbation, DEGs with positive LFCs were tested for enrichment against the GO Biological Process 2025 gene set library using Enrichr (Fang et al., 2023; Kuleshov et al., 2016; Aleksander et al., 2025). Dot size encodes − log_10_(adjusted *𝑝*-value), capped at 8; only gene set terms with adjusted *𝑝 <* 0.05 are shown. **E.** Heatmap of the ICA loading matrix derived from *Perturb Sapiens* perturbation effect profiles (significance-filtered LFC, aggregated across donors). For each module (independent component), the top five genes ranked by absolute loading magnitude are shown. Each module is annotated by its dominant source of variation, including perturbation modality, condition, tissue, or cell class (Methods).

As a representative example, we inspected the effect of IFN-*𝛾* perturbation versus control in *Perturb Sapiens* to assess model generalization beyond the effects observed in the prompt immune cell types. Stack generates a highly realistic differential expression map with cell-type specificity (Fig. 4C). The top differentially expressed genes exhibit near-perfect concordance between prompt and generated immune cells. Known downstream targets of IFN-*𝛾* were activated across a variety of immune and non-immune cell types (e.g., *IFIT3, ISG15, CXCL11, IDO1*). In non-immune cell populations, IFN-*𝛾* induced the antigen-presentation regulator *CIITA* together with *CD74*, while concurrently suppressing lineage and structural/adhesion programs (e.g., *EPCAM, PECAM1, ITGA8, THSD4*). This pattern was accompanied by changes in remodeling and motility-associated genes (*TGFBR3, FHOD3, KIF26B, NAV3, ICAM1, GADD45B*), consistent with IFN-driven inflammatory reprogramming of non-immune compartments. 8/12 of these genes are not differentially expressed in the Parse PBMC data, while the remaining four show differential expression in a subset of PBMC cell types. This indicates that Stack successfully generalized beyond the prompt data to capture immunomodulatory and remodeling responses specific to non-immune lineages. To test whether this generalization extends to drug conditions unseen during training, we next examined dactolisib, a PI3K/mTOR inhibitor that suppresses rather than activates immune signaling. Dactolisib perturbation leads to global repression of interferon-stimulated genes (ISGs) (Fig. S11). Non-immune lineages showed selective remodeling among cytoskeletal/adhesion and stress-response programs (e.g., *TUBA1C, FHL1, LAMA2, NINJ1, MT1X*).

For prompts that only contain T cells, Stack also generates high-quality predictions across all cell types as observed for ketoconazole, *RNF14*, and *FBXW7* (Figs. S12–S14). Notably, ketoconazole exhibited pronounced donor-specific effects across the OpenProblems dataset. *Perturb Sapiens* effectively captures these individualized responses in the prompt donor while maintaining alignment with bulk expression patterns for a number of genes where donor-specific differential expression is not detected (Fig. S12). *RNF14* perturbation induced a proliferative/activation program (*MKI67, TOP2A, BIRC5, IFNG, CD69*), with non-immune lineages showing distinct remodeling of contractile and ECM pathways (*MYL9, TPM1, MYLK, SPARC, FSTL1*, Fig. S13). *FBXW7* perturbation induced a modest response, with an activation signature in immune cells and an ECM/stromal remodeling signature in non-immune lineages (Fig. S14). Both genetic perturbations differ from canonical cytokine signatures characterized by ISG and antigen-presentation induction, supporting Stack’s ability to generalize beyond the Parse training set.

Beyond examining all tissue and cell-type contexts within a single perturbation, *Perturb Sapiens* also enables a complementary view by surveying responses across perturbation conditions. To illustrate this, we evaluated Gene Ontology Biological Process (GO BP) enrichment of up-regulated genes across all perturbation conditions in *Perturb Sapiens* large intestinal epithelial cells (Fig. 4D). Several programs, including cytokine production, mitogen-activated protein kinase (MAPK) cascade, and regulation of cell proliferation, were shared across a majority of drug, cytokine, and genetic perturbations. Other programs were enriched more selectively, either within specific perturbation modalities (e.g. epithelial–mesenchymal transition (EMT)) or across a limited set of perturbations spanning modalities (e.g. TGF-*𝛽* signaling, IFN-*𝛽* response, and PERK-mediated unfolded protein response). Together, these enrichment patterns indicate that *Perturb Sapiens* captures both shared and perturbation-specific biology at the gene-set level, complementing single-gene analyses (Fig. 4C) and enabling systematic query and comparison of perturbation states across conditions and modalities.

Finally, we computed log2-fold-changes (LFCs) and statistical significance of perturbation effects across all available conditions in *Perturb Sapiens*, aggregating effects from multiple donors and stratifying by cell class, tissue, and prompt donor (Fig. S15A, Methods). We defined perturbation effect profiles by retaining only statistically significant LFCs and setting non-significant values to zero, yielding a summary of perturbation effects for each condition. After discarding conditions without significant differentially expressed genes (DEGs), this procedure yielded 60,470 pseudo-bulk perturbation effect profiles spanning 716 perturbations. The resulting compendium enables a unified, multi-scale characterization of drug, cytokine, and genetic perturbations. At a summary level, predicted perturbation similarities clustered coherently by cell lineage and tissue proximity (Fig. S15B–D). At finer resolution, we compressed the concatenated perturbation effect space into 50 dimensions using independent component analysis (ICA), distilling dominant response modules that segregated by perturbation modality, condition, tissue, and cell class (Figs. 4E, S16). Modality-specific components were enriched for interferon-stimulated chemokines and innate immune genes (*CXCL10*, *CXCL9*, *CXCL11*, *IFI44L*, *IFIT3*, *CCL2*, *CCL8*, *IDO1*, *SIGLEC1*). Perturbation-specific modules spanned a broader functional repertoire, encompassing antiviral effectors (*RSAD2*, *IFIT2*, *ISG15*), cytotoxic lymphocyte signatures (*GNLY*, *CD8A*, *CD8B*, *GZMB*, *GZMA*, *IFNG*, *KLRC1*, *KLRK1*), heat-shock and proteostasis genes (*HSPA6*, *HSPH1*, *DNAJB1*, *HSPA1B*, *HSPA1A*, *BAG3*), neutrophil-recruiting chemokines (*CXCL8*, *CXCL1*, *CXCL5*, *CXCL3*), stromal and extracellular matrix remodeling factors (*COL1A2*, *FSTL1*, *OGN*, *PI16*), proliferation markers (*MKI67*, *TYMS*, *RRM2*, *STMN1*), and metal-homeostasis genes (*MT2A*, *FTH1*, *FTL*). Tissue-specific components highlighted genes involved in cell-cycle regulation (*CDKN3*, *BIRC5*, *MYBL2*, *GTSE1*, *UBE2C*), lipid metabolism (*APOA1*, *APOC1*), T cell homing and activation (*CCR4*, *MAL*, *CD40LG*, *LEF1*, *CCR2*, *IL2RB*), and immediate-early transcription factors (*JUNB*, *JUN*, *NR4A2*, *PPP1R15A*). Cell-class-specific modules featured markers of B cell and plasma cell identity (*IGKC*, *IGHM*, *ELL2*, *CD74*, *CD83*), metallothionein family members (*MT1X*, *MT1E*, *MT1A*), and migration-associated receptors (*CXCR4*, *GPR183*). Notably, these component classifications are not strictly exclusive and can reflect co-modulated perturbation signatures. For example, module 7, driven by the afore-mentioned cell-cycle genes, exhibits comparable variance contributions from cell class and tissue (Fig. S17), pointing to a tissue- and cell-class-specific proliferative response axis. Collectively, *Perturb Sapiens* provides a comprehensive, multi-resolution view of perturbation effects across cell types and tissues, representing a rich resource that extends well beyond the analyses presented here.

### 2.5. Systematic validation of *Perturb Sapiens*

We performed a comprehensive evaluation of *Perturb Sapiens* using available perturbation measurements. For drug and cytokine perturbations with multiple cell types represented in the prompt, we used the prompt itself as the reference. We then compared *Perturb Sapiens* predictions of immune and epithelial cells with biological replicates from the same perturbation dataset to assess which more closely recapitulated the cell-type-specific perturbation effects specified by the prompt. Within matched cell types, *Perturb Sapiens* achieved higher scores on 3/5 and 4/5 Cell-Eval metrics across cytokine and drug perturbations respectively, with the strongest gains observed for DE overlap accuracy, although Pearson delta scores remained low, likely due to batch effects (Fig. S18). We further evaluated the cell-type specificity of *Perturb Sapiens* by computing Cell-Eval scores for each predicted perturbation response against ground-truth profiles from all immune and epithelial cell types. In each comparison, the matched cell type ranked highest, while epithelial cell types consistently scored lower, confirming that *Perturb Sapiens* predictions are cell-type specific (Fig. S18, Methods).

We next evaluated predictions of cytokine and genetic perturbation responses in *Perturb Sapiens* non-immune profiles, focusing on the epithelial lineage given the availability of *in vitro* epithelial perturbation datasets (Koh et al., 2023; Lee et al., 2022; Swindell et al., 2018; Replogle et al., 2022) (Fig. 5A). We first benchmarked the model’s ability to reproduce the effects of five cytokines (type I IFN, IL-13, IL-1*𝛽*, TNF-*𝛼*, IL-17A) in single-cell or bulk data using either direct prediction of each cytokine or functionally similar cytokines. Our evaluation used aggregated *Perturb Sapiens* predictions across all Parse prompt donors (Methods). Stack demonstrated strong performance in generating epithelial-specific type I IFN responses, outperforming both prompt and generated immune cells (Fig. 5B). Unlike IFN-*𝛽*, IL-13 is a Th2 effector cytokine that produces weaker transcriptional signals and was administered for a different duration (7 days) in the evaluation data (Koh et al., 2023). *Perturb Sapiens* still demonstrated cell-type and tissue-specific alignment with the real experiment, albeit with overall lower scores (Fig. S19A). *Perturb Sapiens* prediction specificity generalized to other tissues, as observed in the IL-1*𝛽* keratinocyte and TNF-*𝛼* intestinal epithelial stimulation experiments (Figs. 5C, S19B). For IL-17, both the Parse prompt and generated data showed negative association with epithelial response, with absolute values exhibiting cell-type and tissue specificity (Fig. S19C), likely reflecting stimulation duration differences (1 day vs. 7 days) and the context-dependent effects of IL-17 signaling (Chong et al., 2020; Gaffen, 2009).

**Figure 5.**
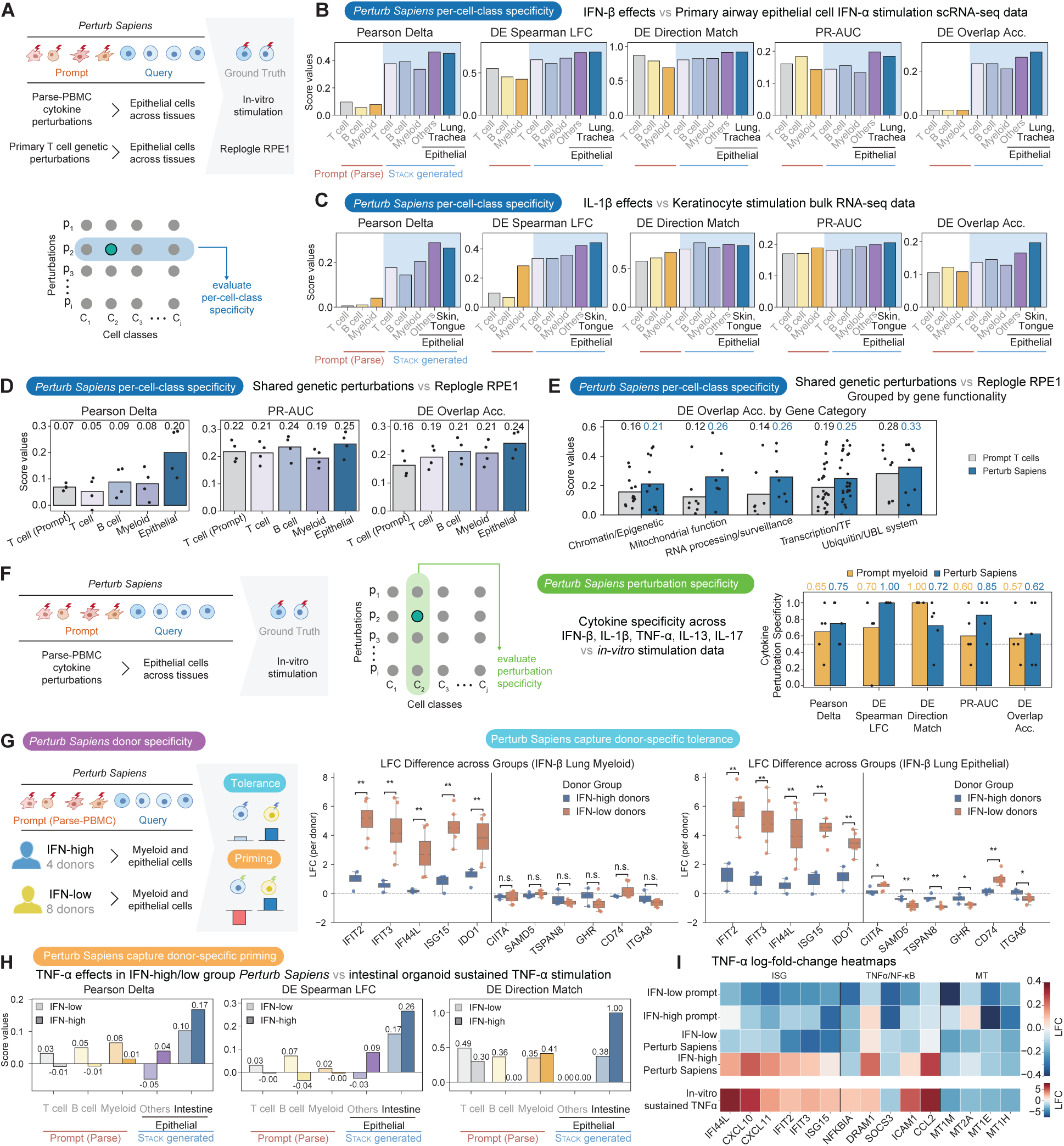
Comprehensive validation of *Perturb Sapiens*. **A.** Overview of *Perturb Sapiens* per-cell-class evaluation. We compare *Perturb Sapiens* tissue-specific and general epithelial predictions with available *in vitro* measurements of cytokine and genetic perturbations in epithelial cells from various tissues and cell lines (Koh et al., 2023; Lee et al., 2022; Swindell et al., 2018; Replogle et al., 2022). **B.** Evaluation of *Perturb Sapiens* epithelial interferon-beta (IFN-*𝛽*) effects using single-cell IFN-*𝛼* stimulation data from primary airway epithelial cells (Koh et al., 2023). **C.** Evaluation of *Perturb Sapiens* epithelial interleukin-1 beta (IL-1*𝛽*) effects using bulk IL-1*𝛽* stimulation data from primary keratinocytes (Swindell et al., 2018). **D.** Evaluation of *Perturb Sapiens* genetic perturbation effects against whole-genome Perturb-seq of immortalized retinal pigment epithelial (RPE1) cells (Replogle et al., 2022). Each point represents the average result for one prompt donor (*𝑛* = 4). **E.** DE overlap accuracy of *Perturb Sapiens* genetic perturbations, grouped by genetic perturbation categories (Table S1). Bars show mean accuracy across four epithelial donors (D1–D4) for *Perturb Sapiens* and Prompt T cells; individual donor-gene values are overlaid as points (*𝑛* = 13, 8, 7, 25, 8). In **D** and **E**, *Perturb Sapiens* epithelial cells refer to selected epithelial populations from the lung and trachea. Only conditions with a positive DE Spearman LFC between the prompt and the RPE1 dataset are shown. **F.** Evaluation of *Perturb Sapiens* prediction specificity on 5 cytokines. Specificity is defined as the normalized rank percentile of a prediction’s matched performance relative to all other available perturbations, where higher values indicate greater specificity (Methods). Bars show mean specificity across cytokines for each evaluation metric; individual cytokine values are overlaid as points. **G.** Donor-specific perturbation responses captured by *Perturb Sapiens*. Prompt donors (Parse Biosciences, 2023) were stratified into two groups based on baseline interferon activity (IFN-high: donors 1, 3, 4, 10, *𝑛* = 4; IFN-low: remaining donors, *𝑛* = 8). *Perturb Sapiens*-predicted log2-fold-changes for lung myeloid and epithelial cells are compared between the two groups. Significance was assessed by two-sided Mann–Whitney *𝑈* test: * *𝑝 <* 0.05, ** *𝑝 <* 0.01. For all box plots in this study, the center line indicates the median, boxes indicate the interquartile range, and whiskers indicate 1.5 times the interquartile range. **H.** Evaluation of *Perturb Sapiens* IFN-high/low group epithelial tumor necrosis factor alpha (TNF-*𝛼*) effects using intestinal TNF-*𝛼* stimulation data (Lee et al., 2022). Donor 1 was removed due to different response patterns with remaining donors (Klein et al., 2025). **I.** Mean LFC heatmaps for *Perturb Sapiens* predictions and the prompt, stratified by donor group, alongside *in vitro* TNF-*𝛼* response profiles (Lee et al., 2022).

For genetic perturbations, we compared *Perturb Sapiens* epithelial cells with Replogle et al. (2022), which measured genetic perturbation responses in immortalized retinal pigment epithelial (RPE1) cells. Notably, the RPE1 data exhibit systematic perturbation effects that confound comparisons (Viñas Torné et al., 2025), which we mitigated using a pseudo-control procedure (Fig. S20, Methods). Following this correction, *Perturb Sapiens* epithelial cells exhibit clear cell-type specificity and achieve overall improved Cell-Eval metrics across all 4 donors (Figs. 5D, S21). Subsetting *Perturb Sapiens* epithelial cells to lung and tracheal epithelial populations yields improved performance compared to using all epithelial cells, likely reflecting enrichment of regulatory programs shared between these cells and RPE1 (Fig. S21). Absolute scores for both *Perturb Sapiens* and prompt remain moderate, consistent with differences in stimulation protocols and readout timing between the two experiments (Zhu et al., 2025; Replogle et al., 2022). The advantage of *Perturb Sapiens* in DE overlap accuracy for genetic perturbations persists across diverse perturbation gene functional categories (Fig. 5E).

A perturbational atlas should resolve distinct condition-specific programs rather than only shared stress or inflammatory signatures. To evaluate this, we assessed the perturbation specificity of *Perturb Sapiens* by ranking each predicted profile against all observed perturbations and measuring the Cell-Eval score rank of the correctly matched condition (Methods). Across the five evaluated cytokines, *Perturb Sapiens* outperformed prompt myeloid cells on all metrics except DE direction match (Fig. 5F). The distinct behavior of DE direction match is likely due to its sensitivity to DEG list overlap.

Beyond cell-type, tissue, and perturbation specificity, differences in the baseline state of the prompt donor can further shape perturbation responses. To validate this, we examined cytokine perturbations for which previous analysis had identified two donor subpopulations with distinct baseline interferon (IFN) stimulation profiles (Oesinghaus et al., 2025). A high baseline IFN state could modulate perturbation responses through priming, tolerance, or other mechanisms (Ivashkiv and Donlin, 2014). For IFN-*𝛽*, *Perturb Sapiens* captures the attenuated ISG activation observed in the IFN-high donor group in both lung myeloid and epithelial cells. In lung epithelial cells, it additionally predicts donor-dependent differences in antigen presentation and epithelial identity genes (Fig. 5G). This observation is consistent with the previously reported refractory effect of IFN signaling on subsequent type I IFN responses (Sarasin-Filipowicz et al., 2009; Mudla et al., 2020).

For TNF-*𝛼*, the IFN-high group shows a stronger association with *in vitro* sustained TNF-*𝛼* stimulation signatures than the IFN-low group (Fig. 5H). At the gene level, *Perturb Sapiens* correctly predicts upregulation of ISGs and 7/12 NF-*𝜅*B signaling genes in the IFN-high group, whereas these genes are mostly downregulated in the IFN-low group (Table S2). This pattern suggests that *Perturb Sapiens* is not simply recapitulating the prompt: in prompt myeloid cells, these genes remain predominantly downregulated, with only scattered additional upregulation in the IFN-high group. Both subgroups show downregulation of metallothionein genes, consistent with real stimulation measurements (Fig. 5I, Table S2). Finally, we confirmed that the IFN-low group shows additional association with the 24-hour keratinocyte epithelial response signature (Swindell et al., 2018) relative to the IFN-high group, supporting the specificity of *Perturb Sapiens*’s donor-level predictions (Table S3). In summary, *Perturb Sapiens* predicts that high baseline IFN activity shifts epithelial TNF-*𝛼* responses toward a more inflammatory state, in contrast to its refractory effect on IFN-*𝛽* responses, aligning with prior knowledge and independent experiments. Our comprehensive evaluations demonstrate the validity and specificity of *Perturb Sapiens* across cell type, tissue, perturbation condition, and donor context.

### 2.6. Stack exhibits donor-specific cytokine response prioritization ability

By jointly training on large-scale donor and perturbation data, the Stack framework is uniquely positioned to capture both donor-specific expression variation and perturbation-induced responses. We hypothesized that Stack could be used to characterize donor-specific perturbation responses, a challenging and practically important task beyond the capacity of existing perturbation prediction models. The Stack post-training procedure does not explicitly target this task, as it focuses on in-context learning for either an individual donor or a single perturbation at a time. We therefore asked whether this more specific capability could arise as a byproduct of large-scale training on diverse donor and perturbation contexts.

Evaluating a method’s capacity for donor-specific response prediction requires perturbation profiles across donors with diverse genetic and disease backgrounds. However, existing perturbation datasets lack such diversity and are therefore insufficient for this evaluation. To address this gap, we collected DiseasePert-3M, the first single-cell perturbation dataset to capture disease and donor diversity at scale. DiseasePert-3M comprises 40 donors: 32 patients spanning 14 diseases (7 cancers, 6 autoimmune diseases, and osteoarthritis) and 8 healthy controls (Fig. 6A, Table S4). Perturbations in DiseasePert-3M include a panel of 11 cytokines spanning a range of functional categories (pro-inflammatory, anti-inflammatory, Th1, Th2, *𝛾*c chain, type II interferon, and growth factor; Fig. 6B). The preprocessed dataset contains 3,330,726 cells from the T/NK lineage, encompassing CD4 T, CD8 T, and NK cell populations (Fig. S22). In our analysis, we focused on three abundant and well-characterized T cell subpopulations: naive CD4 T cells, memory CD4 T cells, and effector CD8 T cells. Across these populations, DiseasePert-3M reveals clear donor-cytokine response heterogeneity, as evidenced by diverse donor response similarity patterns with high variance across cytokines (Figs. S23, 6C).

**Figure 6.**
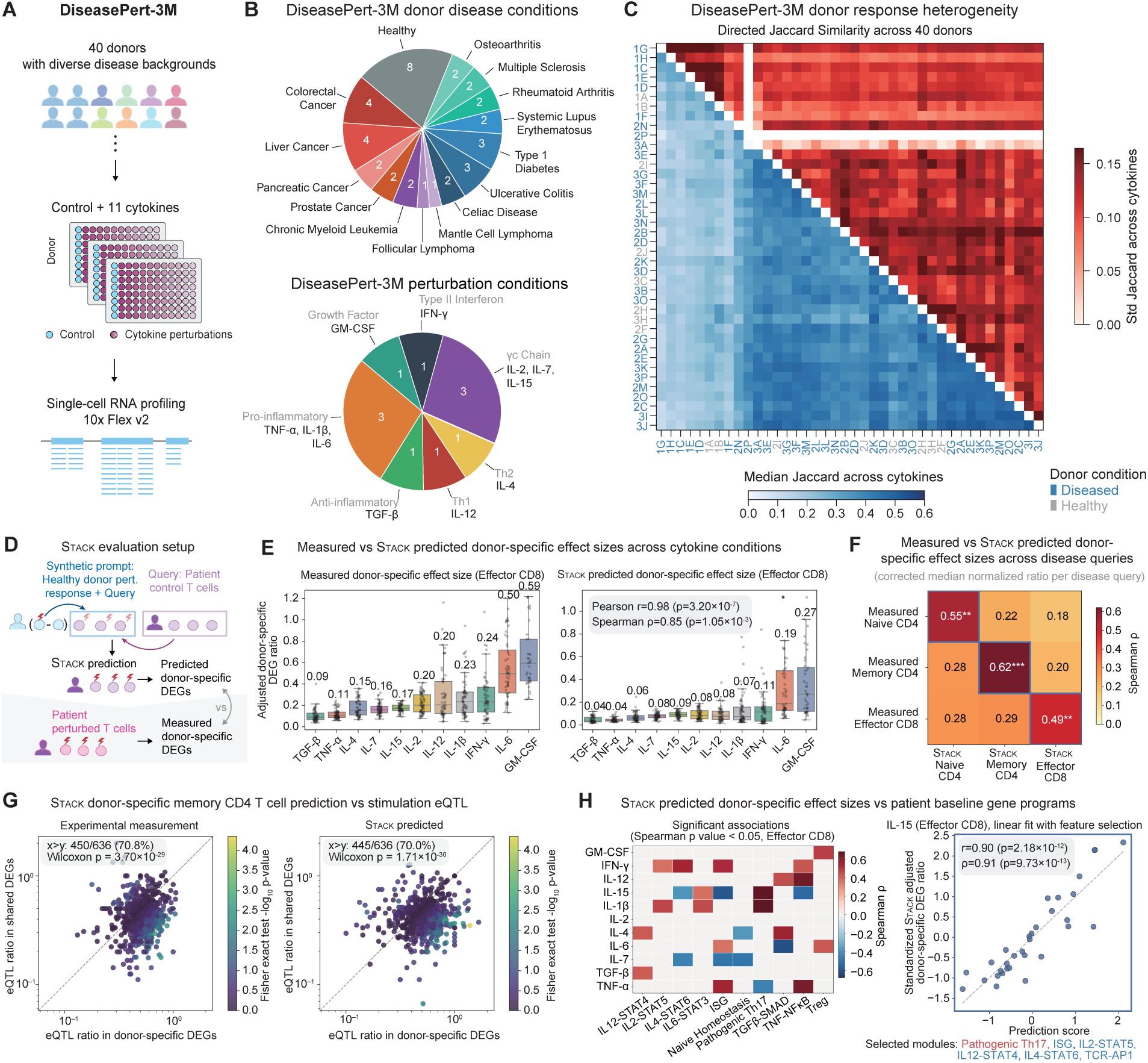
Stack enables donor-specific effect prioritization across perturbations and diseases. **A.** Overview of the DiseasePert-3M data collection workflow. **B.** Summary of donor disease characteristics and cytokine perturbation conditions in DiseasePert-3M. **C.** Summary of donor-donor directed Jaccard similarity across cytokines, averaged over 3 T cell subtypes (Naive CD4, Memory CD4, Effector CD8). Lower triangle: median directed Jaccard similarity across all cytokine perturbations for each donor pair. Upper triangle: standard deviation, reflecting variability of donor concordance across perturbations (Methods). Tick label colors indicate donor disease condition. **D.** Evaluation setup for donor-specific T cell response prediction. Unperturbed patient T cells of a given subtype serve as queries. Synthetic prompts are constructed by adding the healthy-donor perturbation-to-control difference from the same subtype to the patient unperturbed expression profile (Methods). Stack donor-specific DEGs are derived by subtracting healthy-donor perturbation DEGs from Stack-predicted DEGs, and are evaluated against measured donor-specific DEGs. **E.** Box plots of effector CD8 adjusted donor-specific DEG ratios per cytokine perturbation for real measurements (left) and Stack predictions (right). Each point represents a prompt–query pair (*𝑛* = 638); numbers above each box indicate the median adjusted ratio. Pearson and Spearman correlations between measured and Stack-predicted median adjusted ratios (*𝑟*, *𝜌*), along with their associated *𝑝*-values, are annotated. **F.** Heatmap of Spearman correlation coefficients between measured and Stack predicted adjusted median donor-specific DEG ratios across disease queries (*𝑛* = 32). Each entry is annotated with the Spearman correlation (*𝜌*) and its significance level (^∗∗^ *𝑝 <* 0.01, ^∗∗∗^ *𝑝 <* 0.001). **G.** Scatter plots comparing memory CD4 eQTL gene ratios in donor-specific versus shared DEGs between prompt-query pairs (*𝑛* = 636) for real measurements (left) and Stack predictions (right). The dashed line represents *𝑦* = *𝑥*. Each point is colored by the − log_10_ *𝑝*-value of a two-sided Fisher’s exact test on the contingency table of DEG specificity (donor-specific versus shared) by eQTL status. A one-sided Wilcoxon signed-rank test assesses whether donor-specific DEGs are more enriched for eQTLs than shared DEGs. **H.** Association between Stack-predicted donor-specific effect sizes and patient baseline gene programs in effector CD8 T cells. Left: Heatmaps of Spearman correlation coefficients between predicted donor-specific effect sizes and patient baseline gene module scores, computed as the mean gene-level *𝑧*-score within each gene set (Table S5). Only significant Spearman *𝜌* values (*𝑝 <* 0.05) are shown in color. Right: Standardized Stack-predicted IL-15 donor-specific effect sizes plotted against prediction scores from linear regression models using elastic net-selected gene modules (Methods). Each point represents one query donor (*𝑛* = 32). The dashed line indicates a least-squares linear fit to the plotted points, shown for visualization. Pearson (*𝑟*) and Spearman (*𝜌*) correlation coefficients with corresponding *𝑝*-values are shown. Selected gene modules are indicated in red for positive regression coefficients and blue for negative regression coefficients.

By design, Stack captures the biological condition of the prompt donor, but not that of the query donor. To account for query donor heterogeneity, we developed a synthetic prompt strategy with blending that transfers the perturbation-induced expression shift from the original prompt donor onto the baseline expression profile of the query donor (Methods). The synthetic prompt encodes both the perturbation condition and the query context, at the cost of producing unrealistic gene expression patterns unseen during training. We hypothesized that Stack could automatically extract both sources of information from the synthetic prompt, compensating for its imperfections to effectively perform donor-specific response prediction. For evaluation, we used unperturbed patient T cell profiles as queries and constructed synthetic prompts from perturbed healthy donor profiles together with the queries. We then derived predicted donor-specific DEGs by subtracting healthy donor perturbation DEGs from Stack-predicted ones, and evaluated them against measured donor-specific DEGs computed from real patient perturbation data (Fig. 6D, Methods). This setting allows us to assess whether Stack can identify which prompt cytokines and query contexts give rise to stronger donor-specific effects.

Across prompt cytokines, the proposed approach accurately recapitulates the measured trend in donor-specific effect sizes, as quantified by an adjusted DEG ratio index that corrects for the effects of total DEG counts and cell numbers (Figs. 6E, S24, Methods). Agreement between predicted and measured values is consistent across T cell subtypes with moderate subtype specificity (Fig. S25). On the query side, Stack recovers donor-associated differences across all disease contexts represented in the query samples, with smaller but still significant correlations and stronger cell-type specificity than in the cytokine analysis (Figs. 6F, S26). This association across query contexts also holds when each cytokine is evaluated independently (Fig. S27). Donor-specific effects show weak concordance within disease groups in both real measurements and Stack predictions, suggesting that inter-donor heterogeneity is not well covered by disease identity (Fig. S28). Across all analyses, Stack with synthetic prompts outperforms alternative baselines and Stack with original prompts in capturing donor-specific effects with subtype specificity (Fig. S29).

In Cell-Eval analyses, Stack achieves improved precision- and effect-size-based performance in recovering donor-specific DEGs, with larger gains for cytokines that elicit stronger donor-specific responses (Fig. S30A-B). Nevertheless, absolute scores remain low, likely reflecting substantial noise in the measured donor-specific DEG set, consistent with the minimal differences observed across methods under standard Cell-Eval applied to all DEGs (Fig. S30C). To further assess the validity of Stack predictions at the gene level, we compared Stack-predicted donor-specific DEGs with two eQTL reference sets, a CD4 T cell stimulation eQTL set (Soskic et al., 2022) and baseline CD4 and CD8 T cell eQTL sets from OneK1K (Yazar et al., 2022). Relative to shared DEGs, Stack-predicted donor-specific DEGs are significantly enriched for stimulation eQTLs, with statistics closely recapitulating those in the measured data (Figs. 6G, S31A). By contrast, Stack shows no enrichment for baseline OneK1K eQTLs across subpopulations, mirroring real measurements and supporting the specificity of the identified signals for perturbation effects (Fig. S31B).

To examine the biological basis of Stack donor-specific predictions, we systematically evaluated the association between donor-specific effect sizes and patient baseline expression levels across 11 curated gene modules (Table S5). The resulting association map reveals coordinated gene module patterns associated with donor-specific effects, which vary across cytokine perturbations and cell subtypes (Figs. S32, 6H). Strong associations in effector CD8 T cells include positive IL2-STAT5 associations for IFN-*𝛾* and IL-1*𝛽*, ISG associations for IFN-*𝛾* and TNF-*𝛼*, and TNF-NF*𝜅*B associations for TNF-*𝛼* and IL-12, consistent with prior knowledge (Schneider et al., 2014; Kalia and Sarkar, 2018). Predicted associations demonstrate near-perfect alignment with measured associations across all T cell subtypes (Table S6), supporting the use of Stack predictions for *de novo* biological discovery. A complementary feature selection approach confirms that gene modules jointly contribute to predicted donor-specific effect sizes, accounting for collinearity and highlighting individually non-significant modules (Methods). Through this approach, we found that gene module contributions to donor-specific effects are more consistent across cytokines in naive CD4 T cells, but become increasingly entangled and cytokine-specific in memory CD4 and effector CD8 T cells (Fig. S33). For instance, IL-15 in effector CD8 T cells shows a positive association with a pathogenic type 17 program but negative associations with ISG and TNF–NF*𝜅*B, contrasting with these modules’ effects on inflammatory cytokines, as corroborated by both association and feature selection analyses (Fig. 6H). This suggests that a type 17-skewed cellular state may redirect the IL-15 response program (Ding et al., 2020), whereas elevated interferon/STAT1 tone may constrain this divergence. Overall, our findings show that Stack can accurately identify cytokines and patients with donor-specific perturbation effects and link the predictions to biological mechanisms.

## 3. Discussion

As large atlases of single-cell transcriptomic profiles are compiled across tissues, species, and diseases, foundation models present an exciting opportunity to learn universal biological principles and patterns that generalize beyond experimentally observed data (Bommasani et al., 2021). Virtual simulations of cell states could significantly expand our understanding of cell biology through uncovering cellular states that are difficult to probe experimentally but can be inferred through relationships learned from existing data (Bunne et al., 2024; Roohani et al., 2025). However, realizing this promise requires models that transfer robustly across conditions and tasks. Most existing models present several limitations: they often fail to generalize to previously unseen conditions, provide no benefit over methods that are specifically fine-tuned on those datasets, and cannot perform new tasks without being explicitly trained to do so.

Here, we introduce Stack, a single-cell foundation model that leverages information from the cellular context of each cell to create enhanced representations. This design enables Stack to consistently outperform models trained from scratch on each evaluation dataset, highlighting Stack’s ability to meaningfully leverage information acquired during pre-training. This dependence on context also enables a novel capability: engineering context to design cell state. Because context can be defined in many ways, such as an applied perturbation, a disease state, or a new donor, Stack supports inference-time learning of new tasks, including zero-shot generalization to new biological contexts and datasets.

Through this capability of in-context learning, Stack enables a new approach to single-cell modeling in which counterfactual cell states can be generated via prompting with only cells. Importantly, Stack extends existing perturbation modeling by removing the reliance on perturbation labels, cell type encodings, and test-sample-specific controls, enabling direct, label-free comparison of cell states to resolve context-dependent responses that transcend categorical annotations. These results position Stack as a generative cellular model capable of predicting unobserved gene expressions across cell types, perturbations, and novel donors, with potential to accelerate therapeutic and drug discovery cycles. As a concrete demonstration, we applied Stack to generate *Perturb Sapiens*, a resource we believe will be of broad value to the community. Beyond atlas construction, we further demonstrated that Stack can zero-shot prioritize patient-specific cytokine responses given only patient baseline profiles and cytokine response measurements from healthy donors, opening new opportunities for personalized medicine.

While Stack shows strong performance, it also has several limitations that present opportunities for future research. The current model is trained exclusively on human single-cell data. Extending this framework to multi-species applications would require additional design in the tokenization procedure to account for gene misalignment across species. Model calibration for rare cell types and weak perturbation effects remains to be established, marking an important area for future development. Signaling cascades may introduce time-dependent or secondary perturbation effects, which may complicate interpretation of Stack predictions. As revealed by the donor-specific response prediction analysis, Stack can extract information from synthetic prompts that perform arithmetic on gene expressions, while the underlying mechanisms remain to be further elucidated. Finally, as the model is post-trained primarily on *in vivo* cell types, especially immune cells, a more sophisticated data curation and alignment scheme may improve model generalization across *in vitro* perturbation studies and *in vivo* observational data.

## 4. Methods

### 4.1. Stack model

#### 4.1.1. Generative process

Stack models the state of cell *𝑘*, **E**^(^*^𝑘^*^)^ ∈ ℝ*^𝑛^*^×^*^𝑑^*, as an ensemble of *𝑛* token vectors 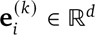, which we term *gene module* tokens, since each token represents a coherent subset of gene variability in the single cell:

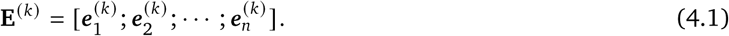

The flattened state vector **Ē**^(*𝑘*)^ ∈ ℝ*^𝑛𝑑^* is linked to the ground truth gene expression ***𝒙*** ^(^*^𝑘^*) ∈ ℝ*^𝐺^* with the generative process considered in scVI models (Lopez et al., 2018; Gayoso et al., 2022). The generative process involves a latent decoding transformation *𝑓* and cell library size scalar *𝑙* ^(^*^𝑘^*) ∈ ℝ. The final expression count is modeled through a negative binomial (NB) distribution with mean and dispersion parameters (***𝝆***^(^*^𝑘^*^)^, ***𝜽***^(^*^𝑘^*^)^):

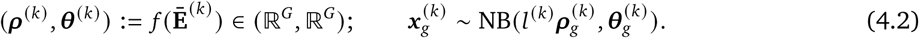

This generative process may be seen as a hybrid of transformer-based data modeling and biophysical-like models considered in scVI. The most prevalent differences from both strategies are summarized as follows.

- Stack does not consider low-dimensional latent variables as in scVI models; the total cell token dimensionality *𝑛𝑑* ∼ 10^3^, which is comparable to large language models and large-scale single-cell self-supervised-learning models (Cui et al., 2024; Rosen et al., 2023; Adduri et al., 2025).
- Stack does not include structural tokens (e.g. CLS tokens) as in several classic (single-cell) transformer models (Devlin et al., 2019; He et al., 2022; Rosen et al., 2023; Adduri et al., 2025). For downstream fine-tuning or linear probing on embeddings, the concatenation of all output tokens are used as the Stack model embedding output.

The generative process corresponds to the model decoding procedure from cell state embedding to gene expression, operating independently across cells.

#### 4.1.2. Stack architecture

During pre-training, the Stack model receives a *cell set* ***𝑿*** ∈ ℝ*^𝐾^*^×^*^𝐺^*, where *𝐾* denotes the number of cells and *𝐺* the total number of genes. The cell set includes cells continuously indexed from the same SRX experiment (for scBaseCount (Youngblut et al., 2025)) or the same dataset (for CELLxGENE (Program et al., 2025), where cells in datasets are primarily sorted by donor ID). The organization of each cell set introduces dependency across cells within the cell set, which serves as rich auxiliary information. Stack encodes the cell set ***𝑿*** ∈ ℝ*^𝐾^*^×^*^𝐺^* with the following architecture:

- **Tokenization.** First, each cell *𝑘* in the cell set ***𝒙*** ^(^*^𝑘^*) ∈ ℝ*^𝐺^* is projected into dimension *𝑛* × *𝑑* with a 1-layer perceptron, where *𝑛* is the number of gene module tokens and *𝑑* denotes the token size. Then a gene token embedding ***𝑷*** ∈ ℝ*^𝑛^*^×^*^𝑑^* is added on the perceptron output. This results in a tensor **Z**_0_ ∈ ℝ*^𝐾^*^×^*^𝑛^*^×^*^𝑑^* for the entire cell set.
- **Tabular transformer layer.** The tensors are processed by a stack of *𝑁_𝐿_* tabular transformer layers 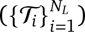 :

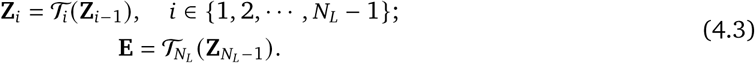 Each layer T*_𝑖_* applies a dual attention mechanism on the cell-set-level representations. In the following description, each attention module is a standard multi-head attention (MHA) block: the input is projected into multiple heads, attention outputs are concatenated and projected back, then combined with the input through a residual connection followed by layer normalization.

1. **Intra-cellular attention.** An intra-cell MHA module operates across the *𝑛* tokens independently for each cell in **Z***_𝑖_* ∈ ℝ*^𝐾^*^×^*^𝑛^*^×^*^𝑑^*. The sequence length is *𝑛*, and the feature dimension is *𝑑*. The attention head number is 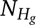 .
2. **Inter-cellular attention.** An inter-cell MHA module operates across *𝐾* cells, with the (*𝑛, 𝑑*) dimensions flattened into *𝑛𝑑*. The sequence length is *𝐾*, and the feature dimension is *𝑛𝑑*. The attention head number is 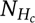.
3. **Feedforward network (FFN).** A position-wise FFN independently processes each *𝑑*-dimensional token, followed by residual connection and layer normalization, yielding **Z***_𝑖_* ∈ ℝ*^𝐾^*^×^*^𝑛^*^×^*^𝑑^*.
- **Cell-wise decoder.** Finally, the cell state embedding **E** ∈ ℝ*^𝐾^*^×^*^𝑛^*^×^*^𝑑^*is decoded into gene expression space independently for each cell in {1, 2, · · · *, 𝐾*}. The decoder *𝑓* is implemented as a multi-layer perceptron (MLP) that maps the flattened embedding **Ē**^(^*^𝑘^*^)^ ∈ ℝ*^𝑛𝑑^* to the negative binomial parameters (***𝝆***^(^*^𝑘^*^)^, ***𝜽***^(^*^𝑘^*^)^) ∈ (ℝ*^𝐺^,* ℝ*^𝐺^*), following the generative process described above.

#### 4.1.3. Pre-training objective

The model is pre-trained on a masked gene reconstruction task. For each input cell set ***𝑿***, a random subset of genes M ⊂ {1, 2, · · · *, 𝐺*} is selected, and their expression values are masked across all *𝐾* cells. The masking ratio is randomly sampled from (*𝑝*_min_*, 𝑝*_max_) = (0.1, 0.8) for each mini-batch. This selection range aims to cover gene dependencies of various strengths, following R^2^MAE (Dong et al., 2025). We adopt a higher *𝑝*_max_ (0.8) than in (Dong et al., 2025) (0.5) as the inter-cellular information brings additional information for implicit denoising. The pre-training objective combines a reconstruction loss with a latent space regularization term:

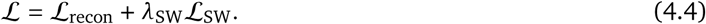

The reconstruction loss, L_recon_, is the negative log-likelihood of the original counts under the predicted NB distribution, computed for the masked genes across all cells in the cell set ***𝑿***:

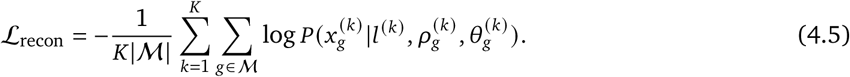

Optimizing solely the reconstruction loss leads to memorization and a reduction in performance. To enforce meaningful pre-training, we incorporate a regularization term, L_SW_, defined as the Sliced Wasserstein distance between the empirical distribution of the final flattened cell states 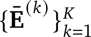 and a batch-centered, multivariate Gaussian prior 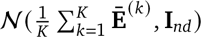 . This term encourages the embedding to decompose into a centered Gaussian component and a cell-set-specific constant vector, thereby regularizing the embedding distribution, motivated by linear identifiability results from (Dong et al., 2024; Khemakhem et al., 2020). The loss term is calculated within a random cell set subset, with subset size uniformly sampled from [32, 33, · · ·, 128]. The hyperparameter *𝜆*_SW_ (default 0.01) balances the two loss components.

### 4.2. Post-training

The pre-trained Stack model is post-trained for in-context prompting tasks through a supervised procedure. The objective is to empower the model to generate novel cell populations, combining cell type information from a set of *query* cells, and the biological context provided by a separate set of *prompt* cells. Our approach comprises four key components: (i) a novel self-distillation procedure, (ii) a training input construction strategy, (iii) architectural modifications, and (iv) a composite loss function that balances generalization with knowledge retention.

#### 4.2.1. Self-distillation

We introduce a self-distillation procedure during post-training. A frozen teacher model, operating without input data masking, computes embeddings and gene expression parameters from cell sets containing individual biological samples; these outputs are then used to calculate the training objectives. The teacher model weights are updated via exponential moving average (EMA) of the student model which is actively post-trained. This approach follows the self-distillation framework established in self-supervised vision models (Grill et al., 2020; Oquab et al., 2023; Zhou et al., 2021).

#### 4.2.2. Post-training input construction

Different from pre-training, the post-training process involves two biological samples with matched cell type annotations. We begin with a cell set of *𝐾* cells 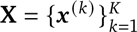 **X** = {***𝒙*** ^(^*^𝑘^*) }*^𝐾^*drawn from the prompt sample. Cells are ordered such that cells of the same type appear consecutively, with cell type order randomized for each sample. This cell set is partitioned into three components:

1. **Prompt condition cells** 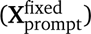 : The first 25% of cells, used unchanged as conditioning context.
2. **Prompt context cells** 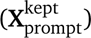: *𝐾*_kept_ cells also retained unchanged, analogous to unmasked tokens in diffusion language models that provide auxiliary context for reconstruction.
3. **Target cell positions**: The remaining *𝐾*_query_ positions, where *𝐾*_kept_ + *𝐾*_query_ = 0.75*𝐾* and the ratio *𝐾*_kept_/(*𝐾*_kept_ + *𝐾*_query_) ∼ U(0, 1).

The target cell positions are filled with *query cells* (**X**_query_) drawn from a different biological context (e.g., a different donor or perturbation condition). Each query cell is matched by cell type to the target cell it replaces. This yields the final input:

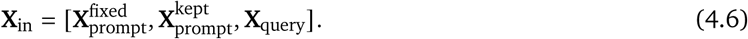

The model’s objective is to predict the gene expression distribution of target cells, conditioning on the prompt cells as reference. To ensure the model learns meaningful biological transitions, each replaced cell must have the same cellular identity (e.g., cell type or cell line, specified by user) as the original cell it replaces. To address imbalanced cell type distributions, we apply a balancing procedure to each training sample: overrepresented cell types are downsampled to the average count per type, then the resulting pool is upsampled with replacement to restore the original cell set size. For replicated cells, only the first instance is used in the distributional loss terms; further details are provided in Section 4.2.4.

#### 4.2.3. Architectural modifications

We introduce two learnable modules to the model: a query position embedding, **P**_query_ ∈ ℝ*^𝑛𝑑^* and an MLP binary classifier *𝑓*_CLS_ : ℝ^2^*^𝑛𝑑^* → ℝ. The position embedding **P**_query_ is added to the token representations of query cells at the first embedding layer. The MLP binary classifier *𝑓*_CLS_ : ℝ^2^*^𝑛𝑑^* → ℝ that takes the concatenation of mean prompt condition embedding and prompt context/query single-cell embedding as input, and predicts whether the cell comes from prompt (0) or query (1). The classifier receives detached embeddings as input, therefore its optimization is independent of other model weights. We register gradient hooks on these newly introduced modules to apply a 10× gradient scaling, which effectively increases their learning rates relative to the pre-trained parameters. Additionally, a causal attention mask is applied within all transformer layers to prevent prompt condition cells from attending to prompt context or query cells, ensuring that information flows strictly from prompt condition cells to the remaining cells.

#### 4.2.4. Post-training objective

Similar to the pre-training setup, the model receives a masked version of the input cell set **X**_in_ with rectangular gene masks, with masking ratio sampled from U(0.1, 0.3). The post-training objective is designed to simultaneously predict unseen target cells and maintain information learned during pre-training:

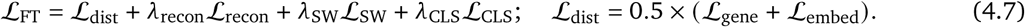

- **Embedding Alignment Loss (**L**_embed_):** The term is defined as the energy distance between Stack embeddings of query cells and target cells. The embedding for the target cells is extracted from the teacher model and is detached for loss calculation. The student model to be fine-tuned calculates the embedding for query cells from input data cell set **X**_in_.
- **Expression Alignment Loss (**L**_gene_):** The term is defined as the energy distance between the predicted and true distributions of log normalized target gene expression. To make the optimization on NB distribution parameters tractable, predictions are generated using a reparameterizable zero-inflated normal distribution sampler that matches first two moments of the log-normalized NB distribution. To address over-smoothing, we estimate a shared over-dispersion parameter for query cells using the median detached over-dispersion computed from prompt cells. The loss is computed on first 1,000 highly variable genes per mini-batch (identified via Pearson residuals (Lause et al., 2021)), and is stratified by cell type like L_embed_.
- **Distributional Alignment Loss (**L**_dist_):** This is the primary alignment objective, which is defined as the average of embedding alignment loss L_embed_ and gene expression alignment loss L_gene_.
- **Reconstruction Loss (**L**_recon_):** To retain the model’s mask reconstruction capabilities, we apply a standard masked gene reconstruction loss, as in pre-training, to the prompt cells. This serves as an auxiliary task that regularizes the model and enforces it to maintain pre-training objective. We use *𝜆*_recon_ = 1.
- **Latent Regularization (**L**_SW_):** The Sliced Wasserstein distance objective from pre-training is retained and applied to the embeddings of all cells in the batch. We use *𝜆*_SW_ = 0.01.
- **Classification Loss (**L**_CLS_):** The classification loss is defined as the binary cross entropy loss (BCE) of the MLP classifier in classifying query from prompt context cells. We use *𝜆*_CLS_ = 1.

#### 4.2.5. Generative procedure

The post-training setup closely resembles a conditional mask diffusion model, where prompt cells represent unmasked tokens and query cell represent masked tokens [mask] that the model must predict. Two key differences distinguish Stack’s generative inference:

1. The gene expression profile of query cells is input to the model in order to encode cell type information. Combined with **P**_query_, it serves a similar role to positional encodings of [mask] tokens in language models.
2. Masked language models derive token-level confidence directly from softmax probabilities over the vocabulary. Since our output space is continuous, we instead train a separate classifier to estimate prediction confidence, which guides the selective unmasking procedure during generation.

At each generative step, the model receives mini-batches containing concatenated prompt condition cells, prompt context cells and query cells. Here, prompt context cells correspond to original cell expression profiles. The ratio of prompt condition cells remains constant at 25%, while the ratio of prompt context cells increases linearly from 0.2 to 0.4 throughout the generative process. A boolean array is_mask is maintained to indicate whether each query position remains to be predicted, and is initialized with all True for query cells. Each step consists of a prediction-and-update cycle guided by a linear masking schedule (1 − *𝑡*/*𝑇*):

- **Prediction:** The model performs a forward pass, generating a complete expression profile for all query cells.
- **Confidence Scoring:** The classifier module assesses each predicted query cell, and outputs a logit vector. Positive values indicate the cell is more like a query cell, negative values indicate the cell is more like a prompt cell.
- **Selective Unmasking:** Based on the masking schedule, a fraction of query cells with is_mask=True are selected to be replaced with model prediction. The cells with the lowest logit values are chosen for the replacement.
- **State Update:** The replaced cells’ is_mask values are set to False. Next, all query cells with logit value *>* 0 (i.e., those classified as query cells) are (re-)set to is_mask=True.

The iterative process concludes when the masking rate in the schedule reaches zero. The final output from this step constitutes the complete, generated expression matrix for the initial set of query cells. The final logit vector is also a part of the output for quantifying generation quality and interpretation.

### 4.3. Model implementation

The Stack model is implemented in PyTorch and trained using the PyTorch Lightning framework. The training process consists of two stages: self-supervised pre-training and supervised post-training for in-context prompting.

#### 4.3.1. Base model

The Stack architecture consists of *𝑁_𝐿_* ∈ {3, 6, 9} tabular transformer layers. Each cell is tokenized into *𝑛* = 100 tokens, each with dimension *𝑑* ∈ {8, 16, 32}, yielding a total per-cell embedding dimension *𝑛𝑑* ∈ {800, 1600, 3200}. The input to the transformer is a cell set of *𝐾* = 256 cells tokenized. The feed-forward network within each layer uses a GELU activation function. The decoder is implemented as a 2-layer MLP with GELU activation. No dropout was applied during pre-training. Configurations of different model settings are detailed in the scaling study section.

#### 4.3.2. Pre-training data

The pre-training data was sourced from the full human scBaseCount (Release 2026-01-12), the scBaseCount subset, or the human CELLxGENE. For scBaseCount, we filtered cells with 300–7,000 detected genes and at least 700 UMIs. No filtering was performed for CELLxGENE datasets. For model training, a unified gene list was created by computing the union of the top 1,000 highly variable genes (HVGs) from each scBaseCount data file, capped at a maximum of 15,012 total genes. HVGs were identified using analytic Pearson residuals (Lause et al., 2021). The dataloader generates samples by creating non-overlapping, contiguous chunks of *𝐾* = 256 cells from each file. Cells in chunks shorter than *𝐾* are dropped. Pre-training data metrics are summarized in Table 1.

**Table 1.** An overview of training datasets for Stack models in this study.

| Training sets | #Training Datasets | #Training Cells |
| --- | --- | --- |
| CELLxGENE | 905 | 73.7M |
| scBaseCount subset | 9004 | 60.2M |
| scBaseCount full set | 19978 | 148.8M |

**Table 2.** An overview of model settings tested in the scaling study. Symbolsindicate the training data used for each model variant (Table 1): § CELLxGENE, † scBaseCount subset, ‡ scBaseCount full set.

| Model Variant | Layers $N_L$ | Total Embed Dim $nd$ | Params | Non-emb Params |
| --- | --- | --- | --- | --- |
| STACK (Base-) † | 3 | 800 | 69.1M | 57.0M |
| STACK (Base) §, †, ‡ | 6 | 800 | 76.7M | 64.7M |
| STACK (Base+) † | 9 | 800 | 84.4M | 72.4M |
| STACK (Medium) † | 6 | 1600 | 186M | 163M |
| STACK (Large) †, ‡ | 9 | 1600 | 217M | 193M |
| STACK (XLarge) ‡ | 6 | 3200 | 506M | 459M |
| STACK (Huge) ‡ | 9 | 3200 | 629M | 582M |

**Table 3.**
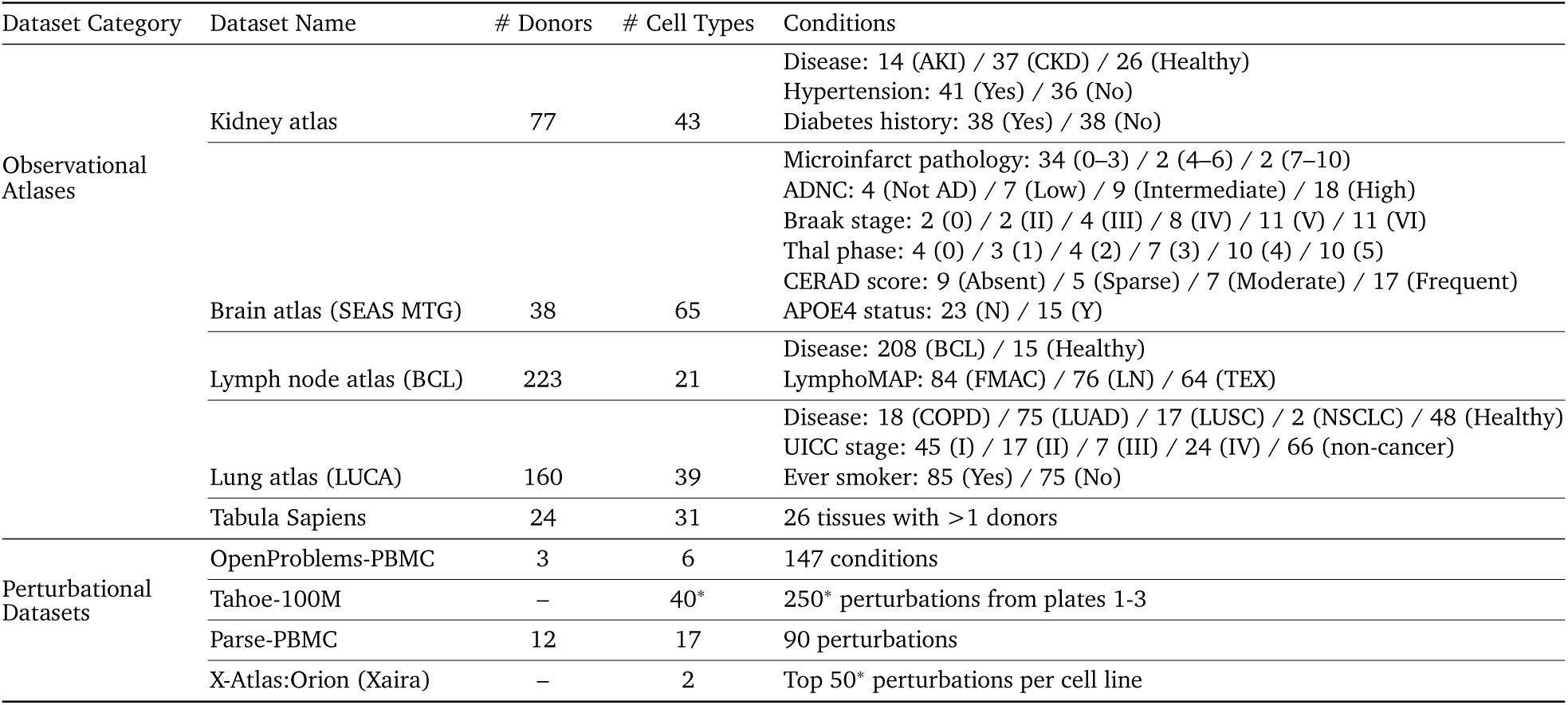
Overview of datasets used in this study. ^∗^ indicates subsampling from the original datase.

**Table 4.**
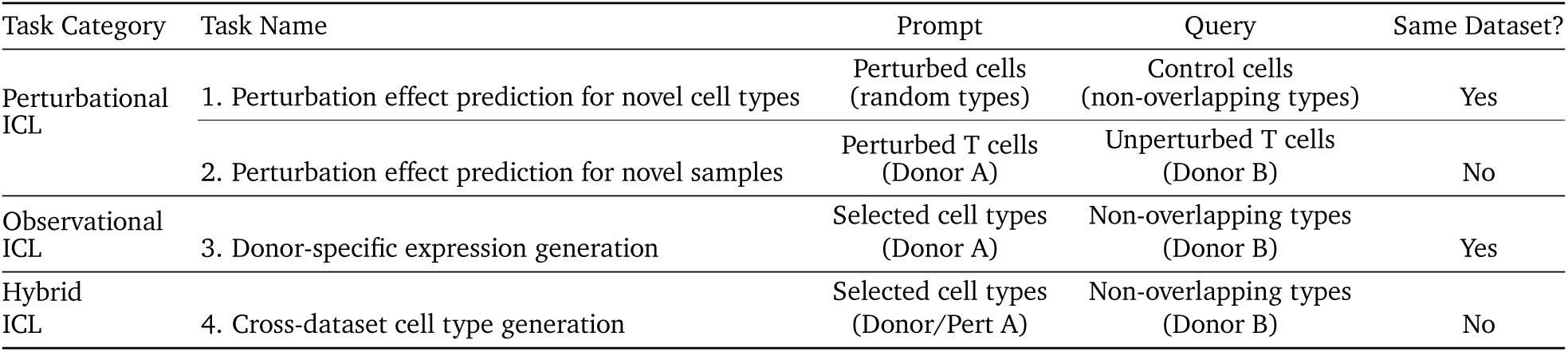
Overview of in-context cell prompting tasks for Stack>.

#### 4.3.3. Pre-training setup

The Stack base/large model was pre-trained for 10 epochs using the AdamW optimizer with a peak learning rate of 1 × 10^−4^ and a weight decay of 3 × 10^−3^. For Stack XLarge and Huge models, the peak learning rate was tuned down to 3 × 10^−5^ to stabilize training. A cosine annealing learning rate schedule was used with a linear warmup over the first epoch. The Sliced Wasserstein regularization weight *𝜆*_SW_ was set to 0.01. All experiments were conducted on a single NVIDIA H100 GPU (80GB HBM) with 320GB system RAM, using bf16 mixed precision with a batch size of 32 and 4 training data-loading workers. We benchmarked data-loading and pre-training speed against a State Embedding model under equivalent settings.

#### 4.3.4. Post-training data

The dataset for supervised alignment was curated from multiple public sources, including the Parse 10M PBMC data (Parse Biosciences, 2023), and a programmatically selected subset of the CELLxGENE database (Program et al., 2025). To ensure suitability for learning donor-specific effects, a CELLxGENE subset was filtered to include only large-scale datasets (*>* 50, 000 cells) with at least five unique donors. As the default “cell_type” column in CELLxGENE is found to be suboptimal, we implemented an automatic procedure to identify optimal cell type annotation column for each dataset, employing a heuristic algorithm that prioritizes author-provided, intermediate-granularity labels (e.g., ‘author_cell_type’, ‘ann_coarse’) over standardized ontologies or overly detailed subtypes. The dataset selection procedure resulted in 45 million cells from 189 datasets. The post-training dataloader first splits the curated datasets into training, validation, and test sets based on donor or sample ID. Each training sample consists of a cell set of *𝐾* = 512 cells, which is further partitioned into *𝐾*_fixed_ = 128 prompt condition cells and *𝐾*_kept_ + *𝐾*_query_ = 384 prompt context/target cells.

#### 4.3.5. Post-training setup

The model was fine-tuned for 8 epochs, starting from the pre-trained weights from the Stack (large) model trained on the full human scBaseCount. We used the AdamW optimizer with a peak learning rate of 2 × 10^−5^ and a weight decay of 3 × 10^−3^. A cosine annealing learning rate schedule with a 1-epoch linear warmup and min learning rate 5 × 10^−6^ was applied. For teacher model updates, we employed an exponential moving average (EMA) with a decay rate of 0.95, applied every 500 optimization steps. The training was configured with a batch size of 8 and 4 steps of gradient accumulation, resulting in an effective batch size of 32. Fine-tuning experiments were conducted on a single NVIDIA H100 GPU (80GB HBM) with 400GB system RAM, using bf16 mixed precision.

### 4.4. Ablation and scaling studies

We performed comprehensive ablation and scaling studies to evaluate the performance of Stack across settings and scales. An overview of tested models is shown below. All models use *𝑛* = 100 gene module tokens and attention heads *𝑁_𝐻𝐶_* = *𝑁_𝐻𝐺_* = 8, except for XLarge and Huge models which use *𝑁_𝐻𝐶_* = 20.

We also evaluated two ablations of Stack (Base) †: (i) without latent regularization, and (ii) without both latent regularization and inter-cellular attention.

### 4.5. Stack embedding evaluations

#### 4.5.1. Probing evaluation

To assess the biological information encoded in the learned cell representations, we implemented a multi-tiered probing framework. All probing experiments employed a group-based splitting strategy, where datasets were partitioned by donor ID for observational data and PBMC perturbation data, or by 50% sample split (grouped by library label) for cell-line perturbational data. This ensures that models are evaluated on cells from entirely unseen donors (or library splits for cell-line datasets). To address class imbalance in the test set, we capped the maximum contribution per donor at 2,000 cells for observational data, 25,000 cells for Parse, and 100,000 cells per sample split for other perturbation datasets. Two main probing approaches were employed to evaluate different aspects of the learned representations:

- **Linear probing:** Logistic regression for classification tasks and ridge regression for continuous variables, trained with 5-fold cross-validation on 80% of donors. This setting trains separate models for each cell type, enabling assessment of cell-type-specific information encoding. All analyses were performed on the top 20 most abundant cell types if the total cell type number exceeds 20 (or 5 per tissue in Tabula Sapiens).
- **MLP probe:** For capturing non-linear relationships, we implemented an MLP consisting of an input layer normalization, a hidden layer with 128 units and ReLU activation, dropout of 0.2, and a task-specific output layer. The model was optimized using AdamW with learning rate 1 × 10^−3^ for up to 80 epochs with early stopping (patience of 12 epochs). L2 regularization strength was selected via grid search over {0, 10^−5^, 10^−4^, 10^−3^, 10^−2^, 10^−1^} based on validation loss, using a 70/15/15 train/validation/test split by donor. The model is trained on 5 most abundant cell types together for each dataset.

Performance was evaluated at the single-cell level. Since the test set caps each donor’s contribution, the calculated cell-level accuracy metrics equal donor-level average metrics when all donors exceed the 2,000-cell threshold, and smoothly approximate it when some donors have fewer cells. For classification tasks, we report balanced accuracy. For regression tasks, we report Pearson correlation coefficients. All evaluations were performed with a fixed random seed of 42.

#### 4.5.2. Batch integration evaluation

We performed batch integration evaluations using the scib-metrics package (v0.5.6). To remove extremely rare cell populations that lead to benchmarking errors, the analysis was conducted on a subset containing the top 20 most abundant cell types, or the top 5 most abundant cell types for Tabula Sapiens. Throughout our study, we use the ‘broad cell class’ annotations provided in Tabula Sapiens to designate cell type, coarsening to the cell class level where appropriate. If the resulting cell count exceeded 100,000, the subset was randomly downsampled to this size. The evaluation was structured around two primary objectives, each assessed by a specific suite of metrics.

1. **Batch Correction:** To quantify the removal of technical batch effects, we measured the k-nearest neigbor Batch Effect Test (kBET), graph connectivity, principal component regression (PCR) score, integration Local Inverse Simpson’s Index (iLISI), and batch-removal-adapted silhouette (BRAS) (Rautenstrauch and Ohler, 2025).
2. **Biological Conservation:** To assess preservation of biological signal, we measured Normalized Mutual Information (NMI) and Adjusted Rand Index (ARI) of Leiden clusters against ground-truth cell type labels, silhouette score for cell type labels, and the cell type Local Inverse Simpson’s Index (cLISI).

These scores are aggregated to compute the total score, following the scib-metrics default (Luecken et al., 2022).

#### 4.5.3. Baseline models

The Stack embedding was benchmarked against several methods:

1. **PC HVG:** We performed library normalization, log1p transformation, and Scanpy (v1.11.2) default highly variable gene selection to identify the top 2,000 highly variable genes. These genes were then reduced to 800/50 principal components. The former setting, matching the dimensionality of Stack (base), excels at probing tasks, while the latter is better for integration evaluations.
2. **scGPT (v0.2.4):** We used the recommended whole-human scGPT checkpoint to generate cell embeddings. Input data was preprocessed with library size normalization and log1p transformation, with a batch size of 256 for inference.
3. **UCE (commit 8227a65):** We employed the 33-layer model checkpoint in (Rosen et al., 2023) with batch size 20 to extract embeddings. The model operates directly on raw count data without normalization. As UCE by default can skip an extremely small amount of cells due to preprocessing, we aligned the output embeddings back to the original cell indices, filling missing cells with zeros.
4. **State (SE) (state v0.9.27):** We used the SE-600M checkpoint on Hugging Face and the state emb transform command to infer embeddings on raw count data. Data loader throughput was measured using: uv run state emb fit model.batch_size=32 optimizer.gradient_accumulation_steps=256 dataset.num_train_workers=4.
5. **TranscriptFormer (v0.6.1):** We used the TF-sapiens checkpoint from (Pearce et al., 2025) due to its advantage on human scRNA-seq data evaluations. We used the provided CLI to infer embeddings from raw count data.
6. **scVI fine-tuned (scvi-tools v1.3.1):** Pre-trained scVI model initially trained on the scBaseCount subset for 10 epochs, with 2 hidden layers of 2000 hidden dimensions and 800 latent dimensions, or 256 hidden dimensions and 50 latent dimensions. The model was subsequently fine-tuned on each target dataset for 20 additional epochs with learning rate 5 × 10^−4^. The fine-tuning employed a streaming mini-batch strategy: samples of 256 cells were drawn from individual files and concatenated to form training batches of 4096 cells. Each cell was tagged with a batch ID derived from its source file. The strategy greatly accelerates scVI training on large single-cell data collections.
7. **scVI from scratch (scvi-tools v1.3.1):** Models were configured with 2 hidden layers of 2000/128 hidden dimensions and 800/30 latent dimensions, and trained for 50 epochs using Adam optimizer with learning rate 1 × 10^−3^ and weight decay 1 × 10^−4^.

#### 4.5.4. Evaluation datasets

The five observational studies for probing and integration evaluations are downloaded from the CELLxGENE portal (Program et al., 2025) (Kidney, BCL, SEAS-AD MTG, LUCA, and Tabula Sapiens). Perturbation datasets are downloaded from their official websites (Tahoe, Parse, X-Atlas:Orion (Xaira), and the OpenProblems competition). We balanced the tissue composition of Tabula Sapiens, downsampling cells in each tissue to 20,000 if the cell number exceeds the threshold.

### 4.6. Cell prompting task evaluation

The prompting evaluation framework assesses how well a model can generate desired expression profiles given prompt and query cells through in-context learning (ICL). We evaluated four ICL tasks:

#### 4.6.1. Evaluation data construction

In tasks with cell type hold-outs (1, 3, 4), a set of broad cell classes present in one donor/condition was randomly sampled and held out (ratio 0.75) as target cells to be predicted. The remaining non-held-out cell types from this donor formed the prompt data, while held-out cell types from another donor/condition were used as query cells. In response prediction across samples (task 2), all data were restricted to a single cell type (T cells). In perturbational data (tasks 1–2), control conditions from the same prompt donor are typically available. Therefore, we additionally utilized these cells as auxiliary prompts to the model and used the output as a “synthetic control” for cell-eval benchmarking. This synthetic control approach was applied to Stack and other baselines when appropriate. For observational prompting tasks, oracle baselines additionally used non-held-out cell types in query donors as auxiliary data, which were not available to Stack. We equalized the number of query, auxiliary, and target cells by downsampling to the minimum available cell count among all three groups. To account for rare cell types in tasks 1 and 3, we upsampled each cell type in the query data to a minimum of 2000 and 1000 cells, respectively.

#### 4.6.2. Baseline models

The prompting performance of Stack was benchmarked against several methods:

1. **Original Query:** The original query cells are used for prediction, representing a zero-change scenario where no information from the prompt is used.
2. **Nearest Cell Type in Prompt:** This baseline computes pseudobulk expression profiles for all cell types in the prompt data. For each query cell, it identifies the most similar cell type in the prompt based on pseudobulk Pearson correlation, then assigns an expression profile from a randomly sampled cell of that matched cell type without replacement. Once all cells from a cell type are exhausted, the second-closest cell type is used, and so on.
3. **Same Cell Type in Prompt:** This baseline assigns expression profiles by sampling from cells of the same cell type in the prompt without replacement when possible.
4. **PerturbMean/DonorMean:** This method computes the average difference in expression profiles (per cell type) between contexts. This difference vector is calculated on non-held-out cell types between prompt and auxiliary data, then added to the expression profiles of the query cells. The synthetic control is not applicable for this baseline, as it would be equivalent to the query data.
5. **scVI:** A 2-layer scVI model (scvi-tools v1.3.1) with 30 latent dimensions and 200 hidden dimensions was trained on combined prompt and auxiliary samples, using dataset origin as the batch key, for 50 epochs. The model was then used to project query cells into the latent space and sample their expression profiles, artificially specifying “prompt” as the batch label.
6. **State:** State models (v0.9.27) were trained using the ST+SE setting (Adduri et al., 2025), where the model predicts cell embeddings and simultaneously decodes them back to gene expression space. Models were trained with Maximum Mean Discrepancy (MMD) loss on prompt, query, and auxiliary data, using the union of highly variable genes in cell-eval benchmarks across random seeds. Only control cells from the query set were used for model training. All models used a hidden dimension of 328 and cell set length of 32. Models were trained for 60,000 steps (batch size 8, learning rate 3 × 10^−4^). Random basal mapping was employed with gene-space outputs. The best checkpoints were selected via validation loss on held out cells.

#### 4.6.3. Evaluation metrics

We evaluated the generated expression profiles against ground truth data using pseudobulk correlation and DE metrics. Both categories of metrics are implemented using cell-eval v0.6.6 (Adduri et al., 2025) with default parameters, employing Wilcoxon rank-sum tests for DE detection and Benjamini-Hochberg correction for multiple testing. All metrics were computed on top 2000 log-normalized highly variable genes, identified from the concatenation of target and query data. For task 4, we additionally evaluated the integration performance of ground truth and predicted gene expression profiles using scIB, with the same configuration as the earlier benchmarking.

- **Pseudobulk correlation.** We measured how well methods capture pseudo-bulk level perturbation effects by two metrics:

**– Pearson Delta**: Pearson correlation between predicted and observed expression changes. For each perturbation *𝑡*, we calculate the expression delta as Δ*_𝑡_* = | *𝑝_𝑡_* − *𝑝*_ctrl_ | for both predicted (Δ̄ *𝑡*) and ground truth (Δ*_𝑡_*) pseudobulks, then compute: Pearson-Δ = corr(Δ̄ *_𝑡_,* Δ*_𝑡_*).
**– DE Spearman LFC**: the Spearman rank correlation between predicted and observed log2-fold-changes, calculated within the set of significantly differentially expressed (DE) genes in the ground truth.
**– DE direction match**: This metric measures whether the predicted direction of gene expression change (i.e., up- or down-regulation) matches the ground truth. It is calculated only on the set of genes that are significantly differentially expressed (DE) in both the predicted and true data. The score is the fraction of these shared DE genes for which the direction of change matches. This metric replaces Pearson Delta in observational prompting tasks. Pearson Delta performs comparably to DE Spearman LFC when many genes are identified as DEGs, while also showing greater vulnerability to batch effects.
- **Differential expression accuracy.** Finally, we evaluated whether the prediction captures differential expression between ground truth data and the input query condition.

**– PR-AUC**: Area under precision-recall curve using binary DE labels and − log_10_(*𝑝*-values) as scores, using sklearn average precision score implementation.
**– Spearman effect size**: To compare the relative effect sizes of perturbations, we calculate Spearman correlation coefficients on the number of differentially expressed genes (adjusted p-value < 0.05) between predicted and ground truth. This assesses whether models accurately capture relative effect sizes across different conditions. We averaged predictions across random seeds within each cell type and perturbation/donor, yielding a Spearman correlation computed across perturbations/donors per cell type.
**– DE Overlap Accuracy**: Computes the overlap between the top-*𝑁* genes from the true DE abs-log-fold-change ranking and the top-*𝑁* genes from the predicted DE ranking, calculated as |top-*𝑁*true∩ top-*𝑁*predicted|/*𝑁*, where *𝑁* is the total number of true DE genes.
**– DE Precision-at-***𝑁*: Computes the overlap between the top-*𝑁* genes from the true DE ranking and the top-*𝑁* genes from the predicted DE ranking, calculated as |top-*𝑁* true ∩ top-*𝑁* predicted|/*𝑁*, where *𝑁* is the total number of predicted DE genes. This metric assesses the fidelity of predicted DE gene lists and replaces Spearman effect size correlation in Setting 2, where the limited number of perturbations with similar effect sizes makes rank-based correlation unreliable.
**– Jaccard similarity**: For each perturbation, we compute the Jaccard index between the set of predicted DE genes and ground truth DE genes, defined as the size of the intersection divided by the size of the union of the two sets.

#### 4.6.4. Evaluation datasets

For evaluations, we used 1. OpenProblems drug perturbation (Luecken et al., 2025), 2. Cytokine stimulation (Dong et al., 2023), 3. Immune aging (Wells et al., 2025), 4. Tabula Sapiens (The Tabula Sapiens Consortium et al., 2022), 5. Kidney atlas (De Boer et al., 2021), 6. Lymph node BCL (Li et al., 2025), 7. Liver atlas (Edgar et al., 2025), 8. Parse cytokine perturbation dataset (Parse Biosciences, 2023). Observational atlases (3-7) were downloaded from the CELLxGENE portal, and the cytokine stimulation data was downloaded from Dryad (Dong et al., 2023). The cytokine stimulation data (Dong et al., 2023) comprise 3 donors, but donor 1 has very limited cell numbers. Therefore, we restricted the dataset to include only donors 2 and 3 and only included cells from the acute condition (2 days).

For task 1, the OpenProblems dataset used prompts and queries sampled from the same donor (3 donors total) within each test experiment. For task 2, prompts used T cells from one Parse donor under cytokine conditions (IFN-*𝛽*, IFN-*𝛾*, IL-6, TNF-*𝛼*), while queries used control T cells from donors a or b in (Dong et al., 2023). Ground truth evaluation used corresponding individual and combination conditions (IFN-*𝛽*, IFN-*𝛾*, IL-6, TNF-*𝛼*, IFN-*𝛽*+IFN-*𝛾*, IFN-*𝛽*+IL-6, IFN-*𝛽*+TNF-*𝛼*) from (Dong et al., 2023). For combinatorial perturbation evaluations, we computed weighted sums of single-perturbation Stack predictions. Due to dosage differences between the Parse dataset and Dong et al. (2023), we performed a grid search over weights [0.1, 0.3, 0.5, 0.7, 0.9] to determine the optimal weighted average of normalized gene expression conditions in the original prompt T cells. The optimal weight was selected based on Pearson Delta and subsequently used to generate Stack predictions for combinatorial perturbations. For task 3, prompts and queries comprised non-overlapping cell types sampled from different donors in each dataset. For task 4, prompts comprised: Parse control cells (PBS condition) with one donor sampled per prompt; Parse donor 1 cells with one perturbation sampled per prompt; immune aging dataset cells with one donor per prompt; OpenProblems cells with one perturbation per prompt. All queries in this task used immune cells from Tabula Sapiens. In tasks 3 and 4, we performed cell class-level hold-out for each evaluated dataset, relabeling cell annotations from original author annotations to align with the broad cell classes defined in Tabula Sapiens for corresponding tissues. Holding out at the cell class level minimizes potential information leakage across similar cell types.

### 4.7. Whole-organism perturbational atlas *Perturb Sapiens*

In the analysis, we used each condition from donors 0-2 in OpenProblems drug perturbation data, donors 1-12 in Parse cytokine perturbation data, and donors 1-4 from primary human CD4 T cell Perturb-seq data as the prompt, and the tissue balanced version of Tabula Sapiens as the query. For drug perturbations, only conditions with more than 332 cells (512 × 0.65, the minimum needed to fill one prompt cell set at the final Stack generation step) were included, yielding 71 drugs for donor 0, 105 for donor 1, and 111 for donor 2, comprising 136 unique drugs in total. For genetic perturbations, we retained only conditions from unstimulated T cells that were originally annotated as having *>* 10 differentially expressed genes and on-target significant with more than 332 cells, yielding 135, 326, 267, and 93 qualifying perturbations for donors 1-4, respectively. Cells without sgRNA were concatenated and subsampled to 10,000 to serve as controls per donor. We used post-trained Stack (T=5) to perform in-context generation. We removed cells with classifier logits above 2.5.

We generated log2-fold-change heatmaps for individual cytokine, drug, and genetic perturbations. For each cell type, up to 10,000 cells per condition were subsampled, normalized, and log-transformed. Genes were restricted to 4,000 highly variable genes identified via Pearson residual in the example ADSF *Perturb Sapiens* dataset (Lause et al., 2021). Differential expression analysis was performed using a two-sided Wilcoxon rank-sum test with Benjamini-Hochberg correction (DEG: FDR *<* 0.05, |log_2_FC| *>* 0.5). Ribosomal protein genes (*Rps*, *Rpl*), mitochondrial genes (*mt-*, *Mt-*), and predicted genes (*Gm*) were excluded from the final results.

To aggregate IFN-*𝛾* DEG results across donors in Parse, we computed the median log2-fold-change for each gene. Per-donor *𝑝*-values were combined using an order-statistic approach: for *𝑘* donors, the *𝑟*-th smallest *𝑝*-value (*𝑟* = ⌊*𝑘*/2⌋ + 1) was evaluated against a Beta(*𝑟, 𝑘* + 1 − *𝑟*) distribution to obtain a combined *𝑝*-value. Genes for which fewer than 60% of donors agreed on the direction of effect were assigned as non-significant.

Only genes tested in at least half of the donors (and a minimum of 2) were retained. Combined *𝑝*-values were corrected for multiple testing using the Benjamini–Hochberg method. This procedure was applied to both *Perturb Sapiens* and real Parse PBMC profiles, with LFC thresholds of 0.5 and 0.375, respectively. For all remaining heatmaps without cross-donor aggregation, a uniform LFC threshold of 0.5 was used.

For each perturbation (drug/cytokine/genetic) and its matched control, we identified tissue–cell type combinations with sufficient cell counts (≥ 1,000 cells, capped to 10,000 max). We then applied the same differential expression analysis pipeline described above to compute LFCs and adjusted *𝑝*-values. Cell types were mapped to the following cell classes (Epithelial, Endothelial, Germ, T Cell, Innate Lymphoid, B Lymphoid, Myeloid, Stromal/Fibroblast, Contractile, Stem/Progenitor, Erythroid/Hematopoietic, Platelet/Megakaryocyte, Glial, Neuron, Other Lymphocyte, and Other) based on keyword matching of their original annotations. We applied a simplified procedure to aggregate LFCs and significance across donors and cell types in the same class: we took the median LFC and median FDR, and genes for which fewer than 60% of donors agreed on the direction of effect were assigned as non-significant. Perturbation effects are defined as log2-fold-changes, with non-significant values set to zero. To harmonize perturbation effects across modalities in *Perturb Sapiens*, we first clipped perturbation effects to the range (−10, 10), normalized each modality to a mean absolute effect of 0.01 across samples and genes, and then applied a second round of clipping to (−5, 5). We then performed dimensionality reduction by first computing 100 principal components on the concatenated perturbation effect matrix, followed by independent component analysis (ICA; *𝑘* = 50 components) on the mean-centered PCA scores using the FastICA algorithm with default parameters. Gene-level ICA loadings were recovered by projecting the PCA loading matrix onto the ICA mixing matrix. For visualization, we subsampled perturbations by hierarchically clustering their mean ICA profiles and selecting evenly spaced representatives from each cluster. To characterize the dominant source of variation captured by each independent component, we computed variance decomposition using nested ANOVA *𝜂*^2^ statistics across four factors: perturbation modality, perturbation identity (nested within modality), tissue, and cell class. Each component was classified by its factor with the largest *𝜂*^2^. To characterize pathway-level perturbation effects in a specific tissue context, we performed Gene Ontology Biological Process enrichment on significant up-regulated genes for each perturbation in large intestine epithelial cells using gseapy’s (v 1.1.11) Enrichr against the GO Biological Process 2025 gene set library (Fang et al., 2023; Kuleshov et al., 2016; Aleksander et al., 2025).

### 4.8. Evaluations of *Perturb Sapiens*

For evaluation on non-immune cells in cytokine *Perturb Sapiens*, we downloaded three epithelial *in vitro* cytokine perturbation datasets from the GEO database (Koh et al., 2023; Swindell et al., 2018; Lee et al., 2022). All evaluations were restricted to the top 4,000 highly variable genes identified from the example ADSF *Perturb Sapiens*. Log2-fold-changes, *𝑝*-values, and adjusted *𝑝*-values were computed using PyDESeq2 (v0.5.2) on either pseudobulk (Koh et al., 2023) or bulk RNA-seq data (Swindell et al., 2018; Lee et al., 2022). Differential expression analysis on *Perturb Sapiens* was performed using Scanpy’s Wilcoxon rank-sum test implementation, with the same aggregation procedure as in the heatmap analysis. Airway and keratinocyte epithelial populations in *Perturb Sapiens* were constructed by selecting cell types annotated as “epithelial” from their respective tissues (lung, trachea for airway; skin and tongue for keratinocytes). To ensure fair comparison across *Perturb Sapiens* cell types, each cell type population was capped at a maximum of 15,000 cells. We applied an LFC threshold of 0.25 and a FDR threshold of 0.05 for cell-eval evaluations.

We define the specificity used for *Perturb Sapiens* evaluation as follows. For each cell-eval metric, we construct a matrix where entry (*𝑖, 𝑗*) corresponds to the metric value between prediction *𝑖* and ground truth perturbation *𝑗*. Specificity is then defined as the normalized rank percentile of a predicted perturbation’s self-match performance (diagonal entry) relative to all other predictions evaluated against the same ground truth, computed as (*𝑟* −1)/(*𝑛*−1) where *𝑟* is the rank and *𝑛* is the number of valid entries. This is related to but distinct from the Perturbation Discrimination Score (PDS) in cell-eval (Adduri et al., 2025), which uses L1 distance and instead ranks over ground truth perturbations for each prediction. We alter the evaluation metric for two reasons. First, the original PDS evaluates specificity only on L1 distance, a single pseudobulk metric over all available genes; here we instead assess specificity across all cell-eval metrics for more thorough evaluations. Second, a number of cell-eval metrics are defined with respect to ground-truth differentially expressed genes, so their absolute values are not directly comparable across ground truth perturbations, and ranking over them introduces artifacts. Our approach addresses both issues and systematically evaluates model prediction specificity. Here to ensure valid specificity comparisons, we used a common gene set across all (*𝑖, 𝑗*) pairs, rather than reusing the cell-eval benchmarking results, which identify a separate HVG list for each condition.

To benchmark cytokine and drug perturbations in *Perturb Sapiens* against biological replicates, we performed differential expression (DE) analysis across all cytokine perturbations as well as 14 drugs (shared in donor 1 and 2) with all cell types available. For each perturbation, 2000 highly variable genes were identified from the corresponding treated and control cells across all cell types in the real data. We compared DEG results across cell-type pairs in biological replicate vs. real data (3 × 3 cell-type pairs: T cell, B cell, Myeloid), and *Perturb Sapiens*-generated vs. real data (4 × 3 pairs, additionally including epithelial cells in *Perturb Sapiens*). Within each condition and cell type, cells were randomly subsampled to a maximum of 20,000 per group. DE testing followed the same Wilcoxon rank-sum procedure described above, with significant genes defined by adjusted *𝑝*-value ≤ 0.05 and | log_2_ FC| ≥ 0.25. Cell-eval evaluation metrics were constructed as described in the previous section with one modification: significant genes were defined using an explicit minimum log2-fold-change threshold, consistent with the DEG procedure. For cross–cell-type heatmaps, each per-cytokine metric matrix was inverse-rank normalized within each real (Parse) cell type, then averaged across cytokines to equalize cytokine contributions.

To benchmark genetic perturbations, we used the Replogle RPE1 dataset as ground truth, downloaded from https://plus.figshare.com/ndownloader/files/35775606. We observed that this dataset contains global stress signatures spanning perturbation conditions, which confound downstream comparisons (Fig. S20), consistent with findings reported in (Viñas Torné et al., 2025). To mitigate this confound, we constructed a synthetic control by subsampling 50 cells from each perturbation condition, concatenating all conditions, and drawing a final subsample of 20,000 cells. Because the RPE1 dataset contains fewer genes, we restricted the analysis to genes shared across RPE1, the prompt set, and *Perturb Sapiens*, selecting 2,000 highly variable genes from RPE1 for each evaluation within this common gene set. All remaining DEG-calling procedures and metric definitions were the same as in the previous paragraph.

### 4.9. DiseasePert-3M

#### 4.9.1. Donor Sample Procurement and Selection

A total of 40 cryopreserved primary human T cell samples isolated from PBMCs were procured from 40 unique donors through StemCell Technologies, spanning healthy controls (n = 8), cancer patients (n = 16), and patients with autoimmune/inflammatory disorders (n = 16). Cancer donors included colorectal cancer (n= 4), pancreatic cancer (n = 2), liver cancer (n = 4), MCL (n = 1), follicular lymphoma (n = 1), prostate cancer (n = 2), and CML (n = 2). Autoimmune/inflammatory donors included ulcer colitis (n = 3), type I diabetes (n = 3), lupus (n = 2), multiple sclerosis (n = 2), celiac disease (n = 2), rheumatoid arthritis (n = 2), and osteoarthritis (n = 2). Samples were processed across three batches: Batch 1 included 8 donors, whereas Batches 2 and 3 each included 16 donors distributed across two plates (8 donors per plate).

#### 4.9.2. Primary T cell Isolation and Banking

PBMCs were thawed in a 37°C waterbath, washed with complete media, and counted using an automated cell counter (Countess, Thermo). For the input PBMC population, 2e5 PBMCs per donor were immediately fixed. From the remaining PBMCs, CD3^+^ T cells were negatively selected using Human CD3 T Cell Enrichment kit (17951, StemCell) according to manufacturer’s instructions. Isolated T cells were counted using an automated cell counter (Countess, Thermo), and 2e5 T cells per donor were fixed as an input timepoint. Isolated T cells were resuspended in XVivo 15 media (04-418Q, Lonza) supplemented with 5% FBS (12306C-500ML, MilliporeSigma) in preparation for the *in vitro* cytokine perturbation assay.

#### 4.9.3. In Vitro Cytokine Perturbation Experiment

Isolated T cells were cultured with cytokines in XVivo 15 media supplemented with 5% FBS for 24 hours. Culture media was prepared with cytokines at the specified working concentrations (Table S7). After 24 hours, T cell samples were fixed using the GEM-X Flex Sample Preparation v2 Kit (10X Genomics, 1000781) according to manufacturer’s instructions. Samples were fixed in 96-well plates and stored short term at 4°C and long term at −80°C (10X Genomics Use Guide, CG000833 | Rev A) prior to subsequent GEM-X Flex v2 Gene Expression multiplex workflow.

#### 4.9.4. Single-Cell Sequencing of Basal T cell States and Perturbed States

Fixed samples, both the basal T cell states for each donor before any cytokine perturbation and also T cells with cytokine perturbation, were thawed and hybridized with GEM-X Flex Human Transcriptome Probe Kit v2, 96 samples (10X Genomics, 1000901) overnight at 42°C without further cell number standardization. After hybridization of the Barcode Oligo (10X Genomics, 1000894, 1000897, 1000898, 1000899), probed samples were pooled for post-hybridization washes followed by GEM generation, barcoding, recovery, and GEM-X Flex v2 library preparation according to the manufacturer protocol (10X Genomics Use Guide, CG000834 | Rev A). Specifically, 695,000 cells were loaded per GEM-X lane to target 500,000 cell recovery, and 4 individual index PCR reactions were prepared per lane. Indexed libraries were quantified by Qubit dsDNA Quantitation, High-Sensitivity kit (Thermo Fisher Q32851) for total yield and High-Sensitivity D1000 TapeStation (Agilent 5067-5585) to ensure expected library size. Pooled libraries were paired-end sequenced in house on Illumina NovaSeq X: Read 1: 54, I1: 10, I2: 10, Read 2: 50.

The demultiplexed FASTQ files generated from Illumina BCL Convert Software were then aligned to a human probe-set reference (v2.0.0_GRCh38-2024-A) using Cell Ranger v10.0.0 (10X Genomics).

#### 4.9.5. Preprocessing of DiseasePert-3M

Count matrices of DiseasePert-3M were independently preprocessed for each plate. Cells with fewer than 100 or more than 8,500 detected genes were removed. For cell type annotation, we performed standard Scanpy analysis as follows. Read counts were library-size normalized and log-transformed. The top 2,000 highly variable genes were selected for PCA, followed by neighbor graph construction, UMAP embedding, and Leiden clustering. Clusters were annotated to major T cell subtypes based on canonical marker gene expression. CD4 T cell subsets (naive, memory) were resolved at single-cell resolution using gene-signature scoring of naive markers (*CCR7*, *SELL*, *LEF1*, *TCF7*) versus memory markers (*S100A4*, *AQP3*, *ANXA1*, *CD44*). Low-quality clusters were removed.

To assess the concordance of transcriptional responses across donors, we quantified pairwise directed Jaccard similarity using cell-eval differential expression (DE) results. We first defined a shared set of 4,000 highly variable genes (HVGs) for DiseasePert-3M from a 1,000,000-cell subsample using the default Scanpy HVG pipeline with batch key provided. Within this gene set, for each donor pair under the same cytokine perturbation and within each T cell subtype (Naive CD4, Memory CD4, or Effector CD8), we identified differentially expressed genes (DEGs) at an FDR threshold of 0.05, requiring at least 50 shared genes per comparison. Directed Jaccard similarity was then computed as the fraction of genes in the union of the two DEG sets that were significant in both donors and changed in the same direction. Donor pairs with fewer than 10 union DEGs were excluded. For each cytokine, donor-pair similarity matrices were obtained by averaging directed Jaccard scores across subtypes. A summary matrix was then generated by computing the median and standard deviation of these cytokine-level similarities across all cytokines.

### 4.10. Stack donor-specific cytokine response prioritization

For donor-specific generation, we developed a synthetic-prompt-based procedure. We first constructed a synthetic prompt by adding the log-normalized difference between healthy-donor perturbed and control profiles to the log-normalized patient control query profile, after library-size normalization to 10,000 counts per cell. The resulting profile was then clipped at zero and projected back to count space using the library sizes of the healthy-donor perturbed cells. During construction of the synthetic prompt, cells from different populations were arbitrarily paired without one-to-one biological matching. Using Stack, we generated a predicted expression profile from this synthetic prompt together with the patient control query, and separately generated a synthetic control by using patient control cells as both prompt and query. To mitigate overfitting to the approximate synthetic prompt, here we used 1-step rather than the default 5-step generation. After library-size normalization to 10,000 counts per cell, we averaged the model-generated prediction with the healthy-donor perturbed profile, and analogously averaged the synthetic control with the healthy-donor control profile. This blending procedure enhanced power for detecting shared transcriptional signals across contexts of the same cell type, while injecting donor-specificity from model predictions.

We used cell-eval to derive differential expression (DE) results on the top 2,000 highly variable genes (HVGs) identified from the ground-truth perturbed and control data, consistent with prior benchmarks. Evaluations were performed independently for each T cell subpopulation: Naive CD4, Memory CD4, and Effector CD8. Prompt–query pairs were constructed by matching healthy donors and patients from the same batch (batches 1 and 3) or the same plate (batch 2). In addition to the prompt baseline and PerturbMean used in previous benchmarks, we introduced a new baseline in which the synthetic prompt was taken as the predicted expression (**synthetic prompt T cells**). For unmeasured patient perturbation conditions, direct evaluation against real measurements was not possible. We nevertheless used Stack to generate predictions for the subsequent gene module analysis, defining the 2,000 HVG list from the concatenation of two copies of the same patient control dataset. We also evaluated Stack in an additional setting using the previously described global set of 4,000 HVGs across all prompt–query pairs.

In addition to the standard cell-eval analysis, we quantified donor-specific prediction performance after excluding non-donor-specific genes identified from the prompt baseline. Specifically, for each prompt–query pair, genes that were significantly DE in both the query and prompt perturbation profiles with concordant effect directions were removed prior to metric computation. Using the filtered DE tables, we recomputed precision as the fraction of predicted significant DEGs recovered in the measured response, and directed precision as the fraction of predicted significant DEGs recovered with matched effect direction. To summarize donor-specific performance at the cytokine level, we further computed a global Spearman effect size within each T cell subset by calculating the fractions of significant DEGs in the measured and predicted responses after filtering, averaging these fractions within each query–cytokine combination with respect to prompt, and then taking the Spearman correlation across query–cytokine combinations. We focused on precision-based metrics because donor-specific DEG measurements in the ground-truth data are inherently noisy, making them more informative than metrics that depend strongly on complete gene set recovery.

To quantify donor-specific responses while accounting for both the total number of DEGs and variation in query cell number, we computed an adjusted donor-specific DEG ratio index for each prompt–query pair across all datasets and T cell subtypes. For each subtype, we collected DE results from the real measurement, the model prediction, and the prompt baseline, and defined DEGs as genes passing an FDR threshold of 0.05 with positive/negative log2 fold change. Unless otherwise specified, evaluations were performed using positively regulated DEGs. We then compared measured DE results versus prompt and model versus prompt separately. For each comparison, we first defined the raw donor-specific DEG ratio as the number of genes unique to either side divided by the number of shared DEGs, with a pseudocount of 5 added to both numerator and denominator for numerical stability. To account for differences in DEG burden, we divided this raw ratio by its expectation under random sampling from the full tested gene set. We then further adjusted the resulting ratio by regressing its log-transformed value on log1*𝑝* query cell number and dataset, separately for each comparison, and removing the fitted covariate effect while preserving the overall mean. The resulting quantity was used as the donor-specific effect size measure in downstream analyses. Conditions with insufficient cell counts were excluded.

For the eQTL enrichment analysis, we obtained T cell stimulation eQTLs from Soskic et al. (2022) and CD4 NC and CD8 ET eQTLs from the OneK1K resource (https://onek1k.org/) (Yazar et al., 2022). Each dataset was preprocessed by retaining the most significant locus per gene, yielding gene-level eQTL significance summaries. For each prompt–query pair and its associated set of 2,000 evaluated genes per subtype, we computed the eQTL gene ratio among real or predicted donor-specific up-regulated DEGs versus shared up-regulated DEGs between each prompt and query, with a pseudocount of 1 for numerical stability. Prompt– query pairs with undefined contingency-table statistics, resulting in undefined Fisher’s exact test *𝑝*-values, were excluded from the scatterplot visualization and significance analyses (Figs. 6G, S31).

To relate donor-specific effect sizes to baseline transcriptional programs, we performed a baseline signature association analysis for each T cell subset. Donor-level pseudobulk profiles were generated from control condition cells, normalized to counts per million, log1*𝑝* transformed, and filtered to exclude mitochondrial, ribosomal, and hemoglobin genes. Predefined gene-module scores were computed by averaging per-gene

*𝑧*-scores within each signature for each donor. We defined the patient signature score as a patient’s original *𝑧*-score minus the mean *𝑧*-score of its corresponding prompt healthy donors. These scores were then associated with the cytokine-specific adjusted donor-specific DEG ratio matrix across donors using Spearman correlation. We additionally fit cytokine-specific multivariable models using leave-one-out-cross-validated elastic net on standardized patient signature scores and batch covariates. We then refit ordinary least-squares models on the selected variables after standardizing both predictors and response, and used these models to report standardized coefficients.

## Data availability

Documentation on accessing scBaseCount can be found at https://github.com/ArcInstitute/arc-virtual-cell-atlas. The generated *Perturb Sapiens* data is deposited on Hugging Face: https://huggingface.co/datasets/arcinstitute/Perturb-Sapiens. The CELLxGENE 45M data used for post-training is also available on Hugging Face: https://huggingface.co/arcinstitute/Stack-CellxGene45M. The Parse 10M PBMC data is available at https://www.parsebiosciences.com/datasets/10-million-human-pbmcs-in-a-single-experiment/#download. We downloaded the primary human CD4 T cell Perturb-seq data from the CZI Virtual Cell and AWS portals as instructed at https://virtualcellmodels.cziscience.com/dataset/genome-scale-tcell-perturb-seq. The DiseasePert-3M data will be available upon publication. All evaluation data used for this project are publicly available; see the Methods section for download details of other datasets.

## Code and model availability

Code for the Stack model is available at https://github.com/ArcInstitute/stack. Model parameters are available on Hugging Face: https://huggingface.co/arcinstitute/Stack-Large, https://huggingface.co/arcinstitute/Stack-Large-Aligned.

## Acknowledgments

We thank Beatrice Bevilacqua, Arshia Nayebnazar, Basak Eraslan, Rishi Verma, Noam Teyssier, Brian Plosky, Silvana Konermann, and Patrick Hsu for helpful discussions. This study was supported by Arc Institute. M.D. and Y.K. acknowledge the support from NIH [U54AG076043, U54AG079759, R01DA063148, UM1DA051410]. We also acknowledge the efforts of our colleagues to generate and release large-scale datasets necessary to train and evaluate our models.

## Author contributions

M.D. conceived the Stack project with input from Y.K. and Y.H.R. Y.H.R., T.L.R., and D.P.B. supervised the project, with Y.H.R. coordinating effort across the team. M.D. developed the data loaders, architecture, pre-training, and post-training of the Stack model, and implemented a unified Stack codebase. C.C., R.S., A.A., and M.D. curated pre-training data for the model. M.D. prepared post-training data and curated evaluation data. M.D. and Y.H.R. designed evaluation metrics and layout of results. M.D. and A.A. designed and ran baseline embedding and perturbation models. Y.H.R., T.L.R., M.D., N.L. and P.Y.T. designed the validation experiments with patient samples. M.M.C., C.K., L.W., Y.C.C., P.Y.T., N.L., T.L.R., and A.D. collected and processed DiseasePert-3M. M.D. performed analysis in the manuscript and visualized results. D.G. analyzed Stack gene modules, improved the codebase, and contributed to the inference scheme for the post-trained Stack model. M.D., C.R.T., and Y.H.R. created visuals for main text figures. M.D. wrote the first draft of the manuscript. All authors wrote the final draft of the manuscript.

## Supplementary Figures and Tables

**Figure S1.**
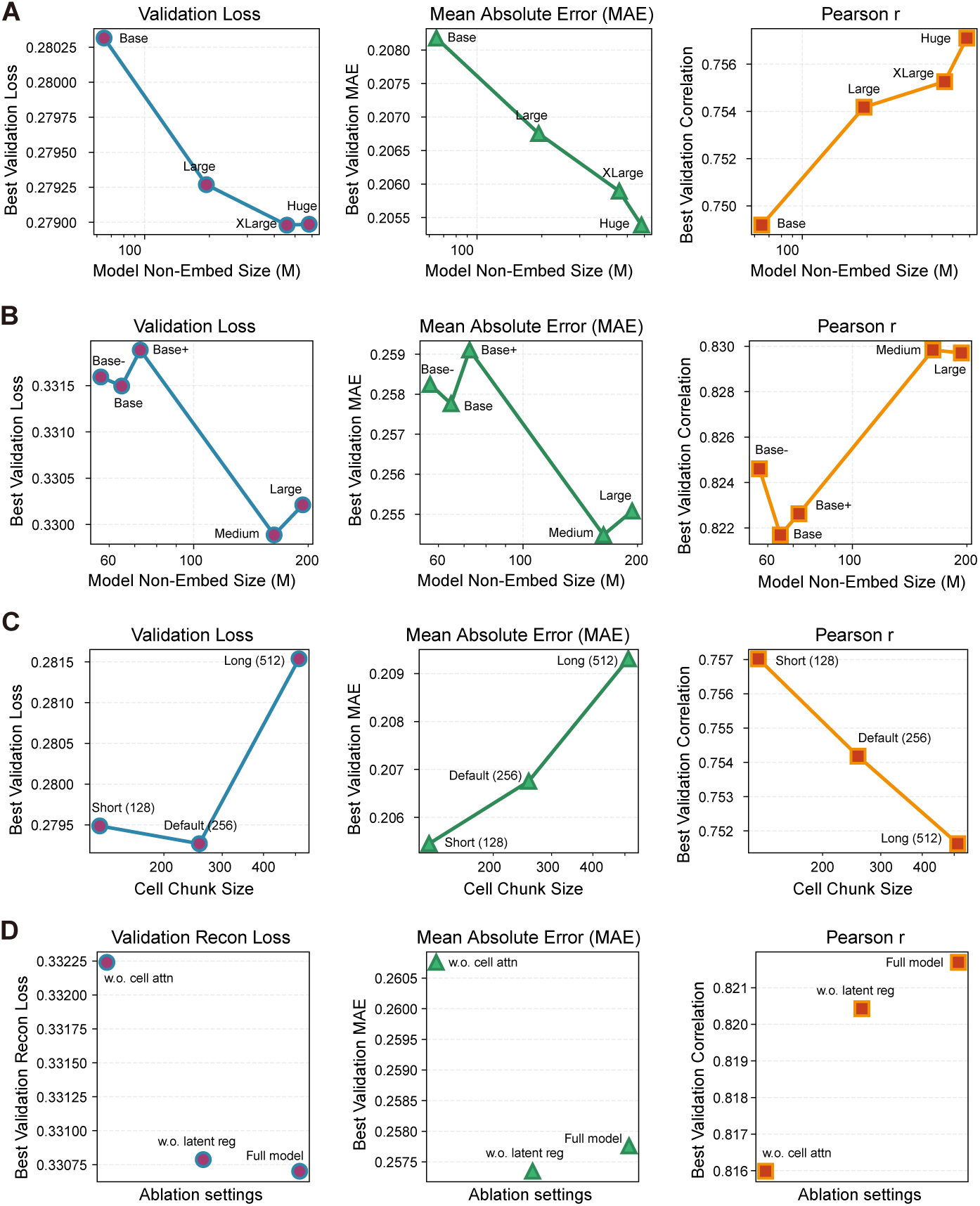
Scaling and ablation analysis of the Stack model. **A.** Validation performance across model sizes for Stack models trained on the full human scBaseCount (Youngblut et al., 2025). **B.** Validation performance across model sizes for Stack models trained on the scBaseCount subset. **C.** Validation performance across cell set sizes for the Stack (Large) model trained on the full scBaseCount dataset. **D.** Validation performance across ablation settings for Stack (Base) models trained on scBaseCount subset (w.o. latent reg: removing only latent regularization; w.o. cell attn: removing both latent regularization and inter-cellular attention). All models in **A**, **B** and **D** use a cell set size of 256. For validation loss and mean absolute error (MAE), smaller is better; for Pearson r, higher is better.

**Figure S2.**
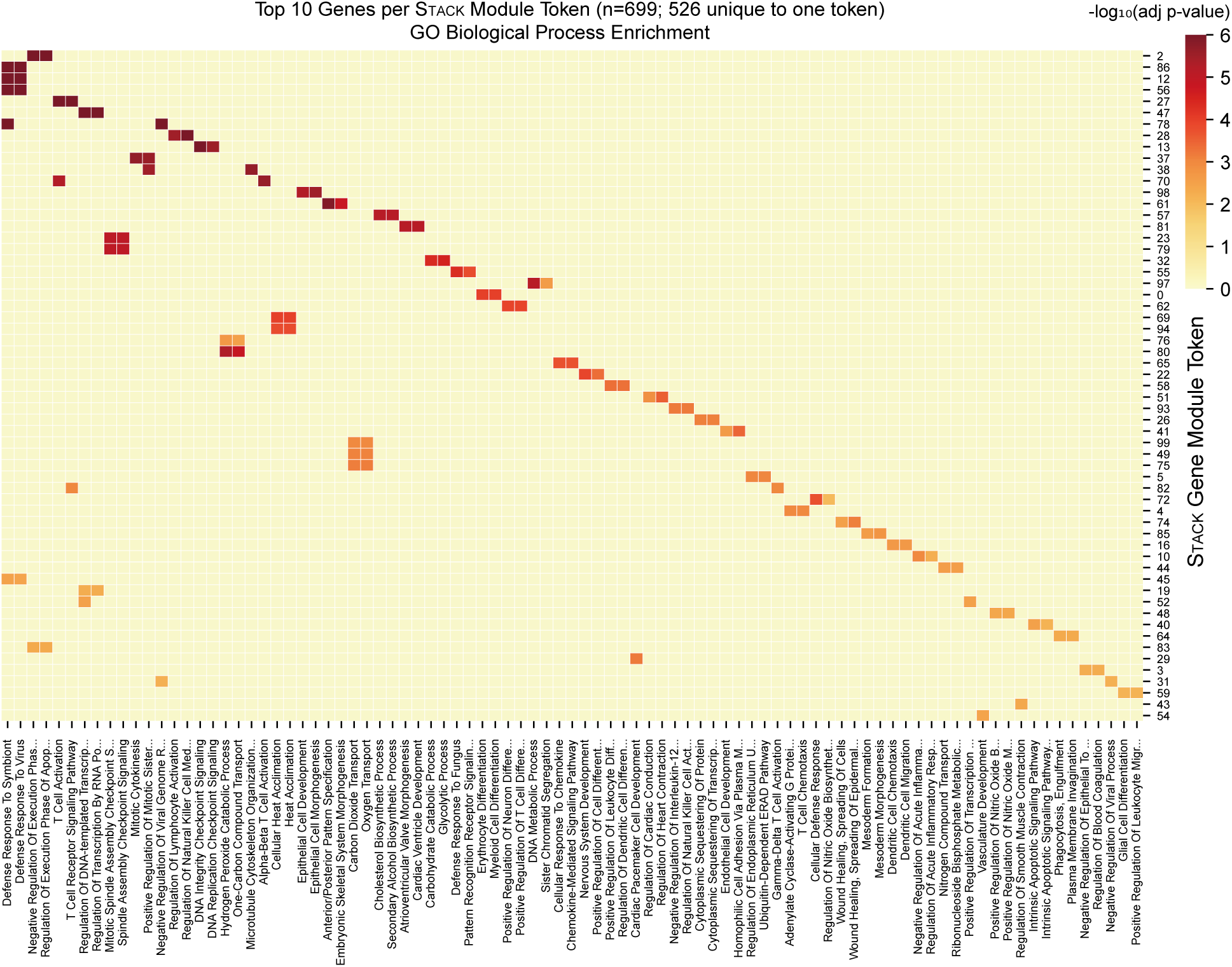
Adjusted *𝑝*-value heatmap of Gene Ontology (GO) biological process enrichment analysis for the top 10 most important genes within each Stack (Large) token after pre-training on full human scBaseCount. Gene importance scores were computed by first reshaping the tokenization weight matrix **W** to ℝ*^𝑛^*hidden ×*𝑑*token ×*𝑛*genes, then calculating the mean absolute weight across the token dimension for each hidden module. The top 10 genes per module were selected based on these importance scores. The top 2 enriched pathways per module are shown. Rows (modules) and columns (pathways) were ordered by hierarchical clustering using average linkage with Euclidean distance.

**Figure S3.**
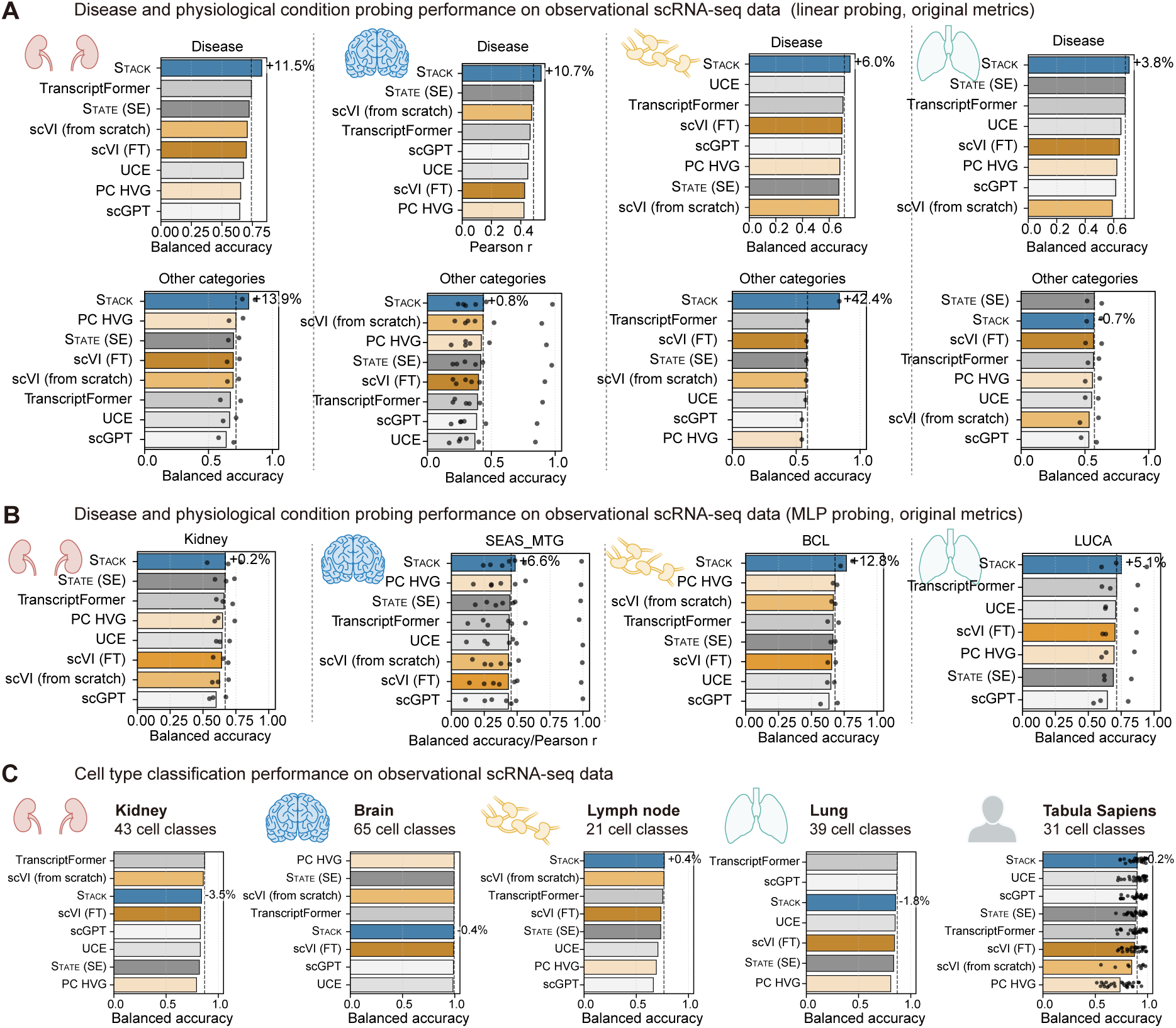
Additional probing evaluation results. **A.** Disease and physiological condition probing performance on observational scRNA-seq data (De Boer et al., 2021; Li et al., 2025; Salcher et al., 2022; Gabitto et al., 2024). One linear classifier is trained per cell type. For the “other categories” group, each point represents one metadata prediction task (*𝑛* = 2, 6, 1, 2). See Fig. S4 for full results. All Stack results presented here are based on one model with the (Large) setting pre-trained on full human scBaseCount. All panels show (average) values of original metrics (balanced accuracy/Pearson r) without normalization. **B.** Disease and physiological condition MLP probing performance on observational scRNA-seq data. One MLP classifier is trained simultaneously on the top 5 most abundant cell types. See Methods for dataset statistics. Each point represents one metadata prediction task (*𝑛* = 3, 7, 2, 3). **C.** Cell type classification performance on observational scRNA-seq data. In Tabula Sapiens evaluation, each point represents a tissue (n=26).

**Figure S4.**
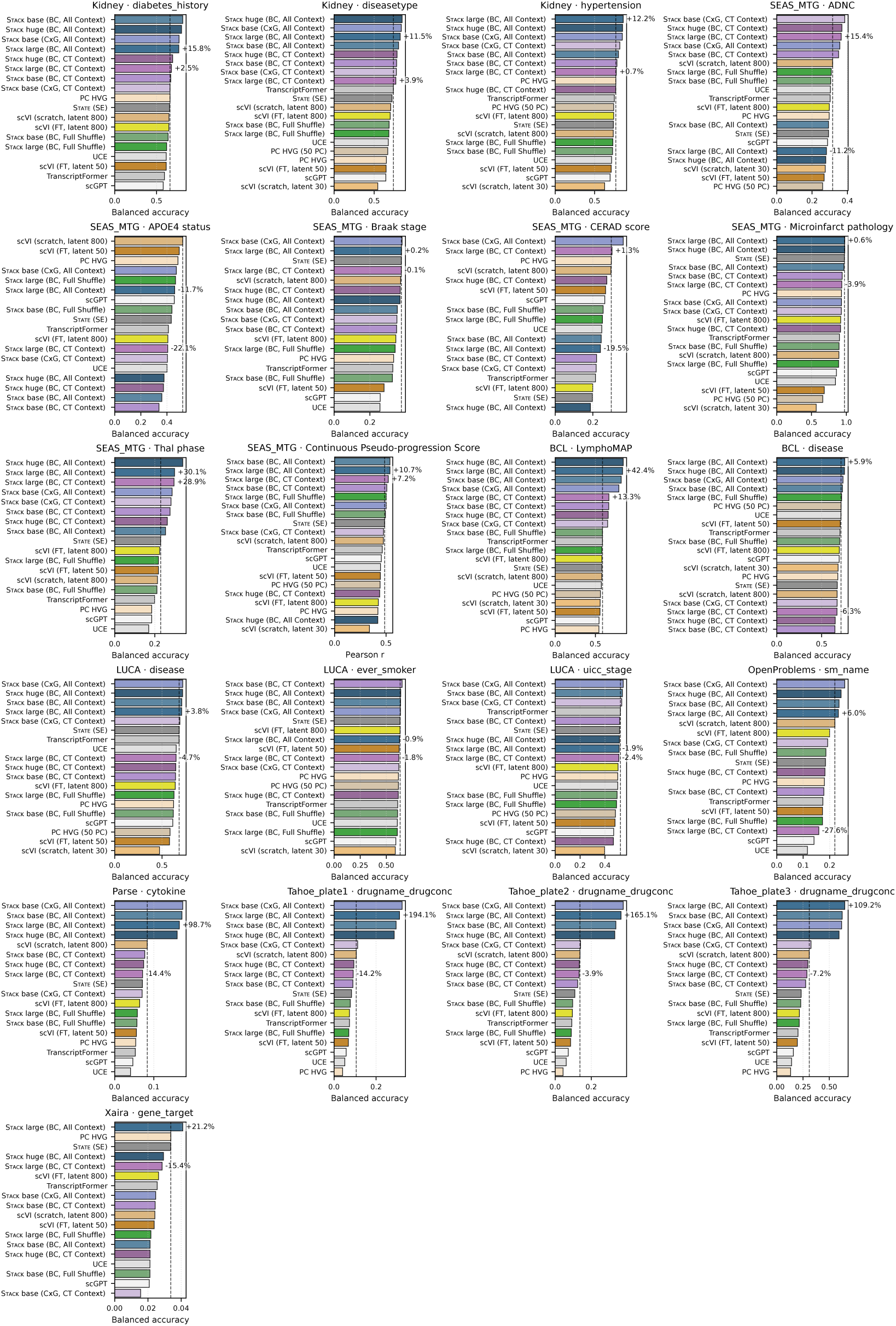
Per-cell-type linear probing results with additional Stack settings. **BC:** the model is trained on full human scBaseCount. CxG: the model is trained on CELLxGENE. All Context: Stack uses the default dataset grouping by sample, utilizing all cell types from each sample as context. CT Context: Stack utilizes cells grouped by cell type per sample as context, generates a set of embeddings for each group, then concatenates them to form the total embedding. Full shuffle: The cell order in the evaluation data is randomly shuffled, effectively removing context information. The number of experiments per dataset (equal to evaluated cell types): *𝑛* = 12, 10, 20, 20, 6, 17, 20, 20, 20, 2.

**Figure S5.**
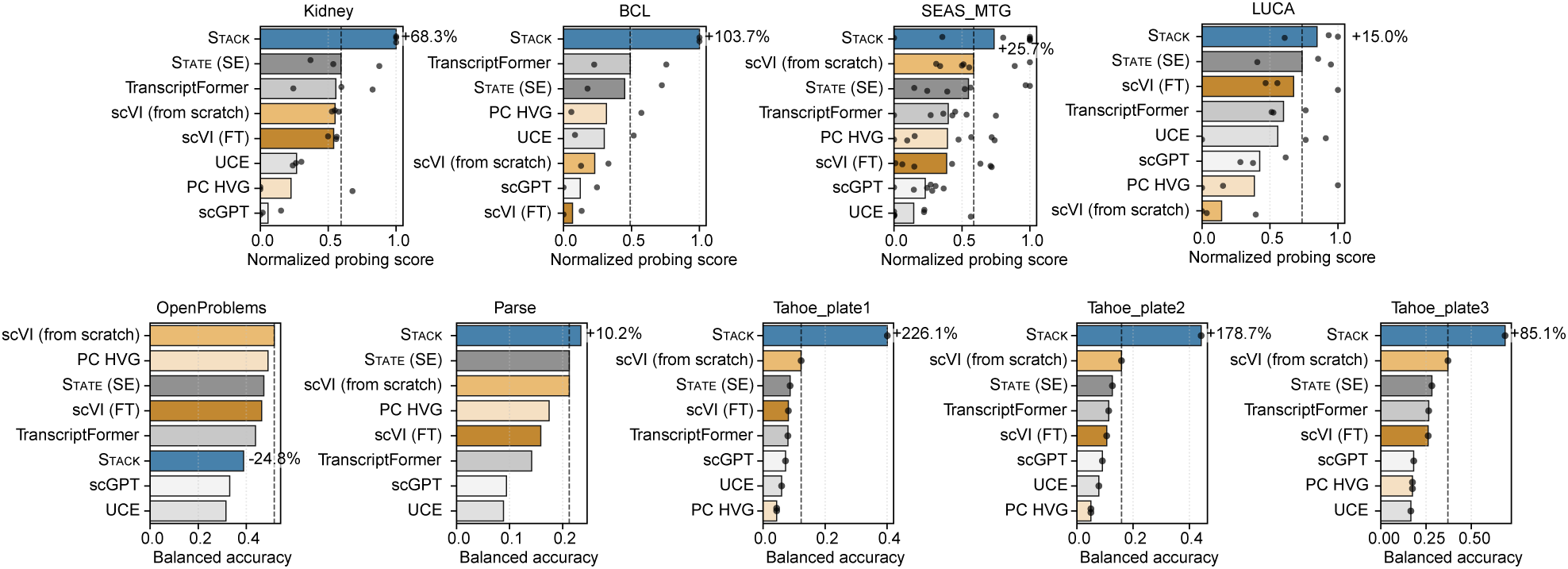
Linear probing results for each dataset, using the overall best-performing cell type per task. Each point represents one metadata prediction task (*𝑛* = 3, 2, 7, 3, 1, 1, 1, 1, 1). The full results for Xaira are included in Fig. 2.

**Figure S6.**
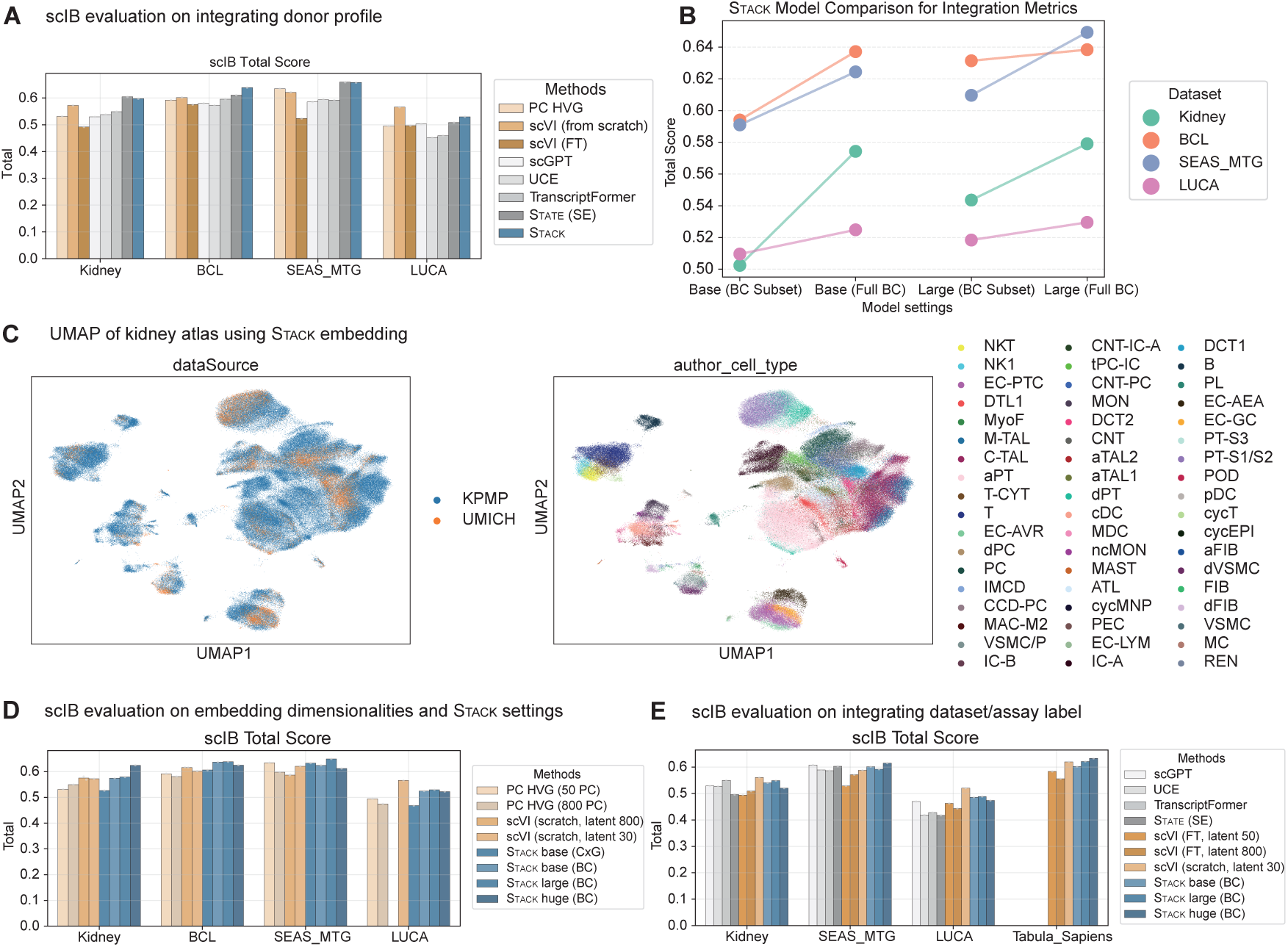
Additional evaluation results of Stack on batch integration. **A.** scIB evaluation of different methods on integrating donor profiles and preserving cell types (Luecken et al., 2022). Presented Stack results are based on one model with the (Large) setting pre-trained on full human scBaseCount. **B.** scIB batch integration total scores of Stack with different sizes, training data, and evaluation datasets. **C.** UMAP visualization of Stack embedding of the Kidney atlas, colored by data collection and fine-grained cell type. **D.** Comparison of integration performance across different numbers of principal components and scVI latent dimensions. Based on these results, the results of 50 principal components and 30 scVI latent dimensions were selected for main batch integration benchmarks. **E.** Comparison of dataset label integration performance. As BCL involves only a single dataset label, it is not applicable to dataset label integration evaluation. Alternative foundation models (scGPT, UCE, State (SE), and TranscriptFormer) were excluded from the Tabula Sapiens evaluation due to their limited performance improvements in the per-tissue benchmark. In **B**, **D**, and **E**, we apply Stack to the full dataset rather than to one sample at a time. This results in a very minor decrease in Stack’s performance. The dataset integration performance in **E** closely matches the donor integration performance shown in **A**.

**Figure S7.**
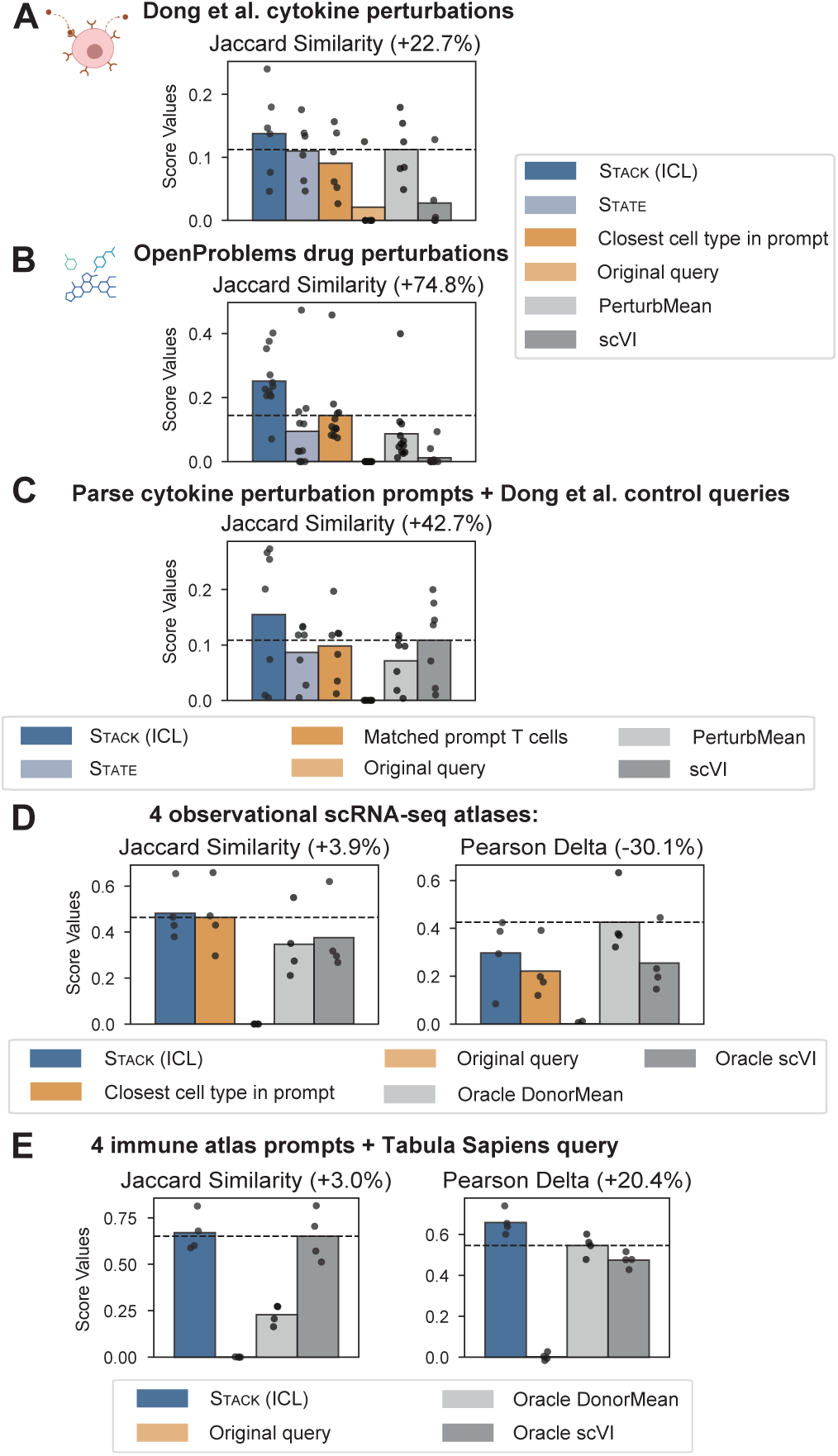
Additional metrics on cell prompting tasks. **A.** Evaluation of perturbation effect prediction across cell types on the Dong et al. (2023) cytokine perturbation dataset (6 cytokines). **B.** Evaluation of perturbation effect prediction across cell types on the OpenProblems drug perturbation dataset (Luecken et al., 2025) (12 drugs). **C.** Evaluation of T cell response prediction across samples (7 cytokine stimulation conditions). **D.** Evaluation of donor-specific gene expression generation across five atlases. **E.** Evaluation of condition-specific expression generation across four PBMC atlases. See Fig. 3 caption for additional details. For all scores, higher values indicate better performance.

**Figure S8.**
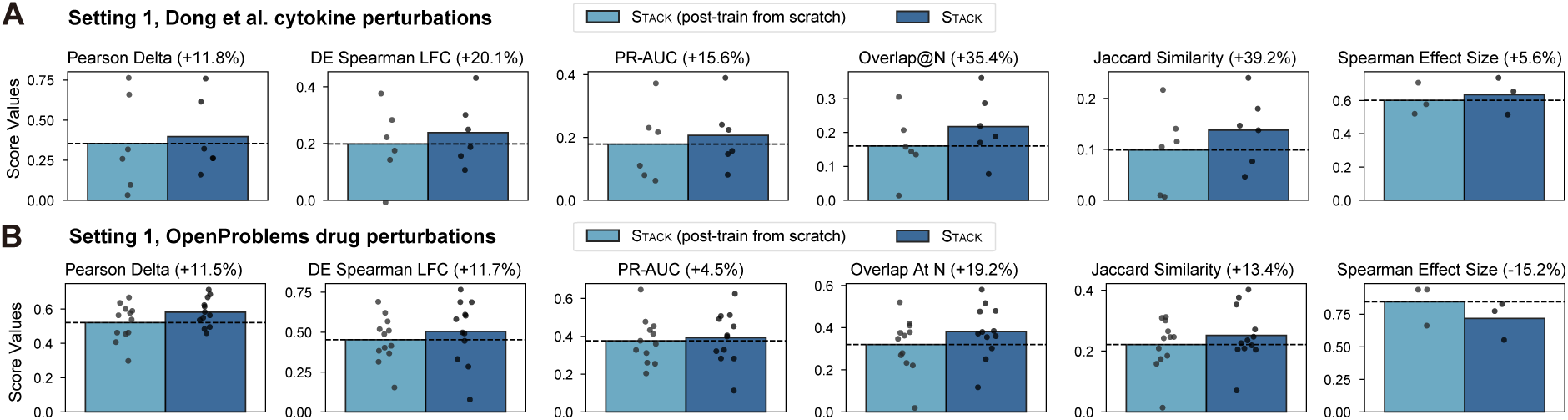
Effect of pre-training on Stack perturbation response prediction in novel cell types. **A.** Results on the Dong et al. (2023) dataset. **B.** Results on the OpenProblems drug perturbation dataset (Luecken et al., 2025). See the Fig. 3 caption for the number of experiments and the definition of scores. All percentages shown represent the average performance improvement of Stack over Stack post-trained from scratch.

**Figure S9.**
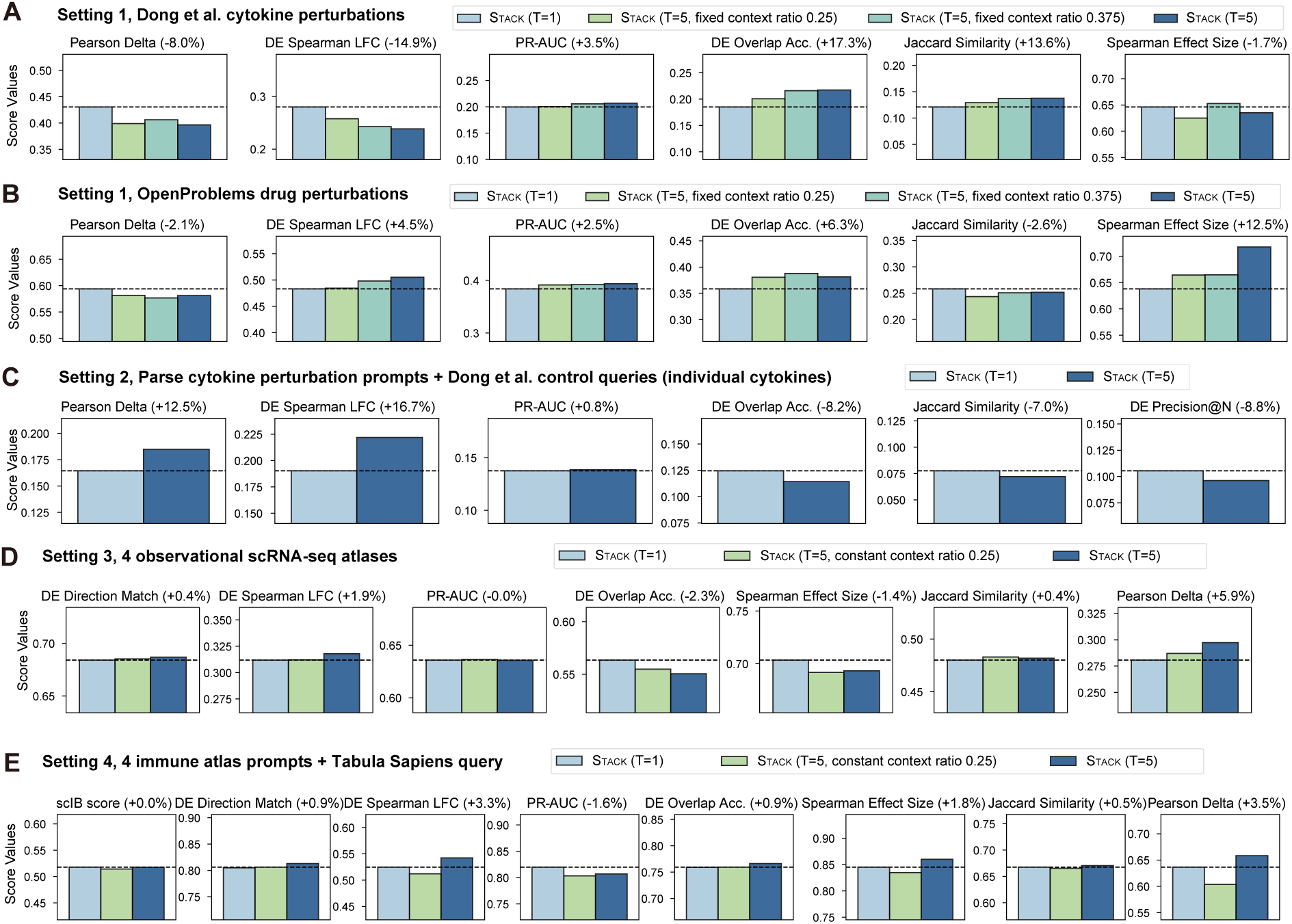
Evaluations of Stack generative procedure on cell prompting tasks. **A.** Comparison of Stack (*𝑇* = 5) and Stack (*𝑇* = 1) for perturbation effect prediction across cell types on the Dong et al. (2023) dataset. **B.** Comparison of Stack (*𝑇* = 5) and Stack (*𝑇* = 1) for perturbation effect prediction across cell types on the OpenProblems drug perturbation dataset (Luecken et al., 2025). **C.** Evaluation of T cell perturbation response prediction across Parse PBMC and Dong et al. (2023) with individual cytokine stimulation conditions. **D.** Evaluation of donor-specific gene expression generation across four atlases. **E.** Evaluation of condition-specific expression generation across four PBMC atlases as prompts and Tabula Sapiens as queries. All percentages shown represent the average performance improvement of Stack generative settings over predictive settings. See the Fig. 3 caption for the number of experiments and the definition of scores. For all scores, higher values indicate better performance.

**Figure S10.**
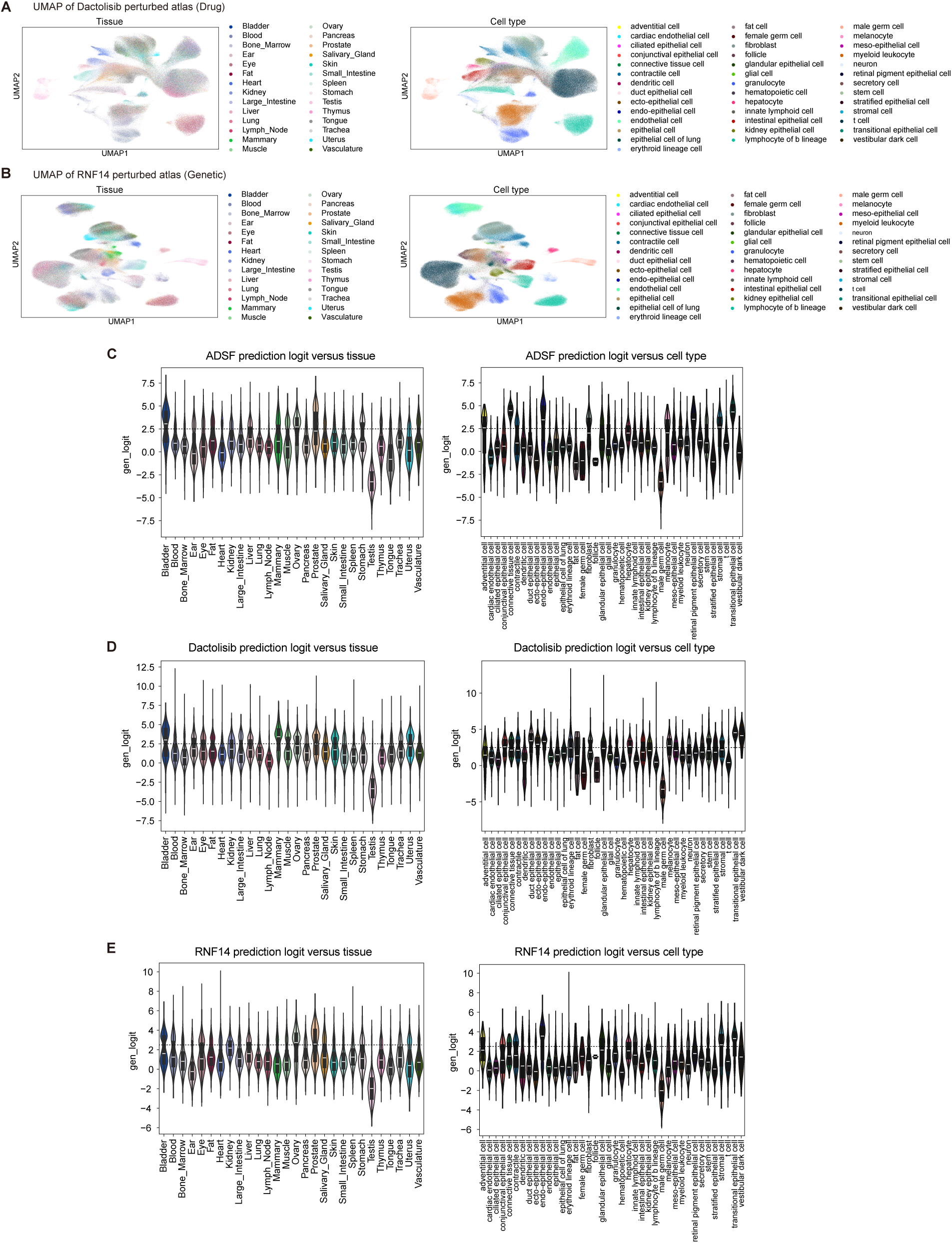
Additional analysis on *Perturb Sapiens*. **A–B.** UMAP visualizations of Stack embeddings, colored by tissue and cell type, for dactolisib *Perturb Sapiens* donor 2 (**A**) and *RNF14 Perturb Sapiens* donor 4 (**B**). **C–E.** Violin plots of classifier-predicted logit values, grouped by tissue and cell type, for ADSF *Perturb Sapiens* donor 1 (**C**), dactolisib *Perturb Sapiens* donor 2 (**D**), and *RNF14 Perturb Sapiens* donor 4 (**E**). Lower logit indicates higher generation confidence.

**Figure S11.**
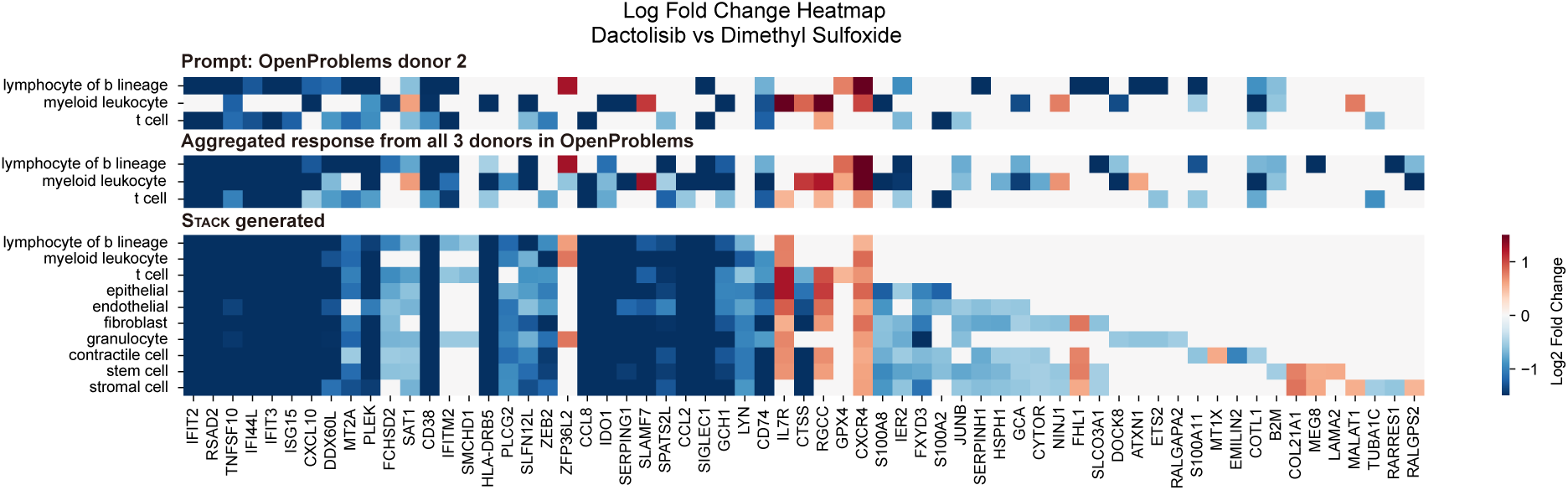
Log2-fold-change heatmap of dactolisib *Perturb Sapiens* versus control. In Figs. S11-S14, only significantly changed genes are shown in color; the rest are colored gray.

**Figure S12.**
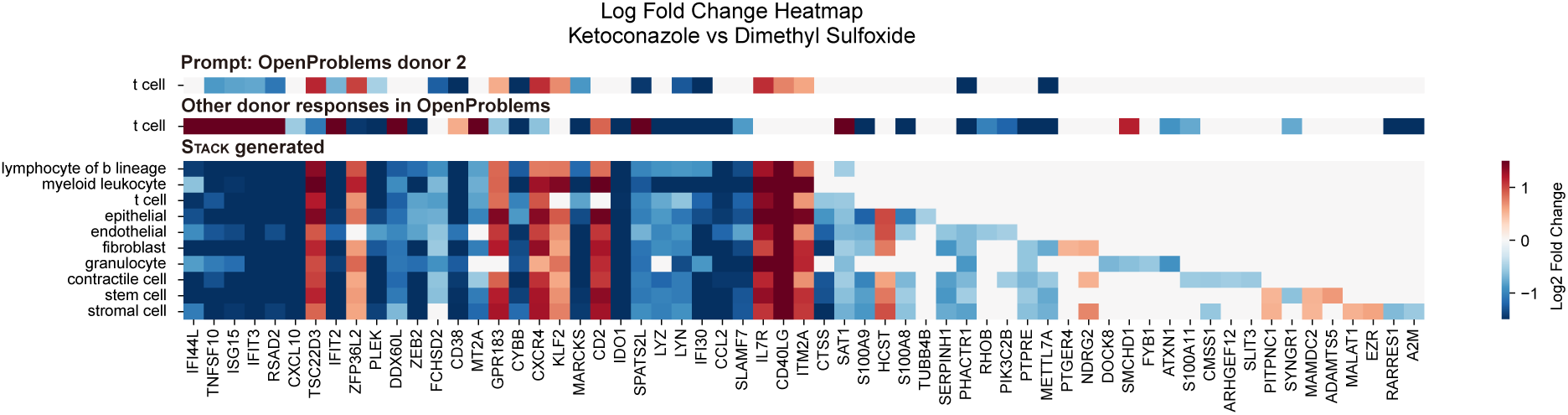
Log2-fold-change heatmap of ketoconazole *Perturb Sapiens* versus control.

**Figure S13.**
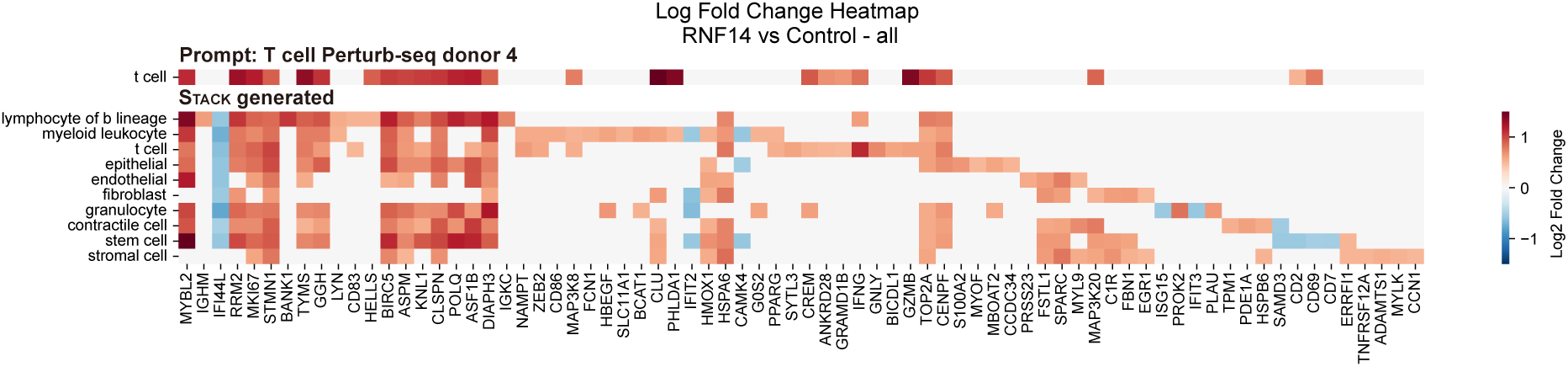
Log2-fold-change heatmap of *RNF14 Perturb Sapiens* versus control.

**Figure S14.**
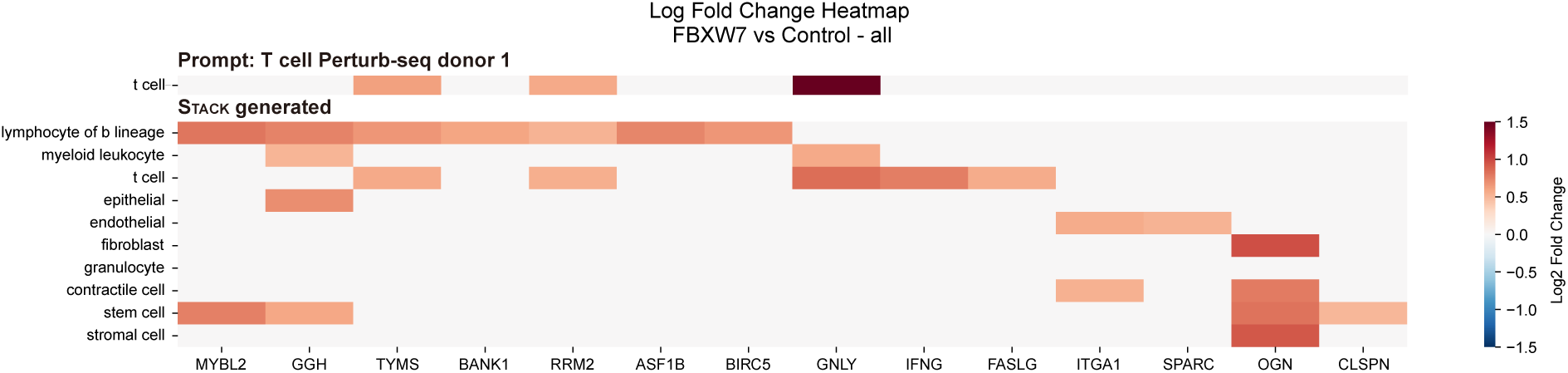
Log2-fold-change heatmap of *FBXW7 Perturb Sapiens* versus control.

**Figure S15.**
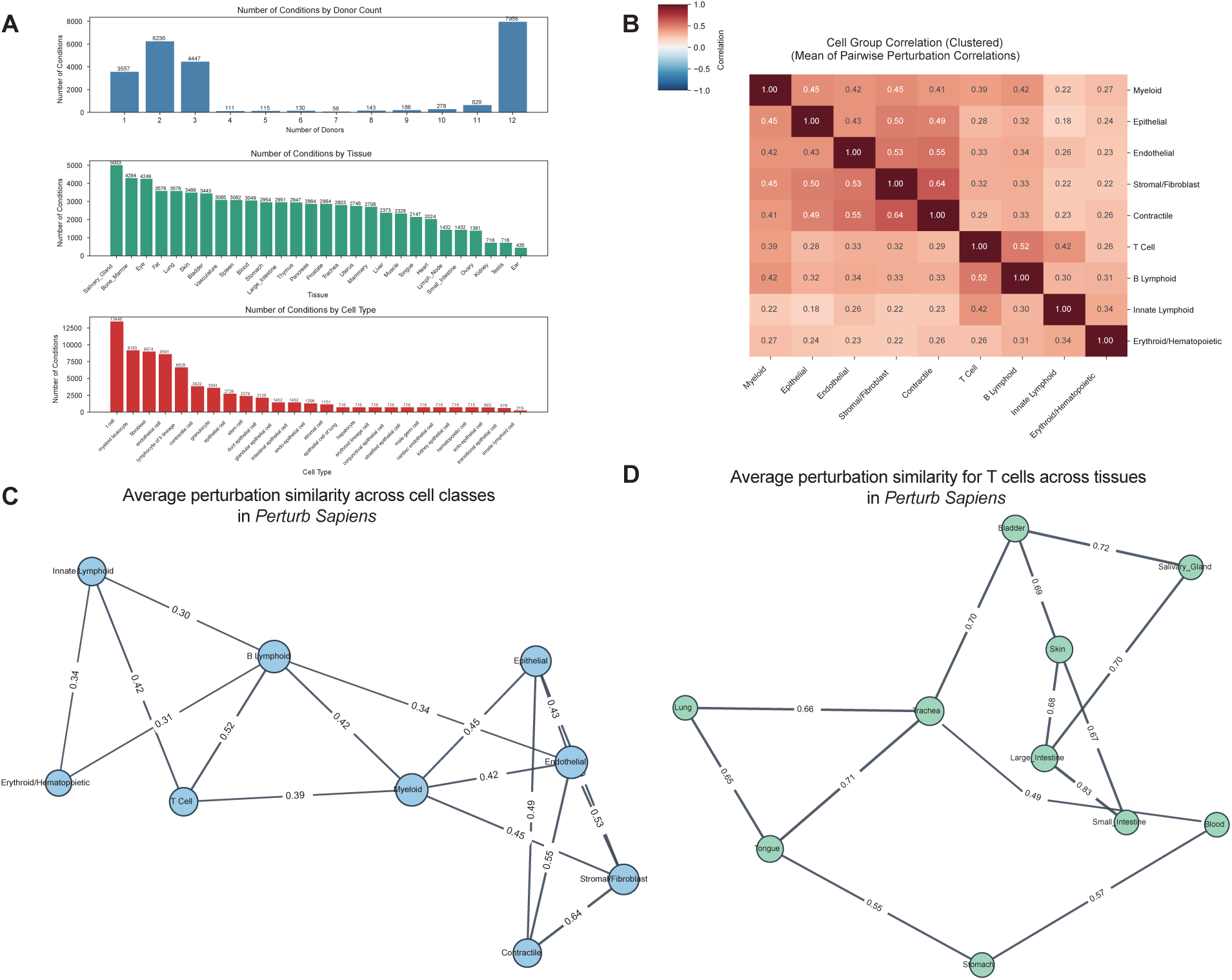
Global characterization of *Perturb Sapiens*. **A.** Overview of the number of conditions by donor count, tissue, and cell type represented in *Perturb Sapiens*. **B.** Heatmap of mean perturbation similarity across cell classes in *Perturb Sapiens*. **C.** Network plot of mean perturbation similarity across cell classes in *Perturb Sapiens*. **D.** Network plot of mean perturbation similarity for T cells across representative tissues in *Perturb Sapiens*. In **B–D**, values represent mean Spearman correlation coefficients, averaged across perturbations using Fisher’s *𝑧*-transformation. For each node, the top two or three edges by correlation strength are displayed.

**Figure S16.**
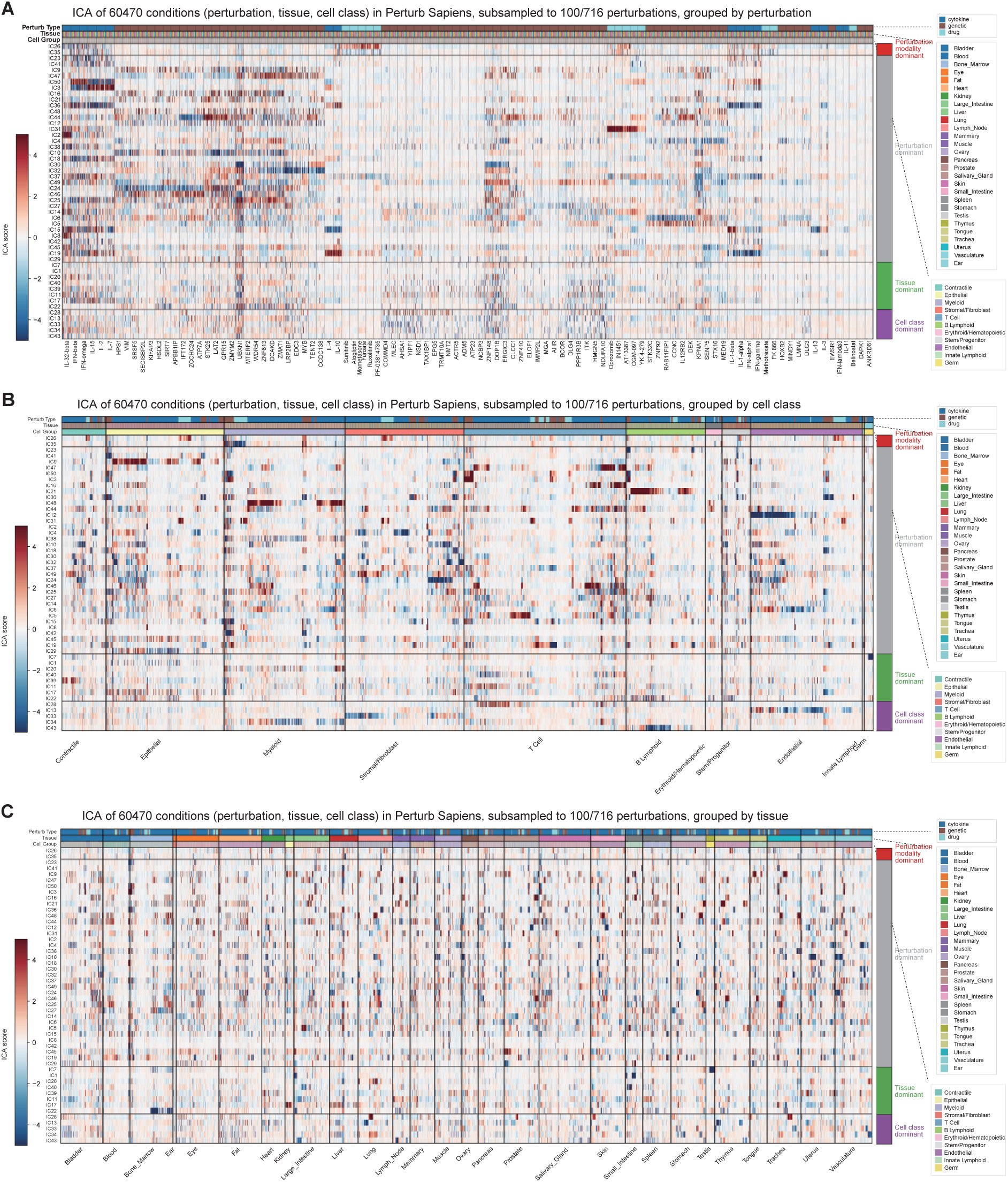
Independent component decomposition of *Perturb Sapiens*. Heatmaps of independent components (ICs) derived from *Perturb Sapiens* perturbation effect profiles (*𝑘* = 50 components). **A.** Ordered by perturbation condition. **B.** Ordered by cell class. **C.** Ordered by tissue.

**Figure S17.**
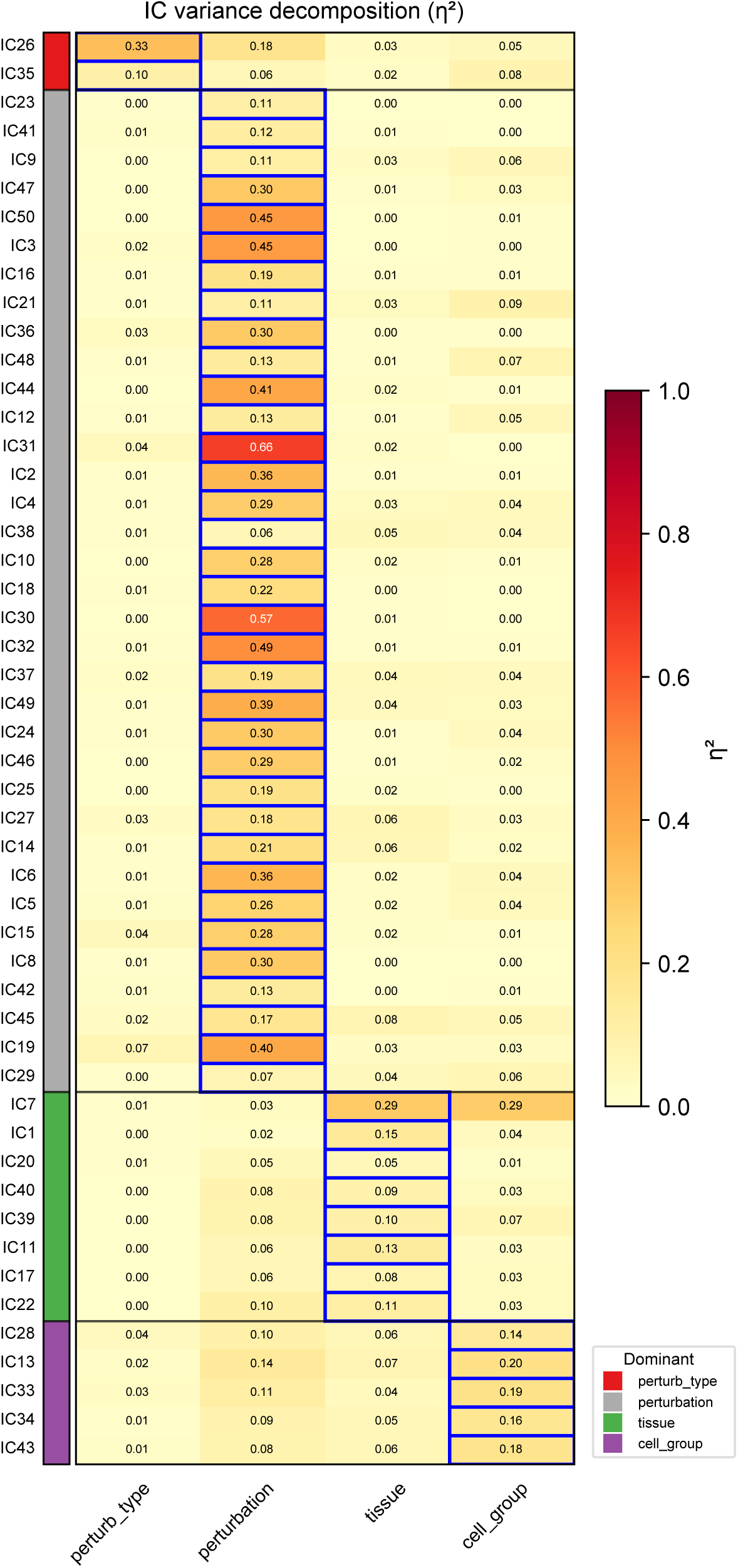
Variance decomposition of independent components (ICs) across perturbation modality (drug, cytokine, or genetic), perturbation identity (conditioned on modality), tissue, and cell class. ICs are ordered and colored by their dominant source of variation.

**Figure S18.**
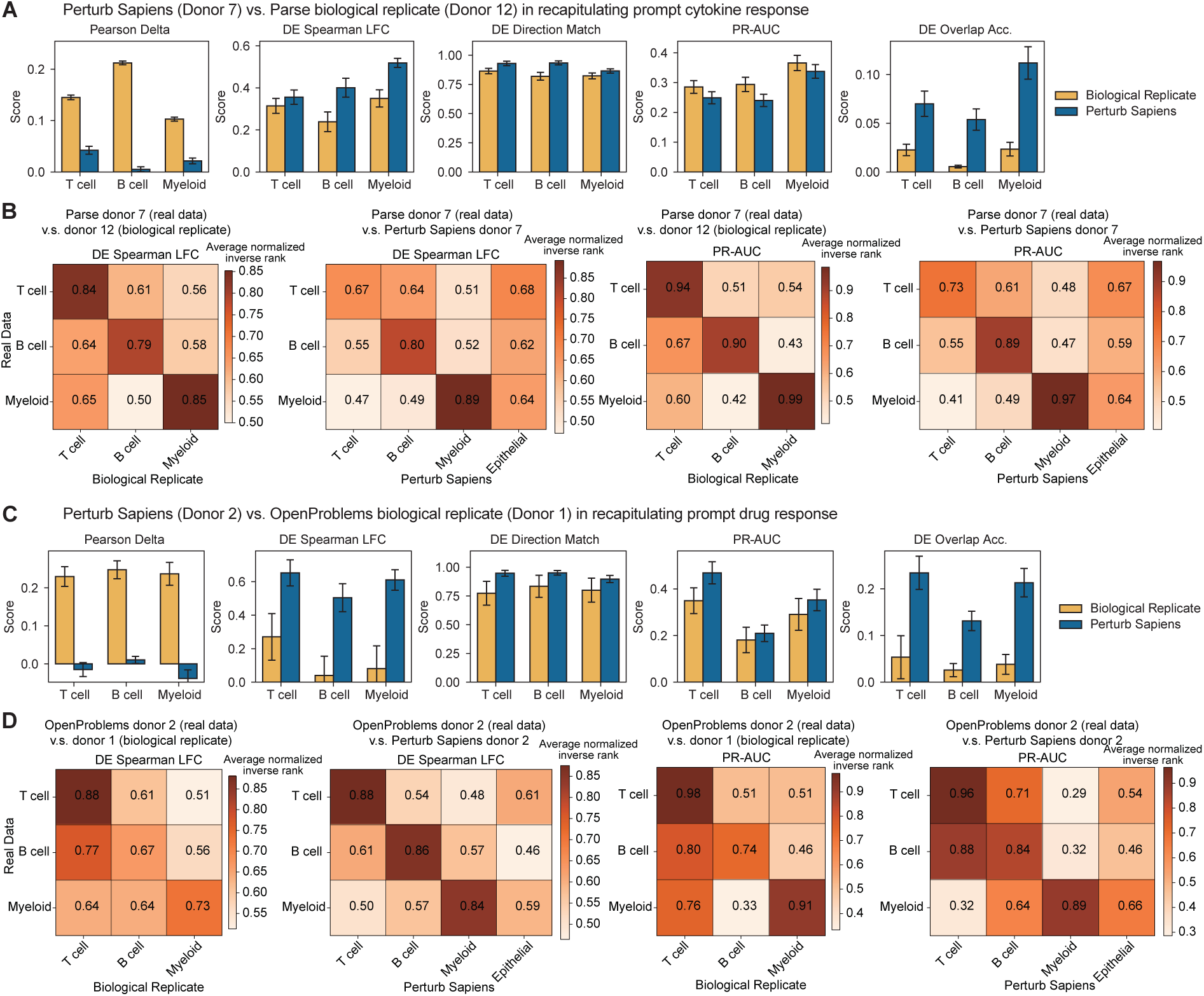
Comparison of *Perturb Sapiens* with biological replicates using prompt data as reference. **A.** Cell-eval metric comparison between *Perturb Sapiens* (cytokine donor 7) and a biological replicate (cytokine donor 12). These two donors were selected based on the similarity of their responses (Klein et al., 2025). Bars indicate mean scores; error bars indicate standard errors. **B.** Heatmap of normalized inverse rank scores for DE Spearman LFC and PR-AUC across cell-type pairs, comparing the biological replicate (cytokine donor 12; left) and *Perturb Sapiens* prediction (cytokine donor 7; right). **C.** Cell-eval metric comparison between *Perturb Sapiens* (drug donor 2) and a biological replicate (drug donor 1). **D.** Heatmap of normalized inverse rank scores for DE Spearman LFC and PR-AUC across cell-type pairs, comparing the biological replicate (drug donor 1; left) and *Perturb Sapiens* prediction (drug donor 2; right).

**Figure S19.**
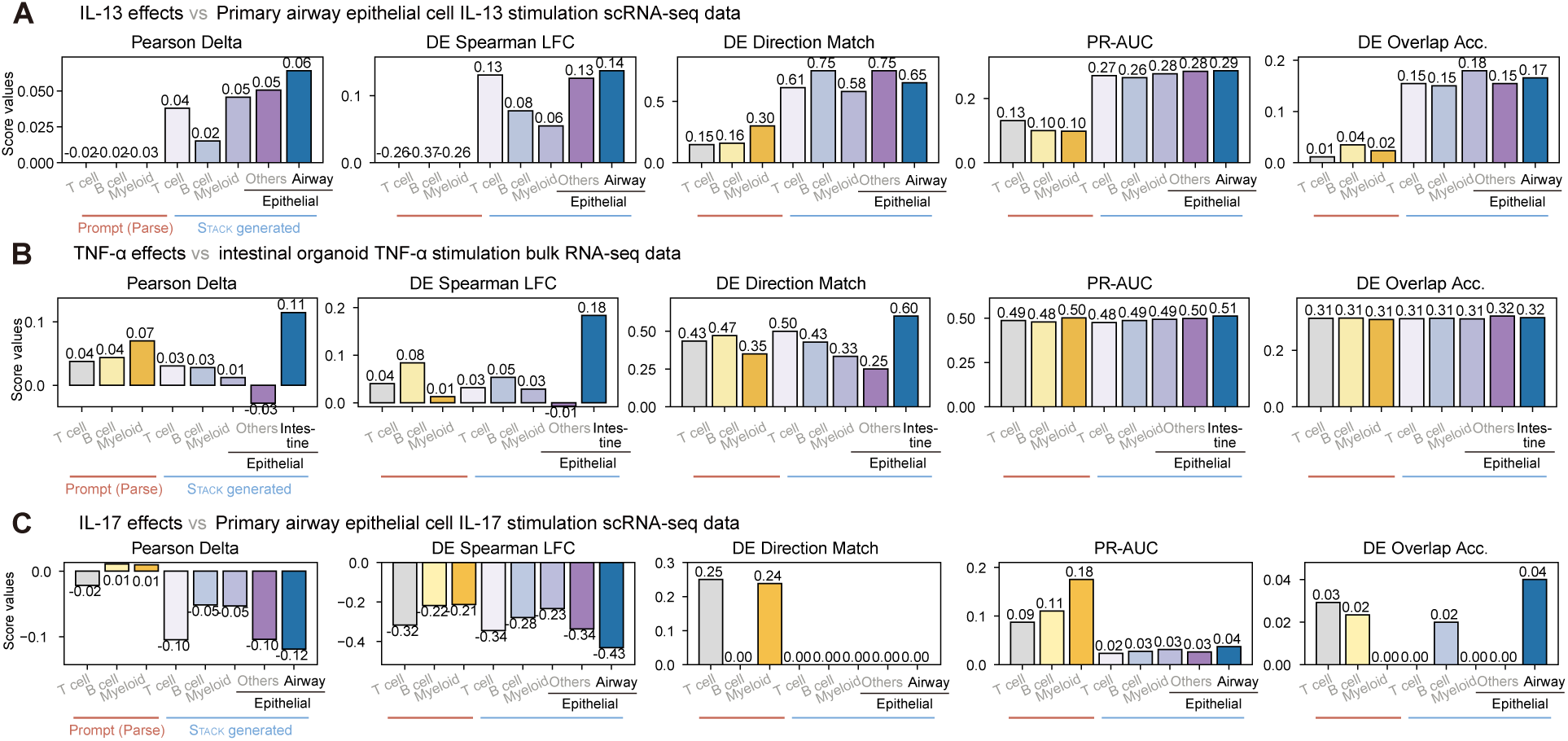
Evaluation of *Perturb Sapiens* epithelial cytokine effects using external stimulation datasets. **A.** Interleukin-13 (IL-13) effects, evaluated using single-cell IL-13 stimulation data from primary airway epithelial cells (Koh et al., 2023). **B.** Tumor necrosis factor alpha (TNF-*𝛼*) effects, evaluated using intestinal TNF-*𝛼* stimulation data (Lee et al., 2022). **C.** Interleukin-17 (IL-17) effects, evaluated using single-cell IL-17 stimulation data from primary airway epithelial cells (Koh et al., 2023). For all panels, log2-fold-changes were aggregated as the median across all 12 donors, with per-donor *𝑝*-values combined using a Beta-distribution test based on the median *𝑝*-value (Methods), followed by Benjamini–Hochberg FDR correction.

**Figure S20.**
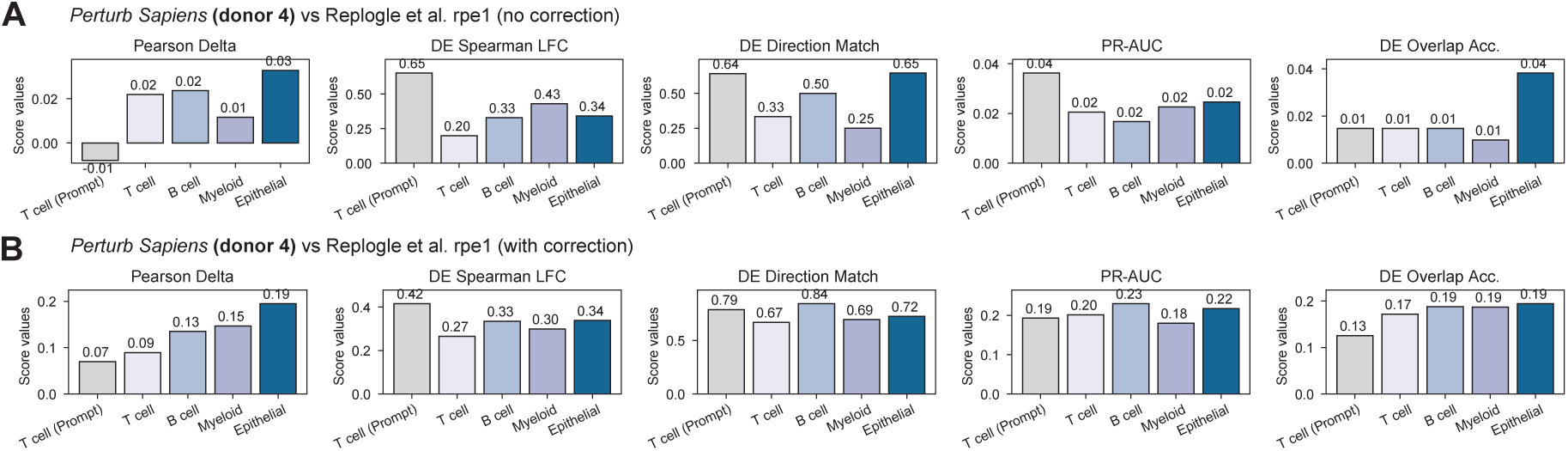
Evaluation of *Perturb Sapiens* (genetic perturbation donor 4) on the RPE1 dataset (Replogle et al., 2022), without (**A**) and with (**B**) control correction. The epithelial category includes all epithelial cells in *Perturb Sapiens*. Bars show mean scores over the 14 perturbations shared between *Perturb Sapiens* (genetic perturbation donor 4) and the RPE1 dataset.

**Figure S21.**
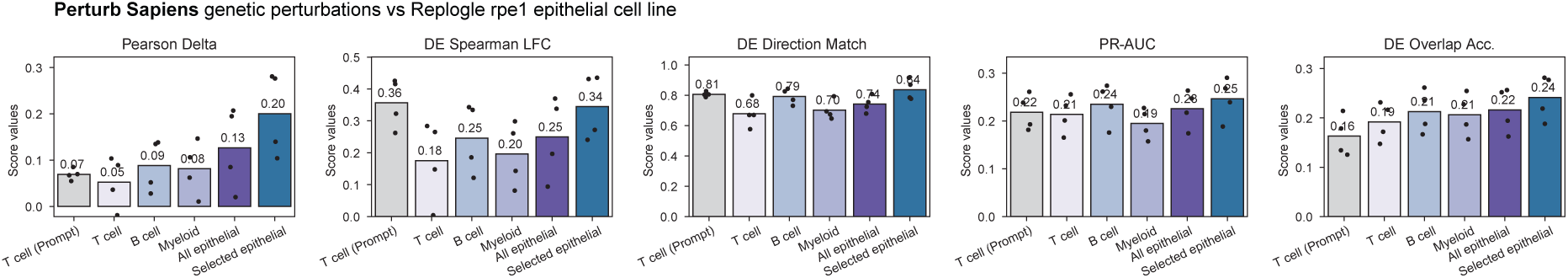
Evaluation of *Perturb Sapiens* genetic perturbations against RPE1 epithelial cell line perturbation measurements (Replogle et al., 2022). Selected epithelial cells refer to *Perturb Sapiens* epithelial populations from the lung and trachea. Each point represents the average result for one prompt donor (*𝑛* = 4).

**Figure S22.**
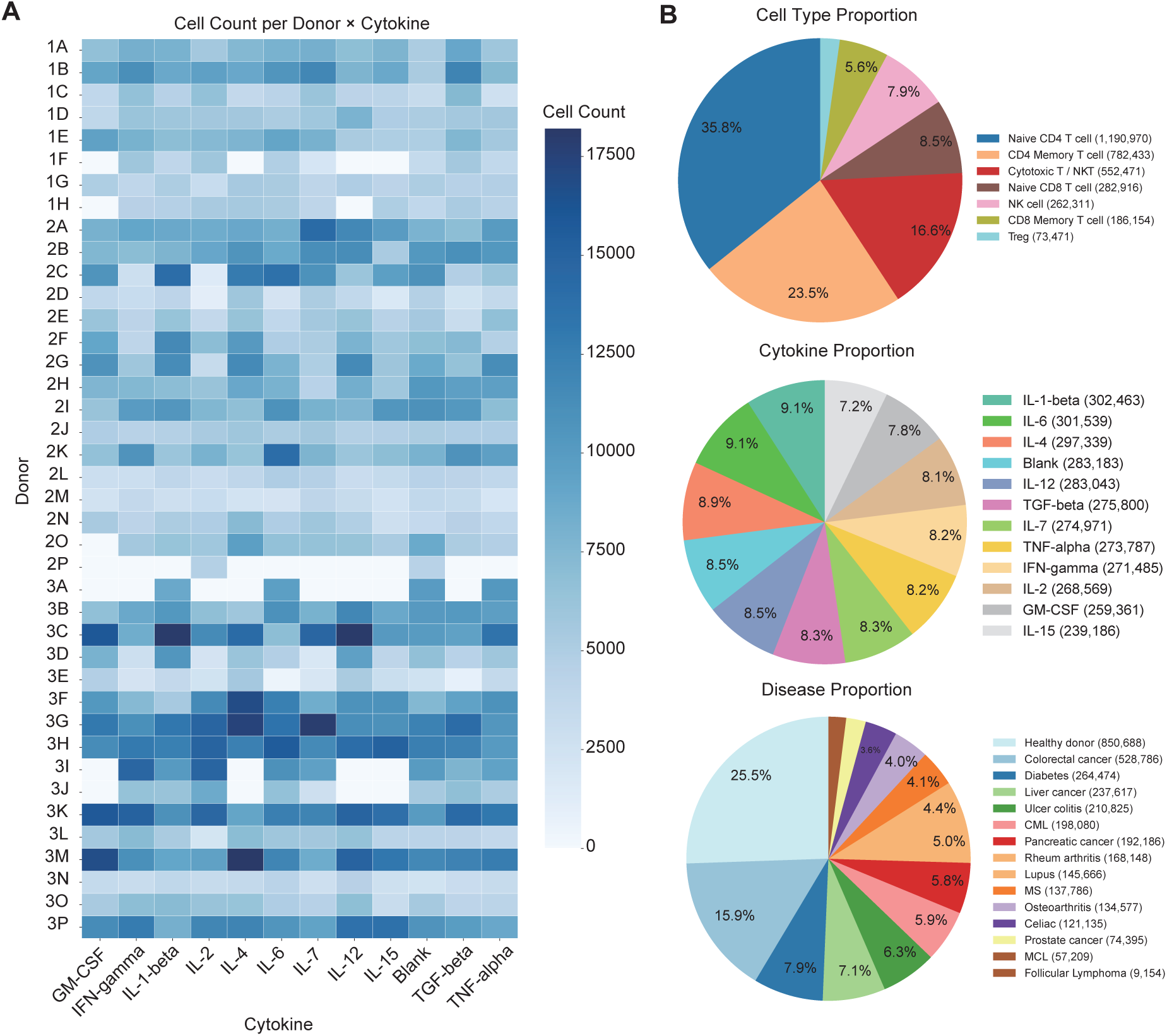
DiseasePert-3M data overview. **A.** Distribution of cell counts across donors and cytokine perturbation conditions. DiseasePert-3M comprises 447 donor–perturbation condition combinations with sufficient cells (>400). **B.** Composition of the dataset by cell type, cytokine condition, and disease status. For downstream analysis, we combined the CD8 memory cluster and the cytotoxic CD8 T/NKT cluster and labeled the merged population as effector CD8.

**Figure S23.**
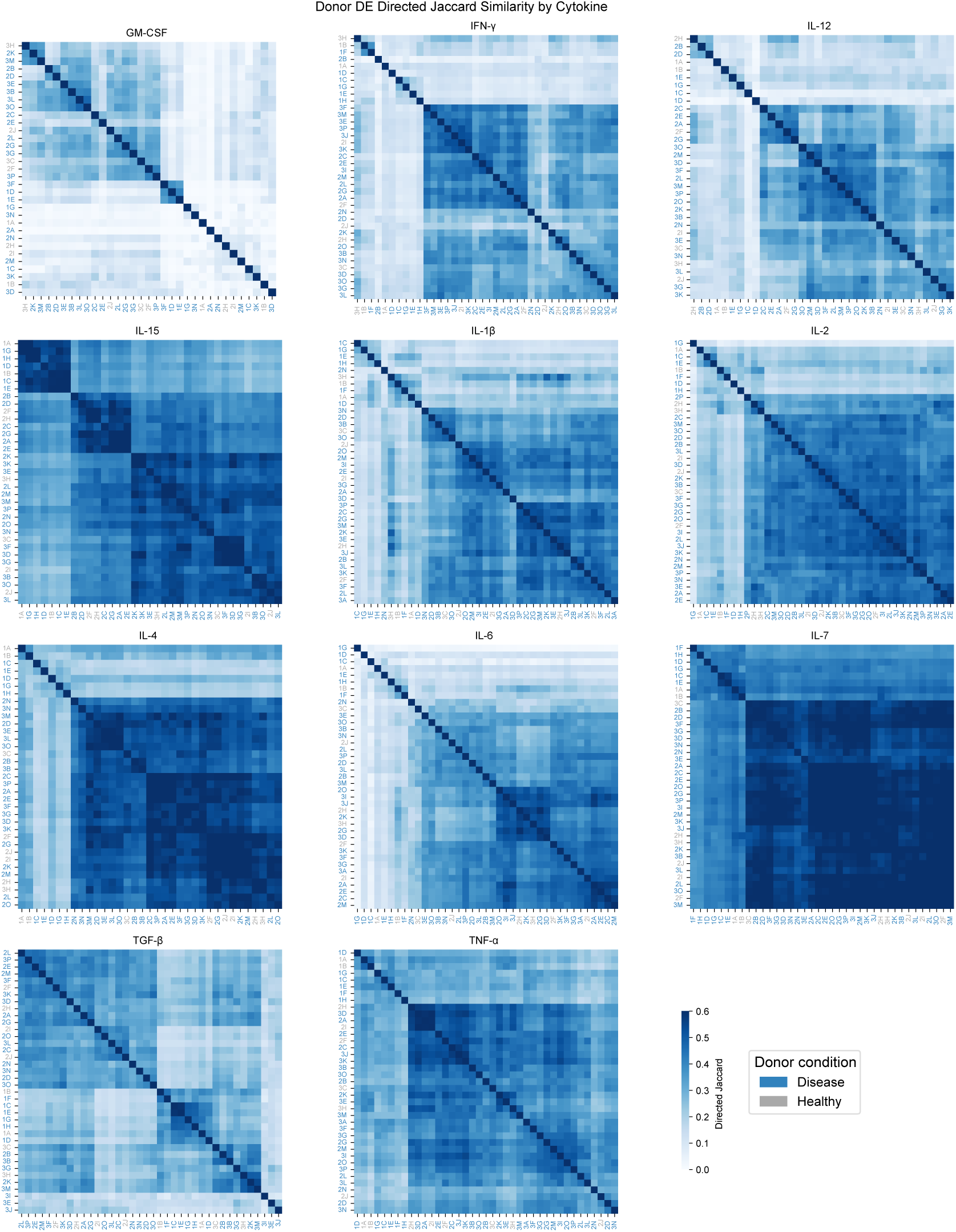
Pairwise directed Jaccard similarity of donor differential expression profiles by cytokine. Each heatmap shows the pairwise directed Jaccard similarity between donors for a given cytokine perturbation, averaged across evaluated T cell subtypes (Naive CD4, Memory CD4, Effector CD8).

**Figure S24.**
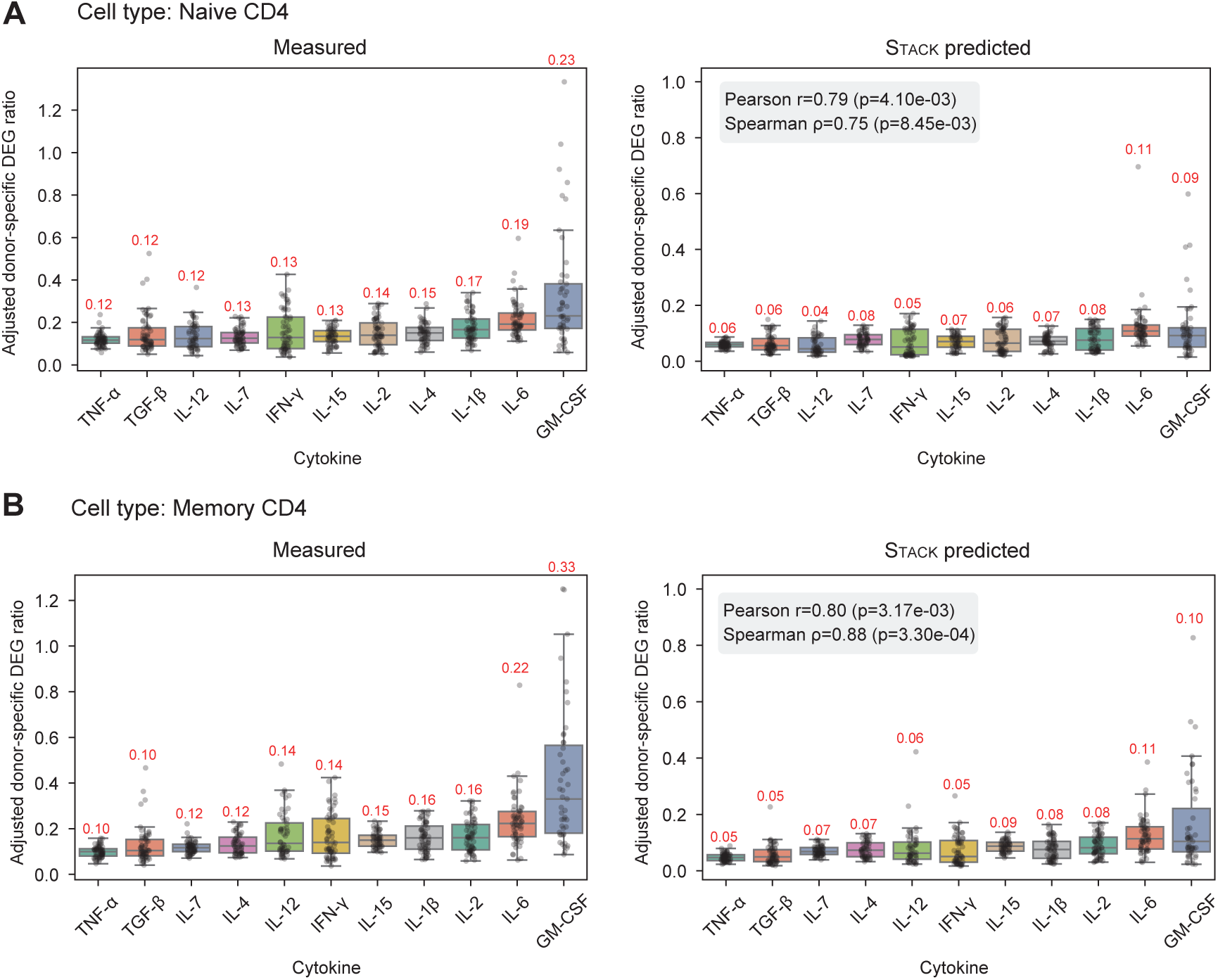
Additional evaluation of predicted donor-specific effect sizes across cytokine conditions. **A–B.** Box plots of adjusted donor-specific DEG ratios per cytokine perturbation for Naive CD4 (**A**) and Memory CD4 (**B**) T cells, comparing real data (left) and Stack predictions (right). Each point represents a prompt–query pair (*𝑛* = 638, 637 for **A**, **B**, respectively). Numbers above each box indicate the median adjusted ratio. Pearson and Spearman correlations between measured and Stack-predicted median adjusted donor-specific DEG ratios (*𝑟*, *𝜌*), along with their associated *𝑝*-values, are annotated in each panel.

**Figure S25.**
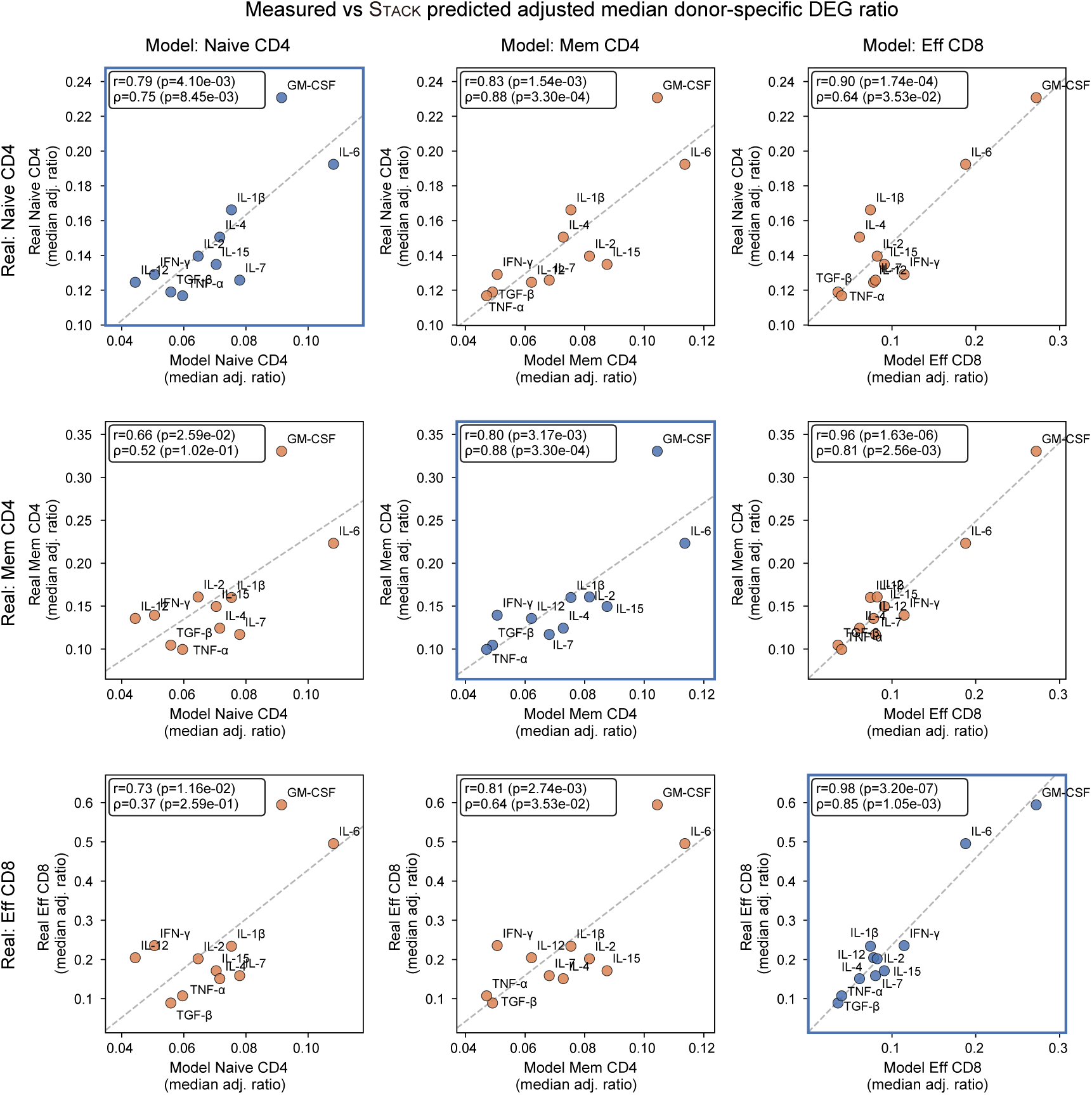
Cell-type specificity of adjusted donor-specific DEG ratios between Stack predictions and real measurements. Each point represents one cytokine (median adjusted ratio across donor pairs). Diagonal panels compare matched cell types; off-diagonal panels compare mismatched cell types. The dashed line indicates the linear regression fit. Pearson and Spearman correlations (*𝑟*, *𝜌*) with associated *𝑝*-values are annotated in each panel. All panels include *𝑛* = 11 cytokines.

**Figure S26.**
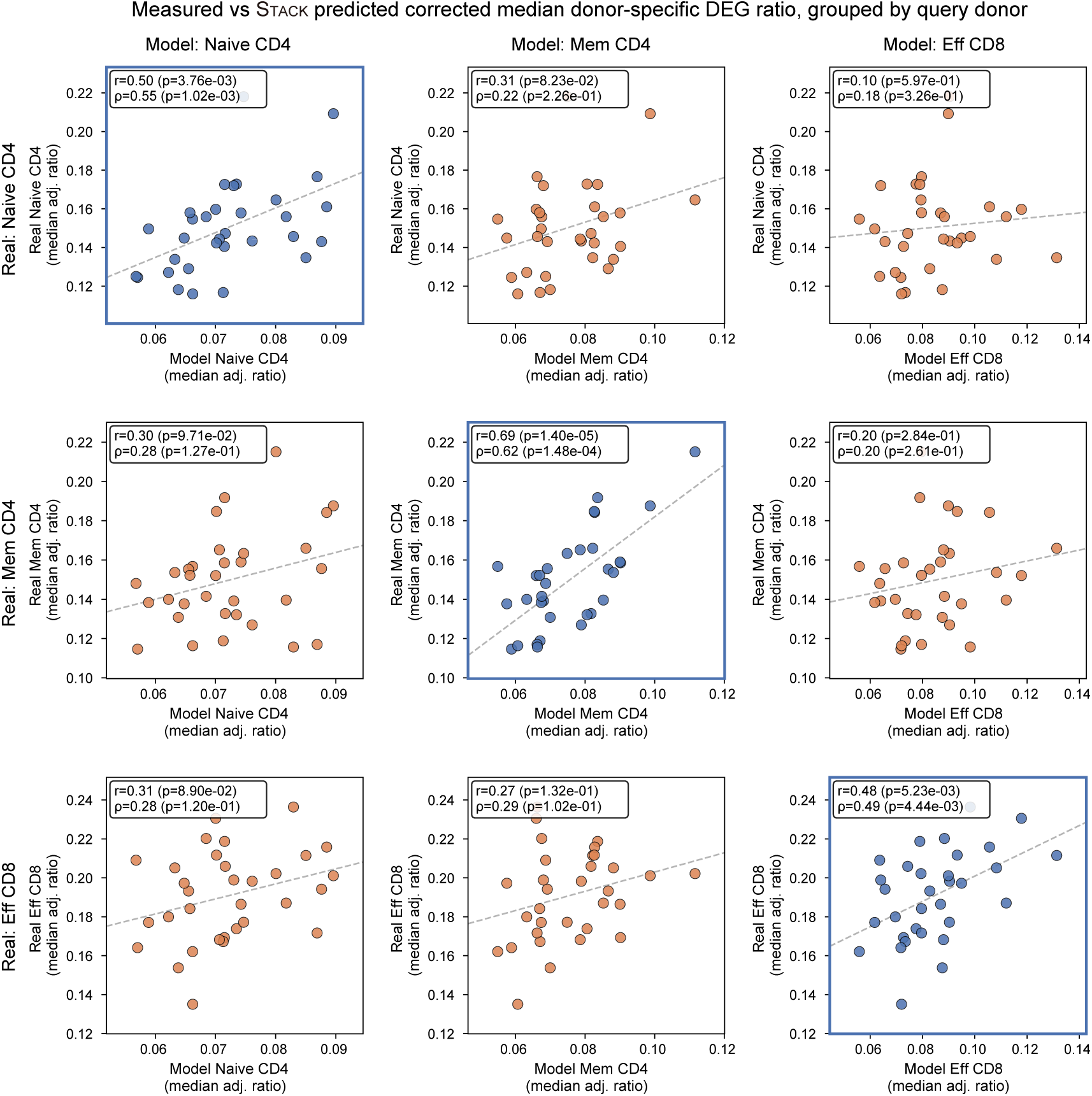
Cell-type specificity of adjusted donor-specific DEG ratios between Stack predictions and real measurements across disease queries. Each point represents one donor (median adjusted ratio across donor pairs). Diagonal panels compare matched cell types; off-diagonal panels compare mismatched cell types. The dashed line indicates the linear regression fit. Pearson and Spearman correlations (*𝑟*, *𝜌*) with associated *𝑝*-values are annotated in each panel. All panels include *𝑛* = 32 patients.

**Figure S27.**
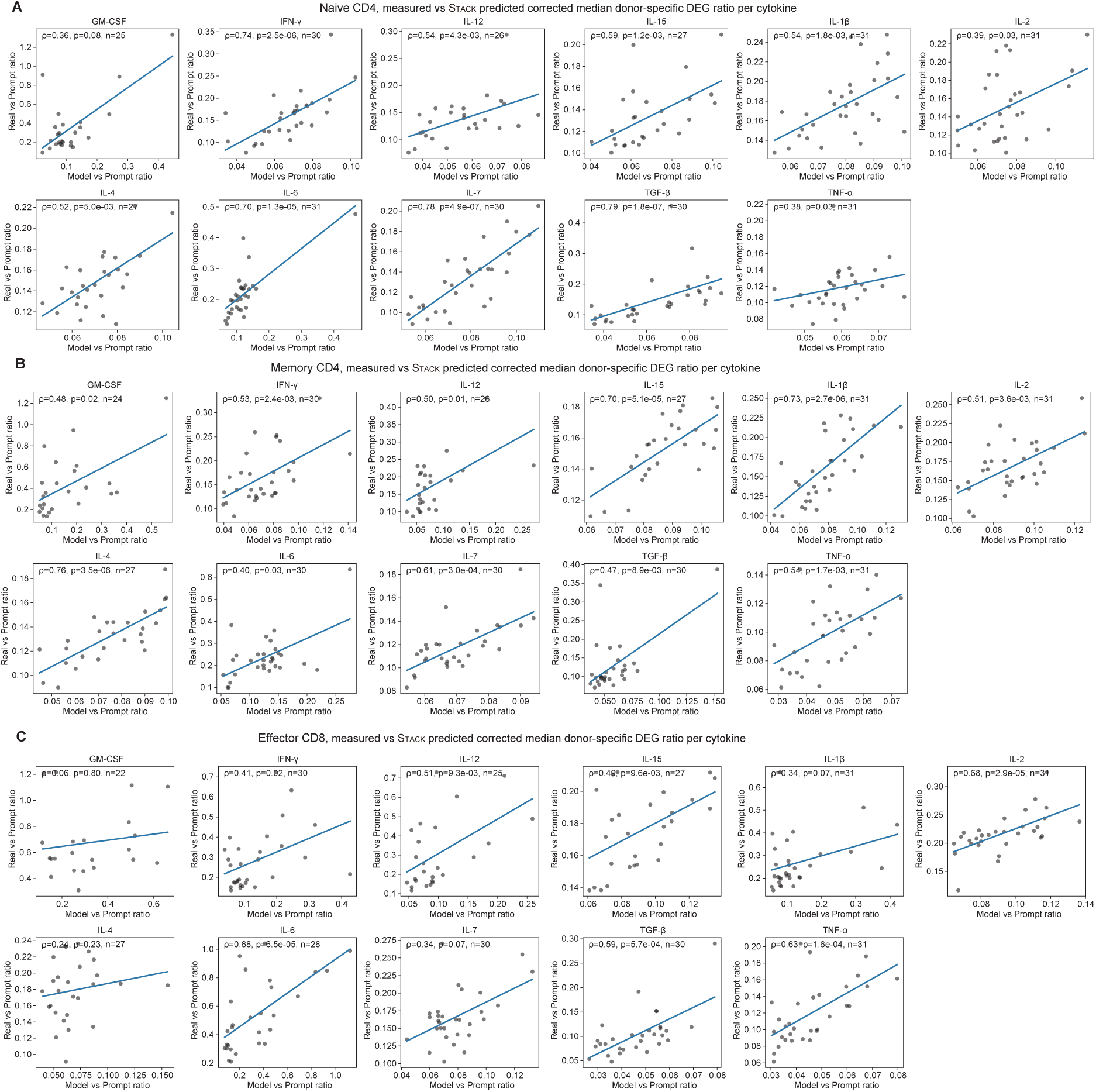
Evaluation of measured versus Stack predicted adjusted median donor-specific DEG ratio per cytokine. The blue line indicates the linear regression fit. Spearman correlation coefficients (*𝜌*) and associated *𝑝*-values are annotated in each panel. **A.** Naive CD4 T cells. **B.** Memory CD4 T cells. **C.** Effector CD8 T cells. The number of evaluated patients (*𝑛*) is indicated in each panel.

**Figure S28.**
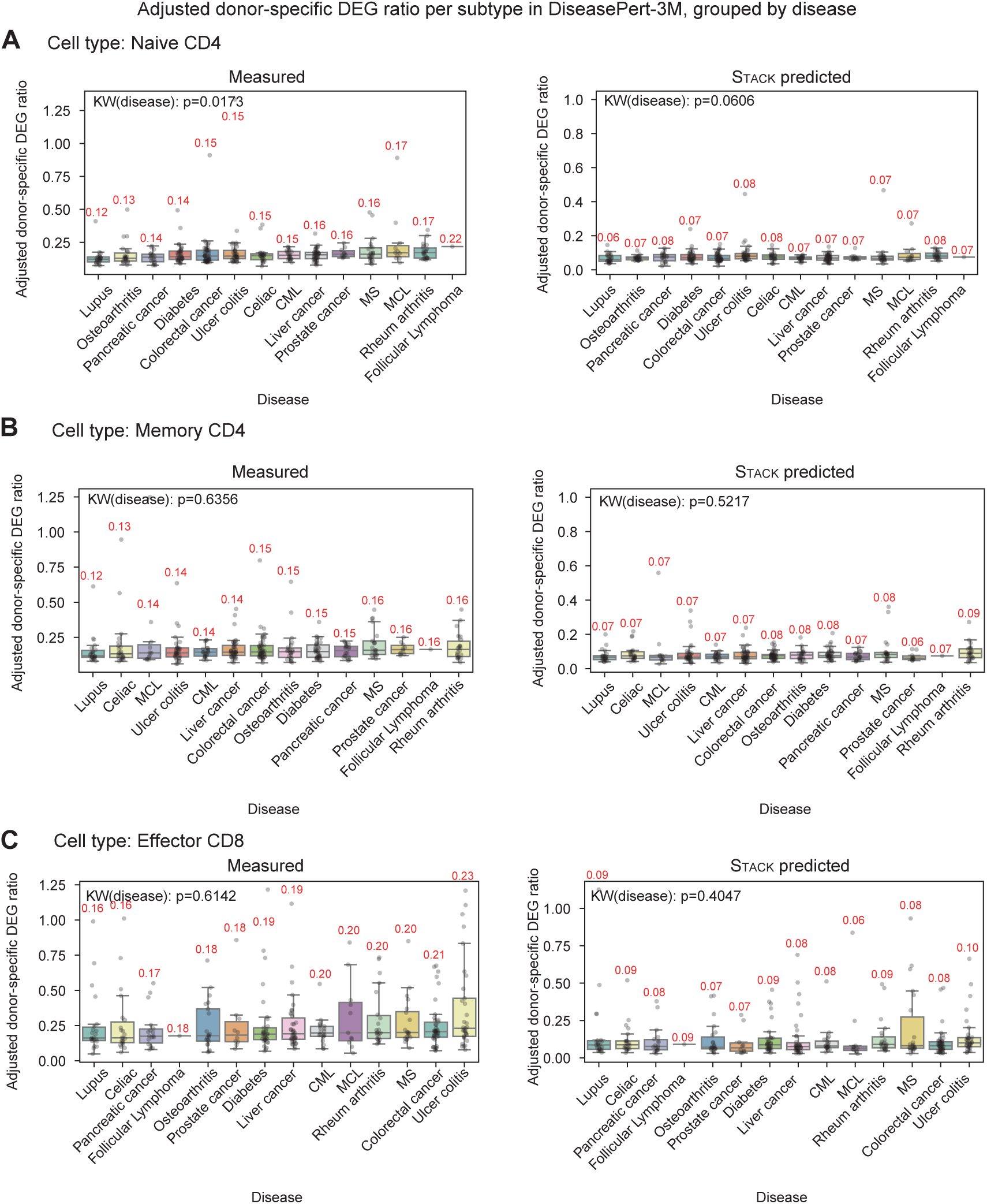
Additional evaluation of predicted donor-specific effect sizes across disease queries. A–C. Box plots of adjusted donor-specific DEG ratios per cytokine perturbation for Naive CD4 (**A**), Memory CD4 (B), and Effector CD8 (**C**) cell types, comparing real data (left) and Stack predictions (right), grouped by disease. Each point represents a query donor for a given cytokine, averaged per prompt (*𝑛* = 319, 318, 319 for **A**, **B**, **C**, respectively). Numbers above each box indicate the median adjusted ratio. Kruskal–Wallis tests were performed on the adjusted donor-specific DEG ratios.

**Figure S29.**
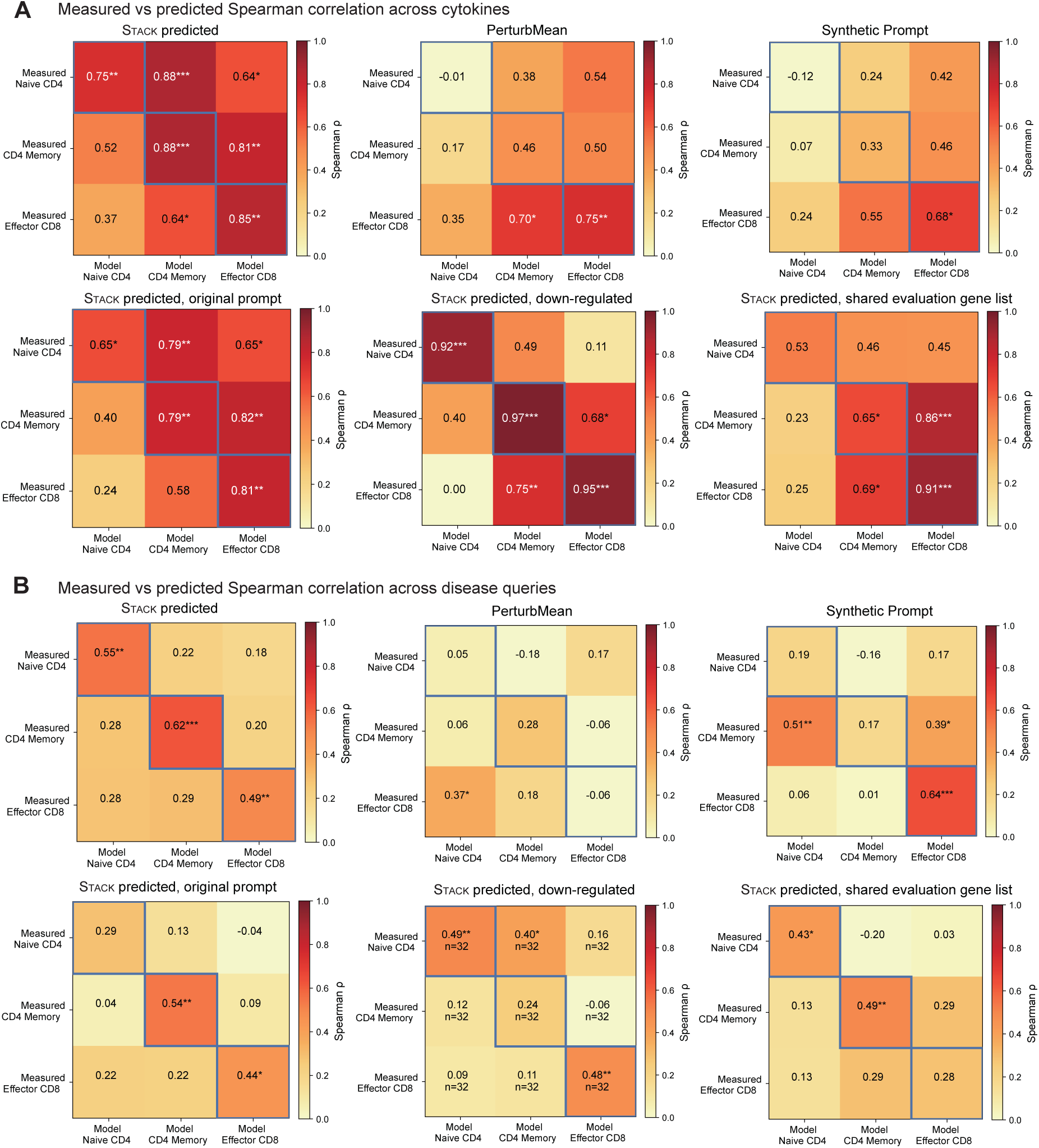
Additional evaluation of predicted donor-specific effect sizes across methods. **A.** Heatmap of Spearman correlation coefficients between measured and predicted adjusted median donor-specific DEG ratios across cytokines (*𝑛* = 11). Each entry is annotated with the correlation coefficient (*𝜌*) and corresponding significance level (^∗^ *𝑝 <* 0.05, ^∗∗^ *𝑝 <* 0.01, ^∗∗∗^ *𝑝 <* 0.001). **B.** Same as **A**, but computed across query donors (*𝑛* = 32).

**Figure S30.**
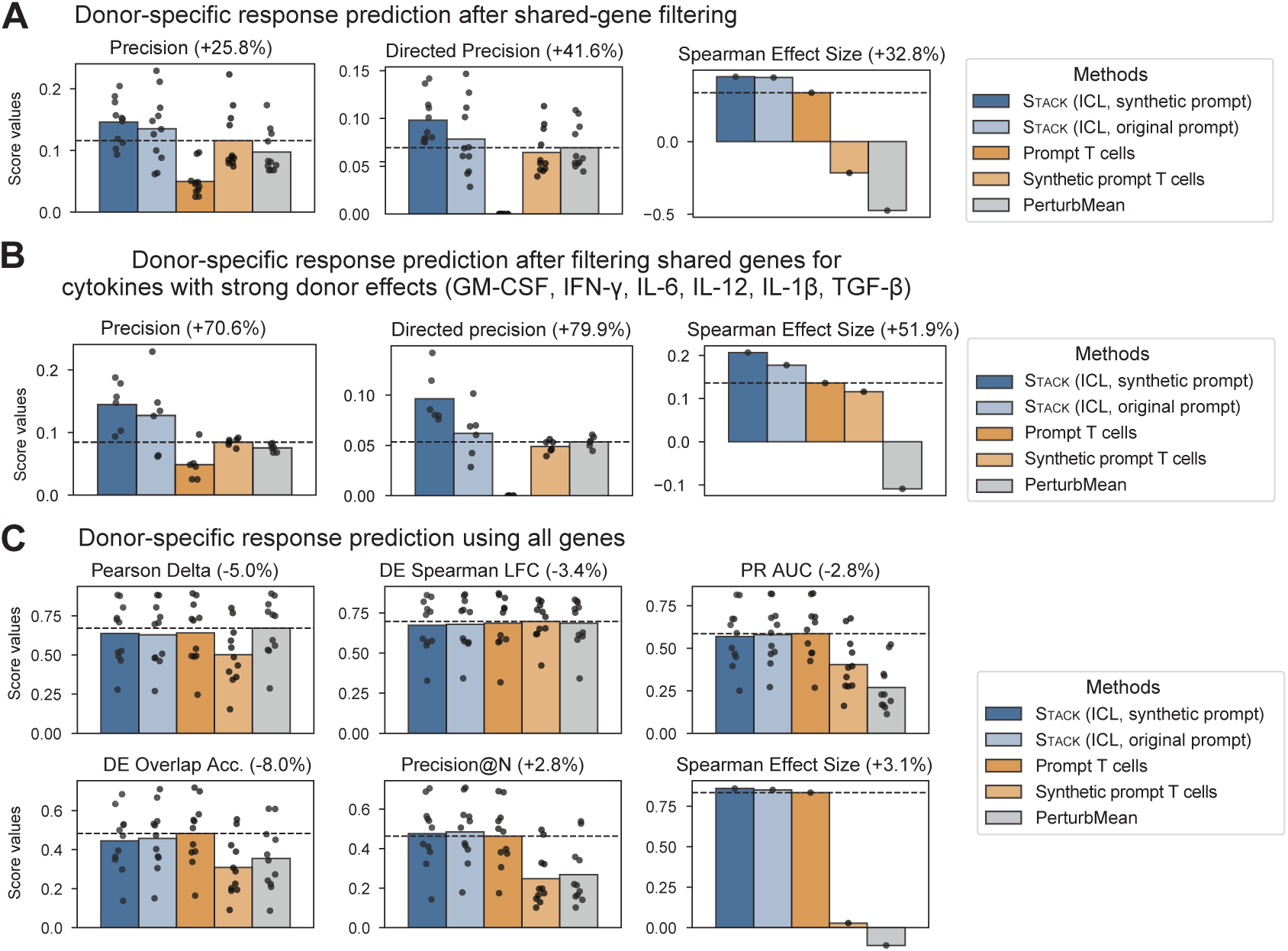
Evaluation of donor-specific T cell response prediction. **A.** Evaluation on non-shared genes per prompt-query pair (Methods). **B.** Evaluation on non-shared genes per prompt-query pair, focusing on 6 cytokines with more than 10 prompt-query donor pairs showing DE Direction Match < 0.75. Here, Spearman effect sizes are calculated from the uncorrected ratio of differentially expressed genes among all non-shared genes to match the cell-eval definition, which differ from the adjusted donor-specific DEG ratio. **C.** Evaluation on all genes. Percentages in titles represent the average improvement of Stack over the best non-Stack baseline. Dashed lines indicate the performance of the best competing method in each evaluation. In all panels except the Spearman effect size plot, each point represents the average result for one cytokine (*𝑛* = 11/6/11).

**Figure S31.**
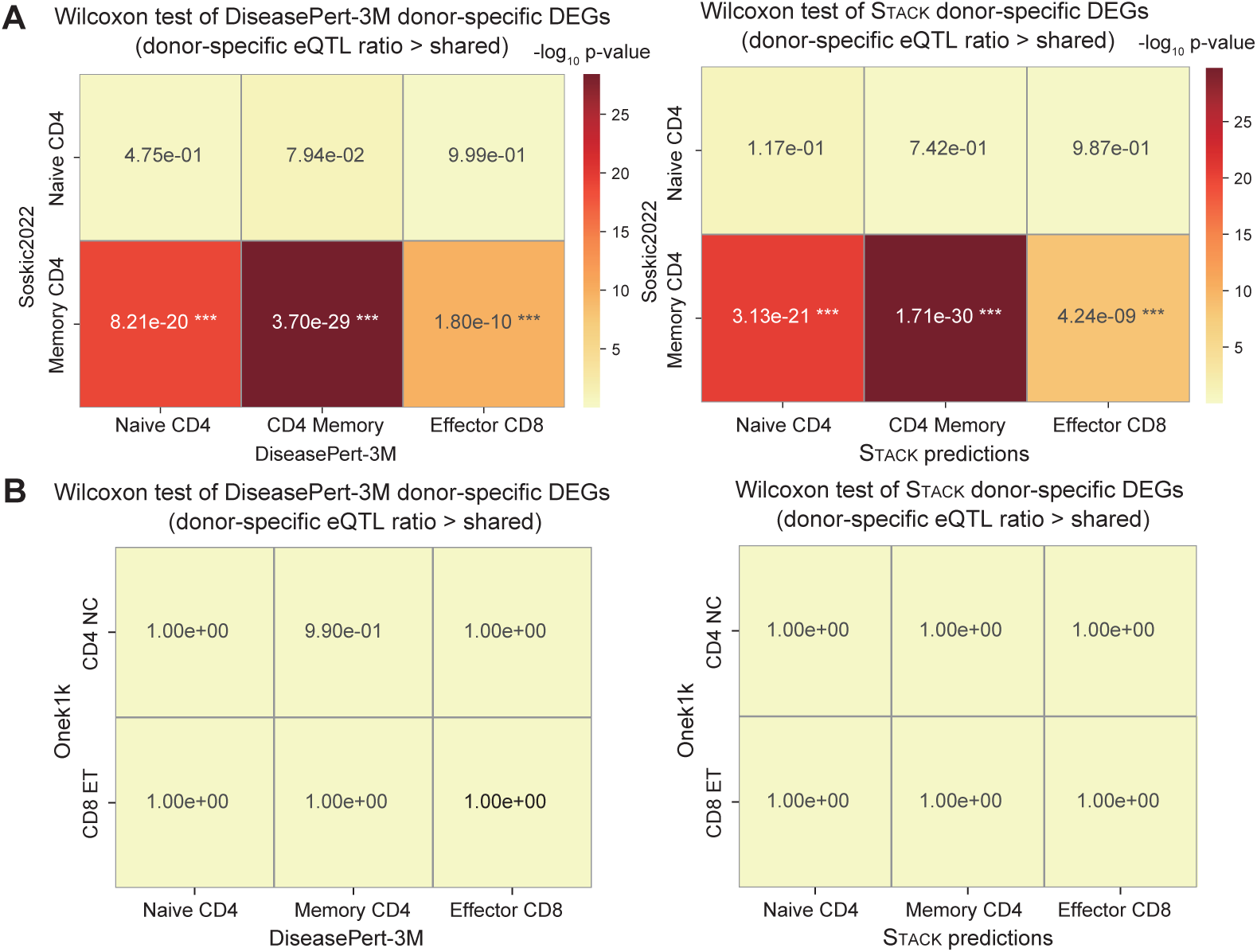
Additional evaluation of Stack donor-specific response prediction versus eQTL gene lists. **A.** Summary of (Soskic et al., 2022) eQTL enrichment significance across eQTL source and donor-specific DEGs per subtype in real data (left) and Stack predictions (right). significance levels are indicated by asterisks (^∗^ *𝑝 <* 0.05, ^∗∗^ *𝑝 <* 0.01, ^∗∗∗^ *𝑝 <* 0.001). **B.** Summary of OneK1K (Yazar et al., 2022) eQTL enrichment − log_10_ *𝑝*-value across eQTL source and donor-specific DEGs per subtype in real data (left) and Stack predictions (right). *𝑛* = 638, 636, 638 for the three DiseasePert-3M T cell subtypes respectively.

**Figure S32.**
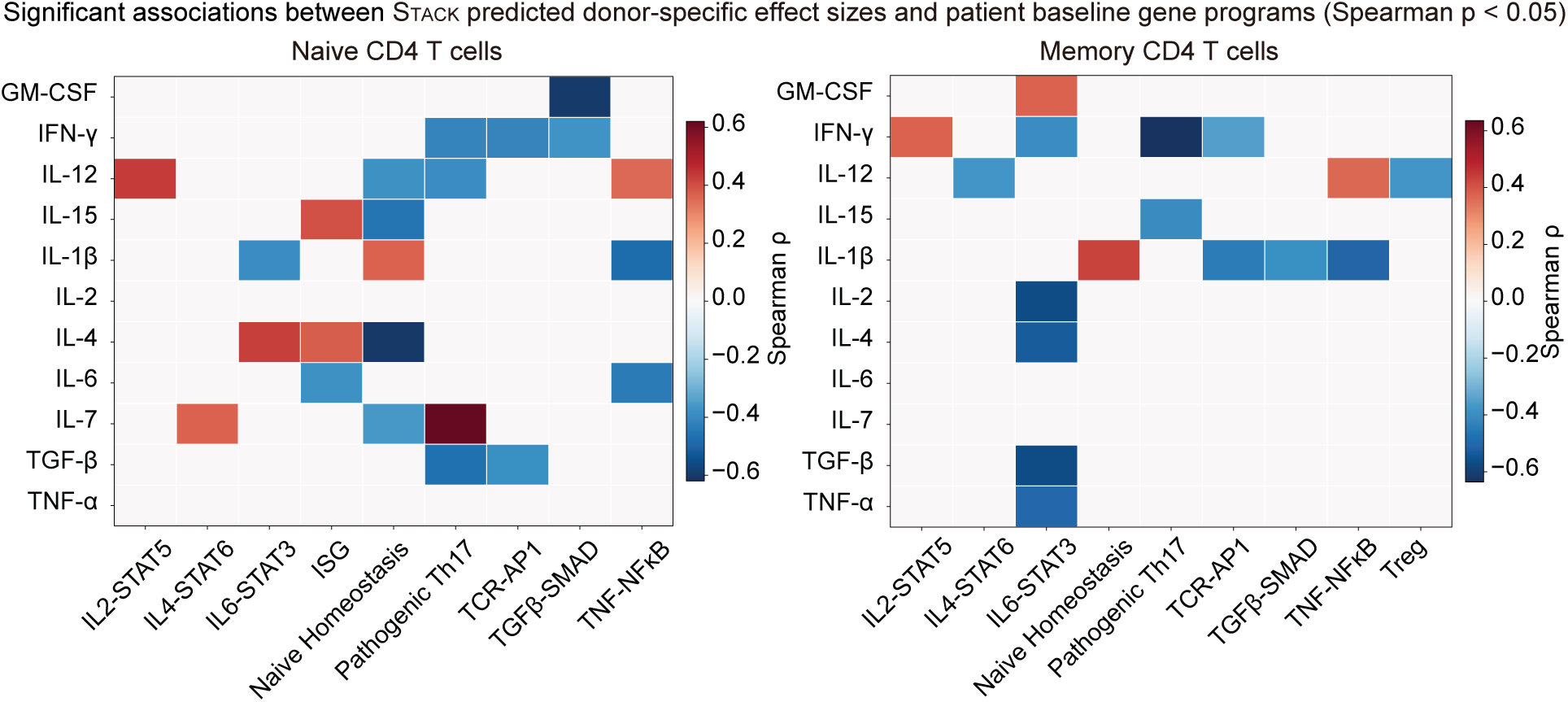
Heatmaps of Spearman correlation coefficients between Stack-predicted donor-specific effect sizes and patient baseline gene programs in naive CD4 T cells (left) and memory CD4 T cells (right). Only significant Spearman *𝜌* values (*𝑝 <* 0.05) are shown in color.

**Figure S33.**
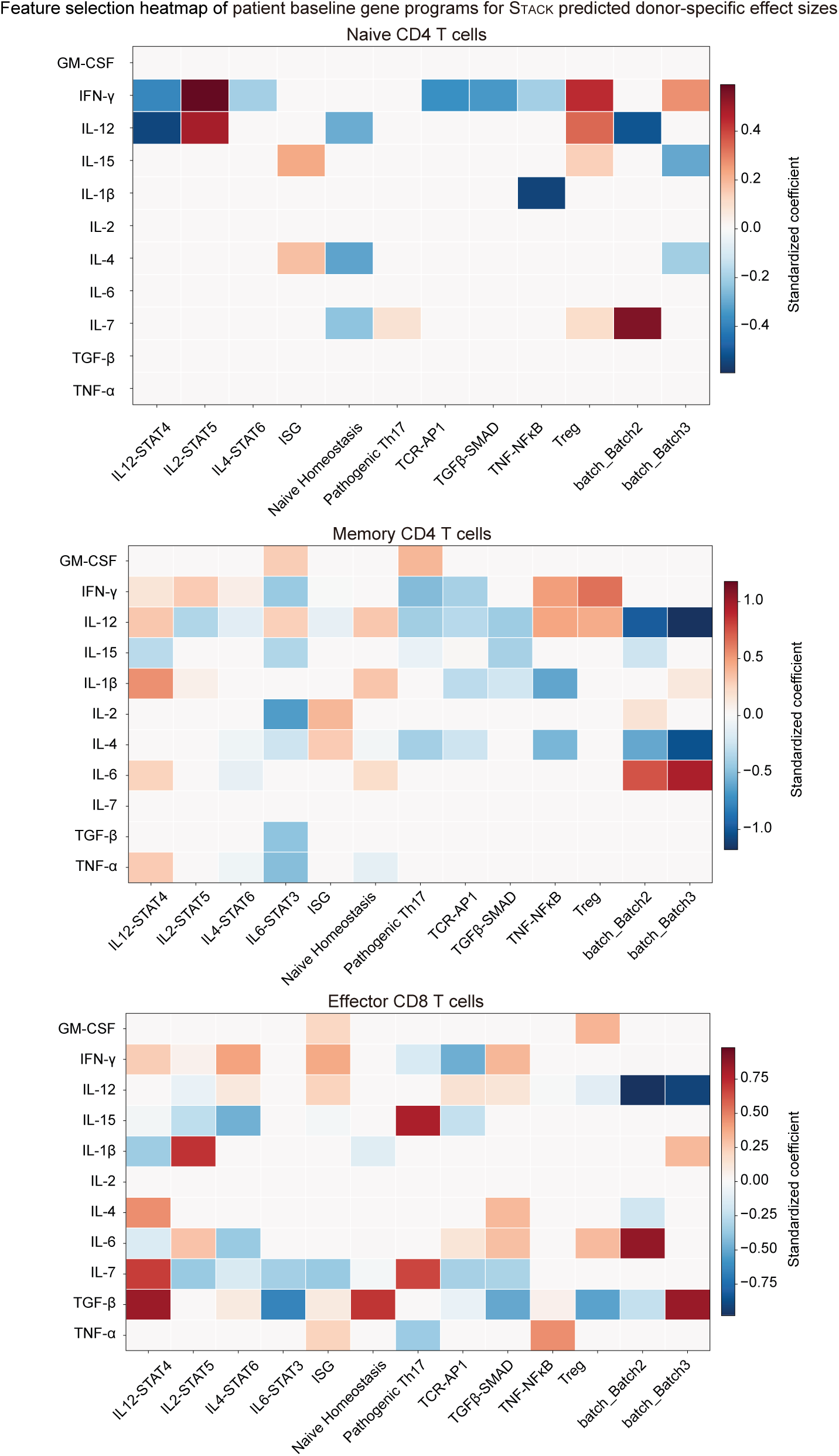
Heatmaps showing standardized linear regression coefficients for gene modules selected by leave-one-out cross-validated elastic net models to predict Stack donor-specific effect sizes (Methods).

**Table S1.**
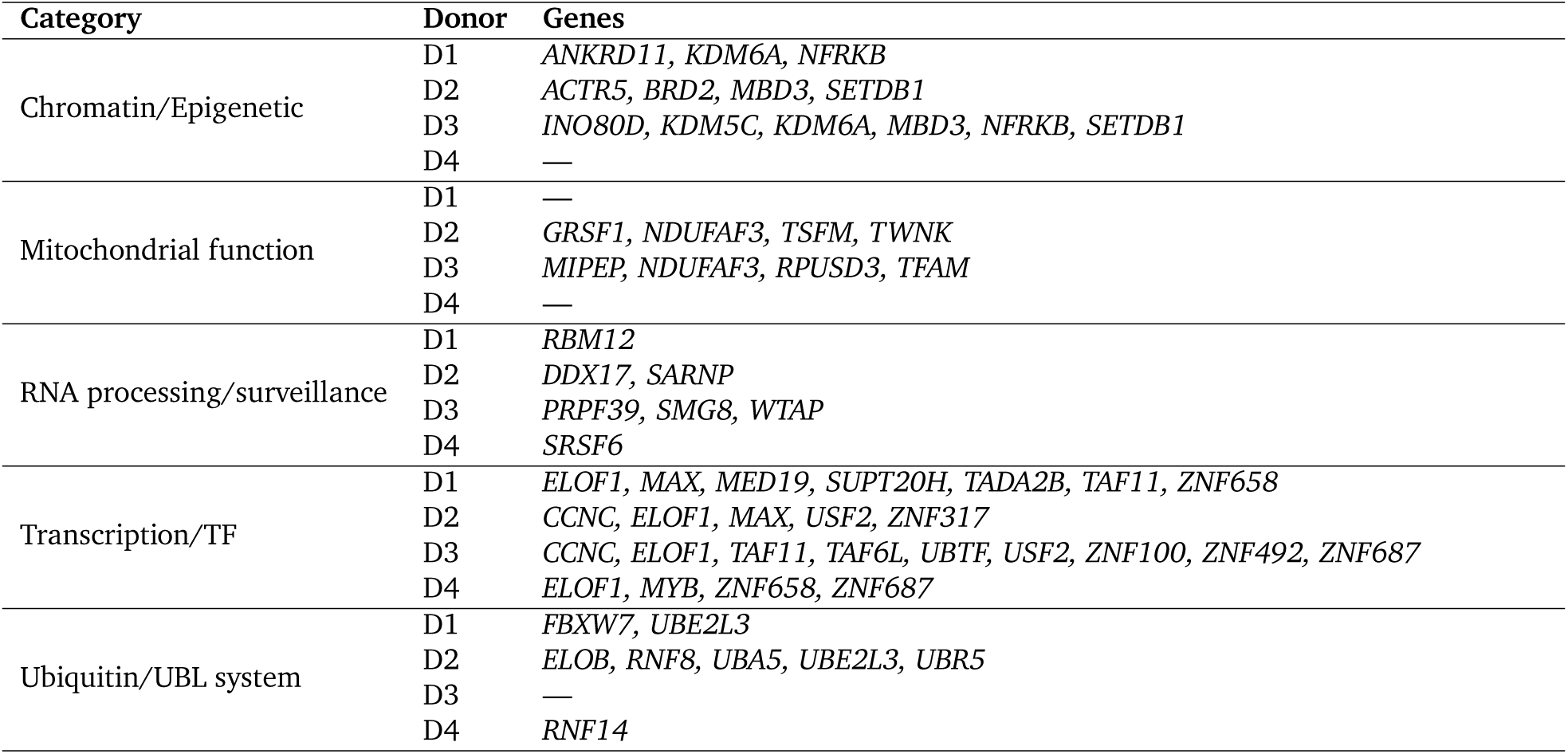
Shared genetic perturbations between the RPE1 dataset (Replogle et al., 2022) and *Perturb Sapiens* within gene functionalities with total gene number > 5, as shown in Fig. 5E.

**Table S2.**
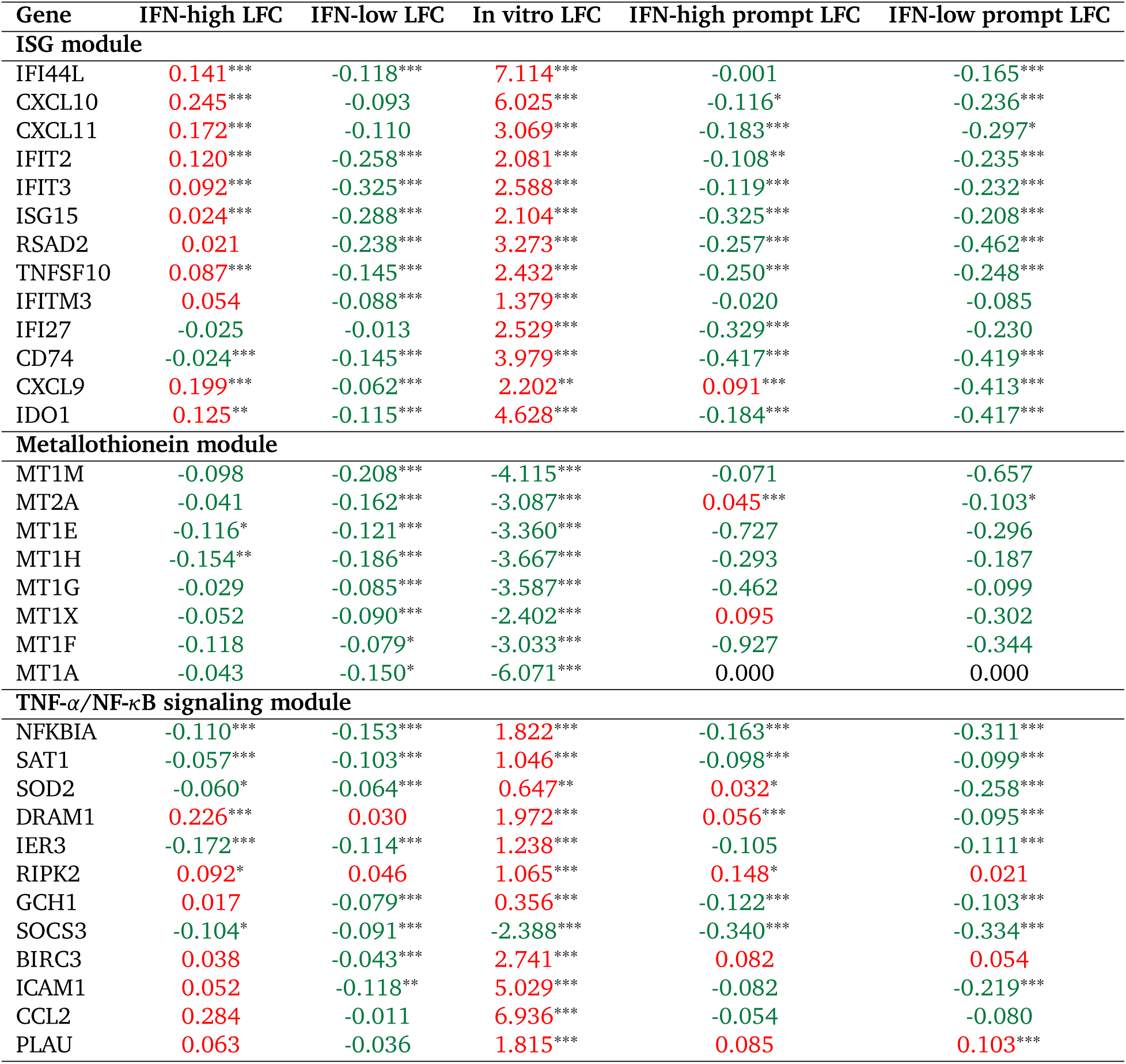
Response comparison across IFN-high *Perturb Sapiens*, IFN-low *Perturb Sapiens*, *in vitro* TNF-*𝛼* data (Lee et al., 2022), and IFN-high/low prompt myeloid cells. LFC values are colored red (upregulated) or green (downregulated); significance levels: ^∗∗∗^*𝑞 <* 0.001, ^∗∗^*𝑞 <* 0.01, ^∗^*𝑞 <* 0.05.

| Gene | IFN-high LFC | IFN-low LFC | In vitro LFC | IFN-high prompt LFC | IFN-low prompt LFC |
| --- | --- | --- | --- | --- | --- |
| **ISG module** |  |  |  |  |  |
| IFI44L | 0.141*** | -0.118*** | 7.114*** | -0.001 | -0.165*** |
| CXCL10 | 0.245*** | -0.093 | 6.025*** | -0.116* | -0.236*** |
| CXCL11 | 0.172*** | -0.110 | 3.069*** | -0.183*** | -0.297* |
| IFIT2 | 0.120*** | -0.258*** | 2.081*** | -0.108** | -0.235*** |
| IFIT3 | 0.092*** | -0.325*** | 2.588*** | -0.119*** | -0.232*** |
| ISG15 | 0.024*** | -0.288*** | 2.104*** | -0.325*** | -0.208*** |
| RSAD2 | 0.021 | -0.238*** | 3.273*** | -0.257*** | -0.462*** |
| TNFSF10 | 0.087*** | -0.145*** | 2.432*** | -0.250*** | -0.248*** |
| IFITM3 | 0.054 | -0.088*** | 1.379*** | -0.020 | -0.085 |
| IFI27 | -0.025 | -0.013 | 2.529*** | -0.329*** | -0.230 |
| CD74 | -0.024*** | -0.145*** | 3.979*** | -0.417*** | -0.419*** |
| CXCL9 | 0.199*** | -0.062*** | 2.202** | 0.091*** | -0.413*** |
| IDO1 | 0.125** | -0.115*** | 4.628*** | -0.184*** | -0.417*** |
| **Metallothionein module** |  |  |  |  |  |
| MT1M | -0.098 | -0.208*** | -4.115*** | -0.071 | -0.657 |
| MT2A | -0.041 | -0.162*** | -3.087*** | 0.045*** | -0.103* |
| MT1E | -0.116* | -0.121*** | -3.360*** | -0.727 | -0.296 |
| MT1H | -0.154** | -0.186*** | -3.667*** | -0.293 | -0.187 |
| MT1G | -0.029 | -0.085*** | -3.587*** | -0.462 | -0.099 |
| MT1X | -0.052 | -0.090*** | -2.402*** | 0.095 | -0.302 |
| MT1F | -0.118 | -0.079* | -3.033*** | -0.927 | -0.344 |
| MT1A | -0.043 | -0.150* | -6.071*** | 0.000 | 0.000 |
| **TNF-α/NF-κB signaling module** |  |  |  |  |  |
| NFKBIA | -0.110*** | -0.153*** | 1.822*** | -0.163*** | -0.311*** |
| SAT1 | -0.057*** | -0.103*** | 1.046*** | -0.098*** | -0.099*** |
| SOD2 | -0.060* | -0.064*** | 0.647** | 0.032* | -0.258*** |
| DRAM1 | 0.226*** | 0.030 | 1.972*** | 0.056*** | -0.095*** |
| IER3 | -0.172*** | -0.114*** | 1.238*** | -0.105 | -0.111*** |
| RIPK2 | 0.092* | 0.046 | 1.065*** | 0.148* | 0.021 |
| GCH1 | 0.017 | -0.079*** | 0.356*** | -0.122*** | -0.103*** |
| SOCS3 | -0.104* | -0.091*** | -2.388*** | -0.340*** | -0.334*** |
| BIRC3 | 0.038 | -0.043*** | 2.741*** | 0.082 | 0.054 |
| ICAM1 | 0.052 | -0.118** | 5.029*** | -0.082 | -0.219*** |
| CCL2 | 0.284 | -0.011 | 6.936*** | -0.054 | -0.080 |
| PLAU | 0.063 | -0.036 | 1.815*** | 0.085 | 0.103*** |

**Table S3.** Evaluation of *Perturb Sapiens* (IFN-high/low donor groups) intestinal epithelial cells using bulk IL-1*𝛽* stimulation data from primary keratinocytes (Swindell et al., 2018). Significance: *** *𝑝 <* 0.001

|  | Pearson Delta | DE Spearman LFC | DE Direction Match | PR-AUC | DE Overlap Acc. |
| --- | --- | --- | --- | --- | --- |
| IFN-high | 0.141*** | 0.288*** | 0.875 | 0.185 | 0.148 |
| IFN-low | 0.213*** | 0.359*** | 0.707 | 0.183 | 0.163 |

**Table S4.** DiseasePert-3M donor disease information.

| Donor | Disease |
| --- | --- |
| 1A | Healthy donor |
| 1B | Healthy donor |
| 1C | Rheum arthritis |
| 1D | Osteoarthritis |
| 1E | Celiac |
| 1F | Prostate cancer |
| 1G | Liver cancer |
| 1H | CML |
| 2A | Colorectal cancer |
| 2B | Rheum arthritis |
| 2C | Pancreatic cancer |
| 2D | Diabetes |
| 2E | Liver cancer |
| 2F | Healthy donor |
| 2G | Lupus |
| 2H | Healthy donor |
| 2I | Healthy donor |
| 2J | Healthy donor |
| 2K | MS |
| 2L | Lupus |
| 2M | Celiac |
| 2N | MCL |
| 2O | Liver cancer |
| 2P | Follicular Lymphoma |
| 3A | Prostate cancer |
| 3B | Ulcer colitis |
| 3C | Healthy donor |
| 3D | Osteoarthritis |
| 3E | MS |
| 3F | Colorectal cancer |
| 3G | Colorectal cancer |
| 3H | Healthy donor |
| 3I | Pancreatic cancer |
| 3J | Liver cancer |
| 3K | Diabetes |
| 3L | Ulcer colitis |
| 3M | CML |
| 3N | Ulcer colitis |
| 3O | Diabetes |
| 3P | Colorectal cancer |

**Table S5.**
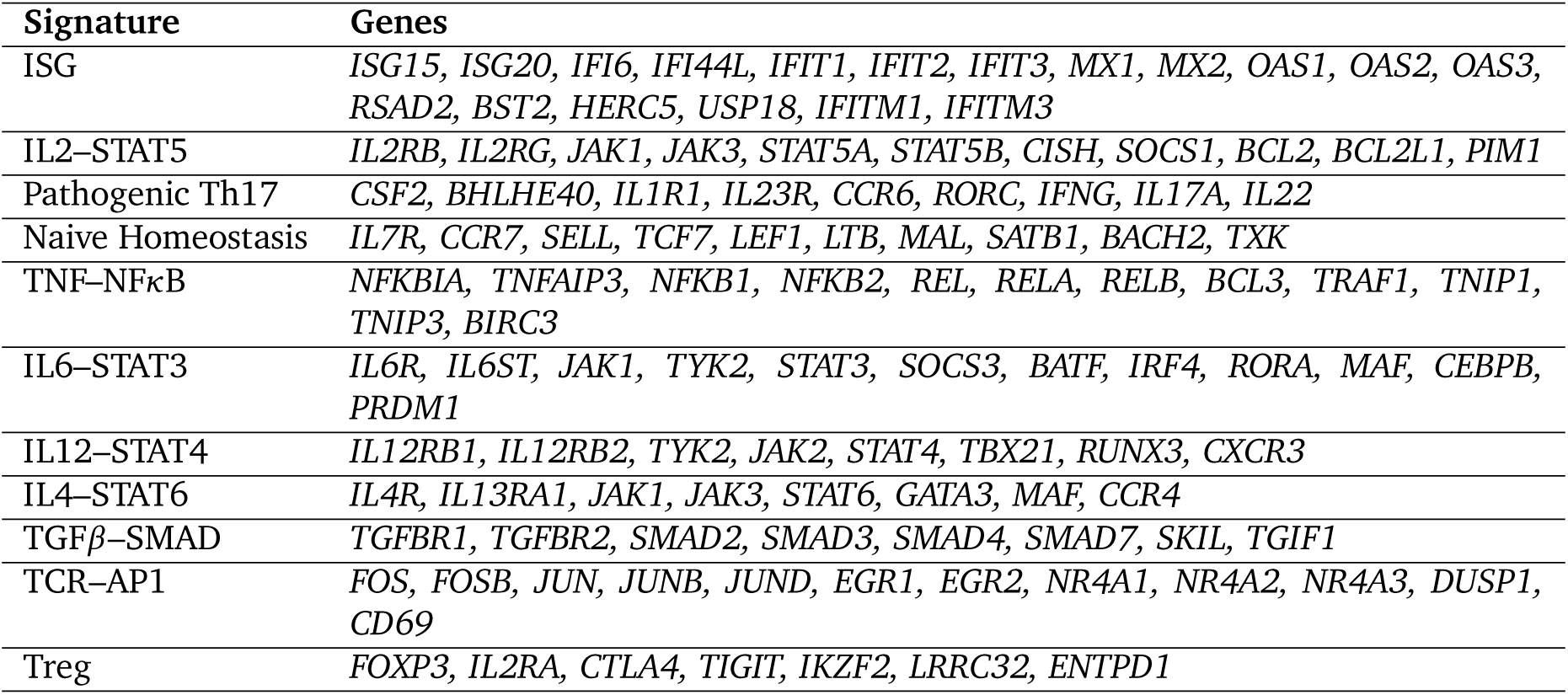
Gene signatures used for gene module scoring.

**Table S6.** Alignment between measured and Stack>-predicted cytokine–gene module Spearman correlations across T cell subsets under two evaluation settings. *Model 𝑝 <* 0.05 requires a significant Spearman correlation in the Stack>-predicted results only, whereas *Both 𝑝 <* 0.05 requires significance in both measured and Stack>-predicted results.

| Cell Type | Setting | #Pairs | Pearson $r$ | Spearman $\rho$ | Dir. Match |
| --- | --- | --- | --- | --- | --- |
| Naive CD4 | Model $p < 0.05$ | 23 | 0.892 | 0.707 | 1.000 |
| | Both $p < 0.05$ | 9 | 0.973 | 0.817 | 1.000 |
| Memory CD4 | Model $p < 0.05$ | 17 | 0.848 | 0.821 | 0.941 |
| | Both $p < 0.05$ | 7 | 0.989 | 0.500 | 1.000 |
| Effector CD8 | Model $p < 0.05$ | 28 | 0.862 | 0.703 | 0.893 |
| | Both $p < 0.05$ | 11 | 0.980 | 0.791 | 1.000 |

**Table S7.** Cytokine panel and working concentrations used for T cell stimulation. *10 ng/mL in batch 1; 12.5 ng/mL in batches 2 and 3.

| Cytokine | Functional Category | Working Conc. |
| --- | --- | --- |
| TNF-α | Pro-inflammatory | 100 ng/mL |
| IL-1β | Pro-inflammatory | 50 ng/mL |
| IL-6 | Pro-inflammatory | 60 ng/mL |
| TGF-β | Anti-inflammatory | 50 ng/mL |
| IFN-γ | Type II Interferon | 200 ng/mL |
| IL-12 | Th1 | 75 ng/mL |
| IL-4 | Th2 | 60 ng/mL |
| IL-2 | γc chain | 50 U/mL |
| IL-7 | γc chain | 10–12.5 ng/mL * |
| IL-15 | γc chain | 10–12.5 ng/mL * |
| GM-CSF | Growth factor | 50 ng/mL |

